# *Arabidopsis* inositol polyphosphate kinase activities regulate COP9 deneddylation functions in phosphate-homeostasis

**DOI:** 10.1101/2020.10.02.323584

**Authors:** Yashika Walia, Medha Noopur, Bhaskar Chandra Sahoo, Ishana Bhattacharya, Mritunjay Kasera, Naga Jyothi Pullagurla, Shobhita Saxena, Guizhen Liu, Gabriel Schaaf, Henning Jessen, Debabrata Laha, Souvik Bhattacharjee, Saikat Bhattacharjee

## Abstract

Plant Cullin RING Ubiquitin E3 ligases (CRLs) facilitate targeted protein degradation during physiological development and adaptation to stress. The deneddylase activity of COP9 signalosome (CSN) regulates cellular ratios of neddylated cullins available for the continuum of CRL functions. While selective inositol polyphosphates (InsPs) function as co-factors in plant responses involving the ubiquitylation of negative regulators, a relationship to CSN-CRL activities has not yet been established. Here, we show that the two *Arabidopsis thaliana* InsP-kinases IPK1 and ITPK1 physically interact and metabolically connect with the CSN holo-complex to modulate cullin deneddylation efficiency. Specifically, functional deficiency of ITPK1 lowers cullin deneddylation rates and disrupts the dissociation equilibrium of CSN5, the deneddylase catalytic subunit, and CUL1 with the holo-complex. Our results identify a novel auto-regulatory switch of CSN functions, defined by deneddylation activity. We further demonstrate that phosphate starvation response (PSR), which is induced in Pi-starved wild-type plants and constitutive in the above InsP-kinase mutants, is orchestrated in part by reduced deneddylation rates that, in turn, affect the stability of SPX4, a key negative regulator of PSR. Pharmacological inhibition of cullin neddylation stabilizes SPX4 and impairs PSR, thus linking CSN-CRL dynamics to phosphate (Pi)-sensing. Conversely, when exposed to compounds that inhibit CSN5 deneddylase activity, wild-type plants display phenotypes similar to the above InsP-kinase mutants. Overall, our data reveal that the regulation of plant Pi-starvation responses by specific InsP-kinases is caused by a direct role of these kinases in balancing coordination between CRL-CSN activities.

---

The versatile functions of soluble inositol polyphosphates (InsPs) in cellular processes in eukaryotes result from their propensity for being differentially phosphorylated by a relatively largely conserved set of InsP-kinases (Kim et al., 2024; Laha et al., 2021; Monserrate and York, 2010; Shears and Wang, 2019; Wilson et al., 2013). In plants, particular InsPs have been linked to phytohormone perception, inorganic phosphate (Pi)-homeostasis, mRNA export, meristem maintenance, and plant immunity (Alcazar-Roman et al., 2010; Dong et al., 2019; Gillaspy, 2013; Gulabani et al., 2022; Jia et al., 2019; Kuo et al., 2014; Kuo et al., 2018; Laha et al., 2015; Laha et al., 2016; Laha et al., 2019; Laha et al., 2021; Laha et al., 2022; Murphy et al., 2008; Poon et al., 2020; Pullagurla et al., 2024; Ren et al., 2024; Riemer et al., 2021; Stevenson-Paulik et al., 2005; Wild et al., 2016; Williams et al., 2015; Zhu et al., 2009). The synthesis of the fully phosphorylated *myo*-inositol derivative, InsP_6_ (*myo*-inositol-1,2,3,4,5,6-hexakisphosphate, also known as phytic acid), from InsP_5_ [2OH] is catalyzed by INOSITOL PENTAKISPHSPHATE 2-KINASE 1 (IPK1) and is sequentially (pyro)-phosphorylated by INOSITOL 1,3,4-TRISPHOSPHATE 5/6-KINASES (ITPK1/2) and then by DIPHOSPHOINOSITOL PENTAKISPHOSPHATE KINASES (VIH1/2) to generate the energy-rich phosphoanhydride bond-containing inositol pyrophosphates, including various InsP_7_ isomers and InsP_8_ (Adepoju et al., 2019; Desai et al., 2014; Laha et al., 2015; Laha et al., 2019; Laha et al., 2022; Pullagurla et al., 2024; Riemer et al., 2021; Stevenson-Paulik et al., 2005; Whitfield et al., 2020; Zhu et al., 2019). Recent studies have indicated (or have proposed) that InsP_6_, and more recently 5-InsP_7_, promote auxin-receptor complex formation that may eventually drive the degradation of AUX/IAA repressors, whereas InsP_8_ may similarly facilitate targeting of JASMONATE ZIM-DOMAIN (JAZ) repressors (Laha et al., 2015; Laha et al., 2022; Pullagurla et al., 2023; Sheard et al., 2010; Tan et al., 2007). Interestingly, the cognate substrate receptors TRANSPORT INHIBITOR RESPONSE 1 (TIR1) and CORONATINE INSENSITIVE 1 (COI1) belong to the F-box family proteins and are components of typical SCF (ASK1/SKP1-Cullin-F-Box; SCF^TIR1^ and SCF^COI1^)-type E3 ubiquitin ligases (Gray et al., 1999; Gray et al., 2001; Sheard et al., 2010; Tan et al., 2007; Williams et al., 2019; Xu et al., 2002).

InsPs have also been recently implicated as sensors of phosphate (Pi)-status in plants (Cridland and Gillaspy, 2020; Dong et al., 2019; Guan et al., 2022; Kuo et al., 2018; Pullagurla et al., 2024; Ried et al., 2021; Riemer et al., 2021; Stevenson-Paulik et al., 2005; Wild et al., 2016; Zhu et al., 2019). Under low Pi-availability from the soil, a highly organized adaptive cascade of events collectively termed phosphate starvation response (PSR) is activated. Such events include the upregulated expression of phosphate-starvation inducible (*PSI*)-genes such as for Pi-transporters (*PHT*s), phosphatases (*PAP*s), and several micro-(*MIR399*) or non-coding RNAs (*IPS1*), that results in increased Pi mobilization in the rhizosphere/soil, as well as in increased Pi uptake and remobilization from source tissues and intracellular Pi-storage reserves (Ham et al., 2018; Kuo et al., 2014; Puga et al., 2014). In Arabidopsis, PSR-associated transcriptional changes are majorly orchestrated by the MYB-transcription factors PHOSPHATE STARVATION RESPONSE 1 (PHR1) and its closest homolog PHR1-LIKE (PHL1) (Bustos et al., 2010; Sun et al., 2016). These transcription factors bind to partially conserved sequences (*P1BS* elements) present in *PSI*-gene promoters to activate the expression of these genes (Franco-Zorrilla et al., 2004; Rubio et al., 2001; Zhou et al., 2008). In consequence, a *phr1 phl1* double mutant is deficient in *PSI*-gene expression both basally and upon exposure to Pi deficiency (Kuo et al., 2014).

The stand-alone SPX (SYG1/Pho81/XPR1)-domain-containing proteins SPX1/2 act as negative regulators of PHR1/PHL1 functions in maintaining phosphate homeostasis. Following the first report that SPX domains can bind specific InsPs, it was subsequently demonstrated that under Pi-replete conditions, SPX-type proteins require InsP_8_ as a co-factor to bind PHR1 and suppress its transcriptional activity (Dong et al., 2019; Wild et al., 2016; Zhu et al., 2019). Upon Pi starvation, intracellular InsP_8_ levels are reduced, relieving SPX1-tethered inhibitions on PHR1, thus allowing successful transduction of PSR (Dong et al., 2019; Riemer et al., 2021) In InsP_8_-deficient *ipk1-1*, *itpk1-2*, *vih1 vih2*, or *spx* mutants, released PHR1 causes constitutive activation of *PSI*-gene expressions, even under Pi-sufficient conditions, accumulating high endogenous Pi levels (Dong et al., 2019; Kuo et al., 2018; Puga et al., 2014; Riemer et al., 2021; Zhu et al., 2019). Besides the well-established role of InsPs in modulating PSR by allosteric regulation of SPX proteins (Dong et al., 2019; Ried et al., 2021; Riemer et al., 2021; Wild et al., 2016; Zhu et al., 2019), there is recent evidence that the cellular Pi status has a direct role in regulating SPX protein stability. Pi-starvation was shown to promote ubiquitylation of at least two SPX proteins (Lv et al., 2014; Osorio et al., 2019; Zhong et al., 2018). Furthermore, the identification of E3 ubiquitin ligases that target rice SPX4 upon Pi-starvation (Ruan et al., 2019) when consolidated with specific InsP-requirement as co-factors affecting the fate/function of hormonal repressors (Laha et al., 2015; Laha et al., 2022) implies that InsP_7_/InsP_8_ are biochemical regulators of stimulus-dependent substrate stabilities.

In a typical eukaryotic cell, ∼20% of targeted protein degradations are mediated by Cullin RING ubiquitin ligases (CRLs), the largest subfamily among the E3 ubiquitin ligase family (Soucy et al., 2009; Sun et al., 2002). A modular CRL contains a Cullin (CUL) scaffold (CUL1-4 in Arabidopsis), an E2-interacting RING-type Rbx/Roc protein, CUL member-guided substrate adaptor and a ubiquitination target-specified substrate receptor (Hotton and Callis, 2008; Hua and Vierstra, 2011; Zheng et al., 2002). CRLs require covalent modification of a Nedd8 moiety on CULs (a process termed as neddylation) (Sun et al., 2002) for activation and complex formation with specific co-receptors, adapters and with the ubiquitin E2 enzyme for subsequent degradation of targeted substrates (Schwechheimer, 2018). In the absence of cognate substrates, CRLs become vulnerable to self-ubiquitylation and are protected by the Constitutive photomorphogenesis 9 signalosome (CSN), an evolutionarily conserved eight-subunit (CSN1-8) macromolecular complex (Chamovitz and Segal, 2001; Dubiel et al., 2015; Stuttmann et al., 2009; Wu et al., 2006). The CSN5 subunit in this holo-complex contains an MPN^+^/JAMM metalloprotease motif that catalytically removes Nedd8 (deneddylation) from CULs, preventing their self-degradation (Cope et al., 2002). Interaction changes between CRL and CSN affect the cellular ratio of neddylated CUL:unneddylated CUL (CUL^Nedd8^:CUL) (Scherer et al., 2016). CUL deneddylation promotes CRL disassembly from the holo-complex and is essential for its functional recycling and establishing associations with newer substrate co-receptors and adapters. Biochemical modes of a CSN operation, including its dynamic rearrangements upon CRL binding, have been gradually unraveled at the structural and functional levels (Dubiel et al., 2015; Enchev et al., 2012; Lingaraju et al., 2014). From these studies, it is evident that optimal CRL activity necessitates regulated cullin neddylation/deneddylation cycles. Recent breakthroughs have revealed highly localized, coordinated activities of mammalian IP5K with IP6K1 in providing InsP_6_ and InsP_7_ to strengthen and loosen CRL-CSN associations, respectively (Lin et al., 2020; Rao et al., 2014; Scherer et al., 2016). However, despite the reported interactions between the Arabidopsis ITPK1 or of SCF^TIR1^/SCF^COI1^ components with the CSN subunits (Feng et al., 2003; Qin et al., 2005; Schwechheimer, 2001) and the implicated role of InsPs as co-factors in response adaptations that require degradation of specific repressors (Laha et al., 2015; Laha et al., 2022; Osorio et al., 2019), the ternary relationship that directly demonstrates the functional coordination between (any) InsP-kinase with CRL and CSN activities, has not been shown so far.

Here, we show that, in Arabidopsis, the metabolically linked pair IPK1 and ITPK1 form a previously unrecognized association within the CSN holo-complex and moderate CUL1 deneddylation efficiency, likely via InsP_7_ synthesis. The corresponding InsP-kinase mutants display enhanced CUL1^Nedd8^ levels due to deficient deneddylation rates *in vivo* and *in vitro*. Impaired deneddylation in these mutants perturb the equilibrium of selective holo-complex-associated and -free monomeric CSN subunits and CUL1, highlighting an intricate auto-regulatory mechanism in the holo-complex activities. Furthermore, using various pharmacological inhibitors that perturb CUL^Nedd8^:CUL ratios, we demonstrate a molecular link between CSN holo-complex deneddylase activities and activation of PSR. In particular, we show that SPX4, a key negative regulator of PSR, is degraded via the CRL pathway upon Pi-starvation. Our data reveal that the maintenance of CSN deneddylase activity and CRL functions are essential to execute a successful PSR in plants. These results unravel novel activities of the plant IPK1-ITPK1 InsP-kinase pair in regulating CSN-CRLs functions during Pi-homeostasis and in responses to Pi-starvation.

## Results

### Arabidopsis IPK1 and ITPK1 interact with CSN subunits

Basal neddylation ratios of cullins are skewed in mutants of mammalian IP5K and IP6K (Rao et al., 2014; Scherer et al., 2016). Similar occurrences are not known in any plant InsP kinase mutant. Anti-CUL1 blots on total protein extracts from Arabidopsis *ipk1-1* or *itpk1-2* plants detected considerably higher levels of neddylated-CUL1 (CUL1^Nedd8^) than in Col-0 (Figure 1A). In other InsP-kinases mutants, namely *ipk2β-1* and *itpk4-1*, that share a common global InsP_6_ deficiency with *ipk1-1* (Kuo et al., 2018; Laha et al., 2015; Laha et al., 2022; Stevenson-Paulik et al., 2005) CUL1^Nedd8^:CUL1 ratios were similar to Col-0 (Supplemental Figure 1A). Enhanced CUL1^Nedd8^ levels observed in *ipk1-1* or *itpk1-2* were restored to Col-0 levels in the corresponding complemented lines *Myc-IPK1/ipk1-1* or *ITPK1-GFP/itpk1-2*, respectively (Gulabani et al., 2022; Laha et al., 2022) (Supplemental Figure 1B). These results suggest that deficiencies limited and specific to *ipk1-1* or *itpk1-2* plants caused perturbed levels of neddylated CUL1.

**Figure 1.**
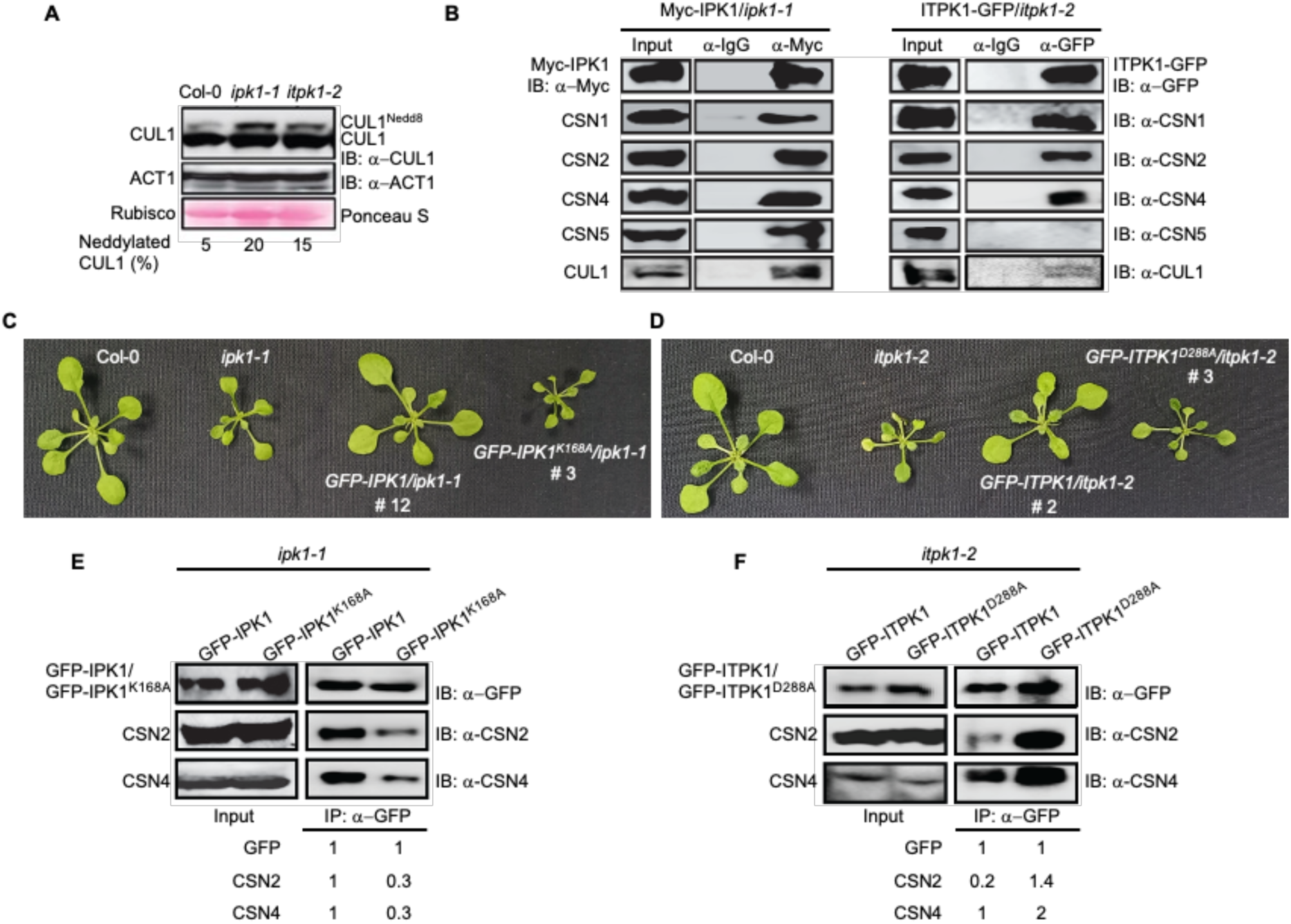
Arabidopsis IPK1 and ITPK1 interact with the CSN holo-complex and modulate its deneddylation activity. **(A)** Anti-CUL1 immunoblot showing basal levels of neddylated: unneddylated CUL1 (CUL1^Nedd8^: CUL1) in *ipk1-1* or *itpk1-2* relative to Col-0. Anti-ACT1 immunoblot or Ponceau S staining for rubisco large subunit denote comparable protein loading across samples. The numerical values below the images indicate the percentage of CUL1^Nedd8^ relative to unneddylated CUL1. **(B)** Immuno-enrichments (IP) of IPK1 or ITPK1 from the corresponding complemented lines (*Myc-IPK1/ipk1-1* or *ITPK1-GFP/itpk1-2*, respectively) probed (IB) with anti-CSN1/2/4/5, anti-CUL1, anti-Myc, or anti-GFP antibodies. IgG-agarose enrichments were used as negative controls. Respective protein amounts in the input extracts are also indicated. **(C and D)** Growth phenotypes of indicated plants at 21 days post-germination. **(E and F)** Immuno-enrichment (IP) of GFP-tagged wild-type or kinase-dead versions of IPK1 or ITPK1 from transgenic plants probed (IB) with the indicated CSN antibodies. The numerical values below the images represent the levels of the probed CSN subunit normalized to the corresponding immuno-enriched InsP-kinase. Respective protein amounts in the input extracts are also indicated.

The COP9 signalosome regulates cullin-deneddylation rates (Lyapina et al., 2001). Previously, mammalian or plant ITPK1 was shown to interact with CSN1 (Qin et al., 2005; Sun et al., 2002). However, their functional implications were not explored. Similarly, human IP5K or IP6K1 enrichments were also shown to co-elute with CSN subunits and cullins (Rao et al., 2014; Scherer et al., 2016). We performed yeast two-hybrid assays to determine physical associations between IPK1 and ITPK1 with various CSN subunits. Both InsP-kinases interacted in yeast with each other and with CSN1 and CSN2 subunits (Supplemental Figure 2A-C). However, only IPK1, but not ITPK1, showed interaction with the CSN5 subunit. Similarly, immuno-enriched Myc-IPK1 or ITPK1-GFP from the above-mentioned complemented plant lines also detected CSN1, CSN2, CSN4 subunits and CUL1 in the co-elutes (Figure 1B). Mirroring the Y2H data, CSN5 was only detected in co-elutes from Myc-IPK1 but not ITPK1-GFP enrichments (Figure 1B). Control pulldowns, either from transgenic plants expressing CaMV 35S promoter-driven GFP *(35S:GFP*) using anti-GFP agarose, or from the InsP-kinase complemented lines using IgG-conjugated beads, did not detect any CSN subunits, validating the specificity of the above interactions (Figure 1B; Supplemental Figure 1C). In bi-molecular fluorescence complementation (BiFC) assays, IPK1 and ITPK1 association with each other or with CSN1, CSN2, or CSN5 were detected as nucleocytoplasmic complexes similar to the localization of the individual proteins (Adepoju *et al*., 2019) (Supplemental Figure 3A). The negative control protein β-Glucuronidase (GUS), did not interact with either IPK1, ITPK1, CSN1, CSN2, or CSN5, again indicating the specificity of the observed interactions (Supplemental Figure 3B). Mutual interaction between IPK1 and ITPK1 was further validated by co-immunoprecipitation assay (Co-IP) wherein immuno-enrichment of transiently expressed GFP-ITPK1, but not of GFP alone, detected the co-expressed Myc-IPK1 (Supplemental Figure 3C). Consolidated, the results demonstrate *planta* associations between IPK1 and ITPK1 with each other and selective CSN subunits.

### Kinase activities of IPK1 or ITPK1 affect their association with the CSN holo-complex

To determine the cellular proportion of IPK1 or ITPK1 pools that associate with the CSN holo-complex, we subjected total protein extracts from Col-0, *Myc-IPK1/ipk1-1* and *ITPK1-GFP/itpk1-2* leaves to size-exclusion chromatography, followed by immunoblotting for the CSN1 subunit, Myc-IPK1, or ITPK1-GFP proteins in the collected fractions. The elution profile of the CSN holo-complex and free CSN subunits from plant extracts have been reported earlier (Dohmann et al., 2005; Gusmaroli et al., 2004). CSN1 was used as the representative marker for CSN holo-complex pools since it exclusively partitions into higher molecular weight fractions (∼400-600 kDa; termed high molecular weight, HMW). Intriguingly, neither Myc-IPK1 nor ITPK1-GFP proteins were detected in fractions containing CSN1, suggesting that under steady-state, only a minor cellular pool of these InsP-kinases is present with the CSN holo-complex (Supplemental Figure 4A). We then investigated whether the activities of these InsP-kinases influenced their association with the CSN subunits. To this end, GFP alone or GFP-tagged wild-type or kinase-dead versions of IPK1 or ITPK1 cDNAs (accordingly named GFP-IPK1, GFP-ITPK1, GFP-IPK1^K168A^, or GFP-ITPK1^D288A^, respectively) were transiently co-expressed with HA-CSN2 in *N. benthamiana* leaves. IPK1^K168A^ harbors a substitution in the conserved kinase motif, whereas the ITPK1^D288A^ contains a similar mutation in the conserved ATP-grasp motif. Thus, these mutant variants are catalytically inactive (Laha et al., 2019; Riemer et al., 2021; Stevenson-Paulik et al., 2005). Anti-GFP immuno-enrichments of the InsP-kinases showed that the kinase-dead IPK1^K168A^ interacted less, whereas more association of ITPK1^D288A^ was observed with CSN2, in comparison to their cognate wild-type proteins (Supplemental Figure 4B). Enrichments of GFP alone did not co-elute HA-CSN2. Interactions between IPK1 and ITPK1 were unaffected by the kinase-site mutations (Supplemental Figure 2C).

Transgenic Arabidopsis *ipk1-1* plants expressing Cauliflower Mosaic virus (CaMV) 35S promoter-driven *GFP-IPK1*, but not *GFP-IPK1^K168A^*, transgene restored growth deficiencies and Pi over-accumulation phenotypes of the *ipk1-1* plants, implying functional complementation by the wild-type, but not by the kinase-dead fusion proteins (Figure 1C; Supplemental Figures 5A, B). These results matched earlier reports (Kuo et al., 2014; Kuo et al., 2018). The lines *GFP-IPK1/ipk1-1#12* and *GFP-IPK1^K168A^/ipk1-1#*3, expressing comparable levels of the respective recombinant proteins, were selected for further assays (Supplemental Figure 5C). Enhanced CUL1^Nedd8^ pools apparent in *ipk1-1*, were restored to wild-type levels in *GFP-IPK1/ipk1-1#12,* but not in *GFP-IPK1^K168A^/ipk1-1#3* extracts, indicating that the maintenance of CUL1^Nedd8^:CUL1 homeostasis requires catalytically active IPK1 (Supplemental Figure 5D). Immuno-enrichments detected lower amounts of co-eluted endogenous CSN2 or CSN4 with GFP-IPK1^K168A^, than GFP-IPK1 (Figure 1E). In Y2H assays, IPK1^K168A^ did not or only weakly interacted with CSN1, CSN2, and CSN5 compared to wild-type IPK1 (Supplemental Figure 2A). When tested similarly for *itpk1-2*, its growth defects were abolished, and high endogenous Pi recovered to wild-type levels, albeit partially, only in the transgenic lines expressing CaMV 35S promoter-driven *GFP-ITPK1*, but not *GFP-ITPK1^D288A^* (Figure 1D; Supplemental Figures 6A, B). Corresponding GFP-ITPK1 fusion proteins were detected in all the above transgenic lines (Supplemental Fig. 6C). We selected *GFP-ITPK1/itpk1-2#7* and *GFP-ITPK1^D288A^/itpk1-2#3* that expressed nearly similar levels of the GFP-fusion protein for further assays. Enhanced CUL1^Nedd8^ level apparent in *itpk1-2* was recovered only in the *GFP-ITPK1/itpk1-2#7*, but not in the *GFP-ITPK1^D288A^/itpk1-2#3* extracts (Supplemental Figure 6D). Further, extracts from these lines immuno-enriched with anti-GFP antibody-conjugated agarose beads detected more endogenous CSN2 and CSN4 in GFP-ITPK1^D288A^ co-elutes than GFP-ITPK1 (Figure 1F). In Y2H assays, ITPK1^D288A^ showed stronger interaction with CSN1 or CSN2 and displayed gain-of-interaction with CSN5, which was not detected for ITPK1 (Supplemental Figure 2B). These results, in accordance with the earlier *N. benthamiana* data (Supplementary Figure 4B), implied that the ITPK1 kinase function may diminish its CSN association. Mutual interaction between IPK1 and ITPK1 was mostly unaffected by the kinase-dead mutations (Supplemental Figure 2C). Taken together, our results suggest that IPK1 and ITPK1 catalytic activities have a reverse effect in their association with CSN. While the kinase function of IPK1 promotes its interaction with the CSN subunits, the ITPK1 kinase activity hinders such equivalent associations. Similar results wherein the mammalian kinase-deficient IP5K^K138A^ protein binds poorly, whereas a kinase-dead version of IP6K1 (IP6K1^K225/335A^) binds more robustly to CSN2 or CSN5 have been reported (Rao et al., 2014; Scherer et al., 2016).

### Equilibrium of association dynamics of the CSN holo-complex is disrupted in *ipk1-1* and *itpk1-2* plants

Locally produced InsPs modulate the tethering efficiencies of cullins to the mammalian CSN (Lin et al., 2020; Scherer et al., 2016). Increased CUL^Nedd8^ levels in the above InsP-kinase mutants may be due to deficient tethering of CUL1 and, hence, decreased deneddylation by the endogenous CSN. To test this, we supplemented Col-0, *ipk1-1,* or *itpk1-2* with purified recombinant CSN2 (His-rCSN2), followed by immuno-enrichments using Ni^2+^-NTA conjugated agarose beads, and then probing for the presence of relative amounts of CUL1 in the elutes. Detection of similar amounts of CSN1 in the His-rCSN2 enrichments from all three extract types suggested the successful and equal incorporation of the recombinant protein into the respective endogenous holo-complex (Figure 2A). Interestingly, higher levels of CUL1 co-eluted with His-rCSN2 in extracts derived from *ipk1-1* or *itpk1-2* plants than in Col-0 samples (Figure 2A). These included pools of both neddylated- and unneddylated-CUL1. Thus, increased CUL1^Nedd8^ levels in the above InsP kinase mutants were not caused by reduced access of CUL1 to the CSN. Control recombinant GFP (His-rGFP) enrichments from the above plant extracts did not lead to the detection of any of the investigated proteins, validating the specificity of the observed interactions (Supplemental Figure 7A).

**Figure 2.**
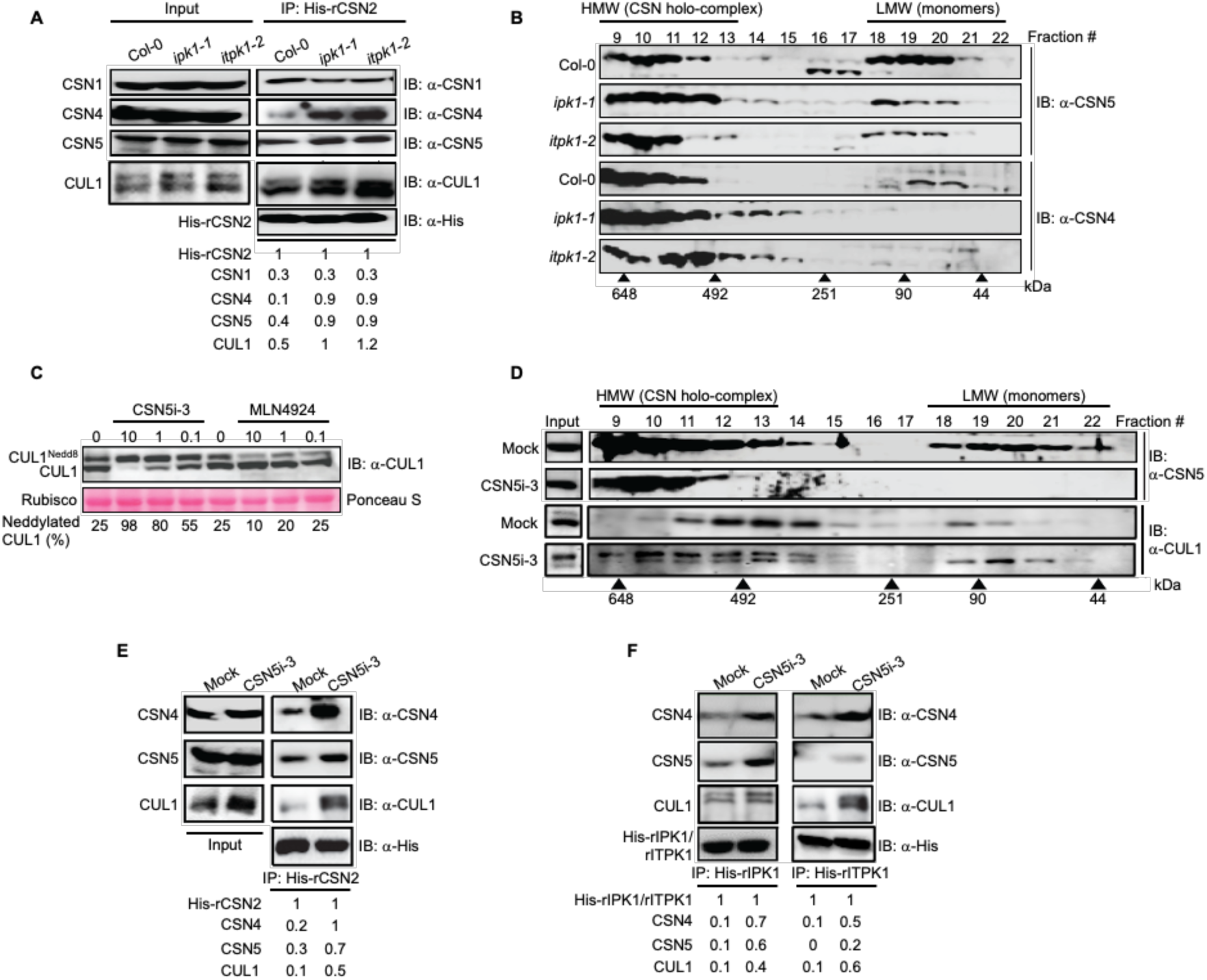
Perturbed InsP-kinase or deneddylation activities disrupt the dissociation equilibrium of CSN4/5 subunits from the holo-complex. **(A)** Affinity enrichment (IP) of His-rCSN2 from Col-0, *ipk1-1*, or *itpk1-2* plants probed (IB) with the indicated antibodies. The numerical values below the immunoblots represent the probed CSN subunit, or CUL1 levels normalized to the corresponding amounts of affinity-enriched His-rCSN2. Respective protein amounts in the input extracts are also shown. **(B)** Size exclusion partitioning profiles of CSN5 and CSN4 from indicated plants probed (IB) with anti-CSN5 or anti-CSN4 antibodies. **(C)** Anti-CUL1 blots from control, CSN5i-3, or MLN4924 treated Col-0 samples at 24 hours post-treatment (hpt). Relative migration positions of CUL1^Nedd8^: CUL1 are indicated. Ponceau S staining shows comparable protein loading across samples. The numerical values below the immunoblot images represent the percentage of CUL1^Nedd8^ relative to unneddylated CUL1. **(D)** Size exclusion partitioning profile of CSN5 and CUL1 from mock (DMSO) or CSN5i-3 treated Col-0 probed (IB) with anti-CSN5, or anti-CUL1 antibodies. **(E)** Affinity enrichment (IP) of His-rCSN2 from control and CSN5i-3-treated plants probed (IB) with the indicated antibodies. The numerical values below the immunoblots represent the CSN subunit or CUL1 amounts relative to the levels of affinity-enriched His-rCSN2. **(F)** Affinity enrichment (IP) of His-rIPK1 or His-rITPK1 from control and CSN5i-3-treated plants probed (IB) with the indicated antibodies. The numerical values below the immunoblots indicate the CSN subunit or CUL1 amounts normalized to the corresponding affinity-enriched recombinant InsP-kinase. In all fractionation data, the fraction numbers and elution positions of molecular weight standards (in kDa) are marked. HMW (high molecular weight) or LMW (low molecular weight) refers to the expected partitioning positions of CSN holo-complex and subunit monomers, respectively. Protein amounts in the input extracts are also indicated for the respective blots.

Selective subunits such as CSN4 and CSN5 are known to undergo dynamic association/dissociation cycles to/from the CSN holo-complex to regulate its deneddylase activity (Dohmann et al., 2005; Enchev et al., 2012; Gusmaroli et al., 2007; Lingaraju et al., 2014; Mosadeghi et al., 2016). Eluates from the above His-rCSN2 enrichments detected more endogenous CSN4 and CSN5 from *ipk1-1* or *itpk1-2* extracts than Col-0 (Figure 2A). Total CSN1, CSN4, or CSN5 levels remained unchanged between the extracts from the different genotypes. We further validated this observation by analyzing the fractionation profile of CSN subunits in Col-0 versus *ipk1-1* and *itpk1-2* extracts via gel-filtration chromatography. Fraction numbers that detected CSN1 were similar between extracts (Supplemental Figure 7B). However, CSN5 and CSN4 profiles from *ipk1-1* and *itpk1-2* revealed comparatively reduced or almost negligible presence, respectively, of these subunits in the lower molecular weight fractions (∼100-50 kDa, termed LMW), as compared to Col-0 (Figure 2B). Fractionated extracts from the complemented lines of these two InsP-kinase mutants did not display these differences, indicating that the alterations were due to consequences of *ipk1-1* and *itpk1-2* mutations (Supplemental Figure 7C). Slight fractionation shifts of CUL1, also towards larger HMW pools, were detected in extracts from *ipk1-1* and *itpk1-2* plants, as compared to Col-0 (Supplemental Figure 7D). These likely represented the CUL1 pools that were associated more with the CSN (Figure 2A). *In toto*, these results implied that a steady-state association dynamics of specific CSN subunits and CUL1 association with the holo-complex is likely perturbed in *ipk1-1* or *itpk1-2* plants.

### Impaired deneddylation activity disrupts CSN equilibrium in the InsP-kinase mutants

Deneddylase activity of CSN5 is active only when it engages with the CSN holo-complex (Gusmaroli et al., 2007). Thus, more CSN5 detected as a part of the holo-complex in *ipk1-1* or *itpk1-2* plants would ideally imply higher deneddylation efficiencies. However, increased CUL1^Nedd8^ levels detected in these mutants contradict this speculation. Indeed, *in-lysate* assays (Lin et al., 2020) demonstrated decreased efficiencies of CUL1-deneddylation in extracts from *ipk1-1* and *itpk1-2* plants, coinciding with an increased CUL1 degradation in these mutant lines (Supplemental Figures 8A, B). Thus, impaired CSN deneddylase activities in *ipk1-1* or *itpk1-2* likely cause augmented CUL1^Nedd8^ levels.

A deneddylation-deficient CSN5 is known to affect the integration of other CSN subunits, such as CSN4, into the holo-complex (Dohmann et al., 2005; Enchev et al., 2012; Gusmaroli et al., 2007). Using CSN5i-3, a recently reported CSN5-deneddylase inhibitor (Schlierf et al., 2016), we then sought to determine the cause-effect relationship between deneddylation deficiencies and altered CSN subunit partitioning. In Col-0 plants, a dose-dependent increment in CUL1^Nedd8^ levels, coinciding with reduced unneddylated-CUL1, was detected at 24-hpt (hours post-treatment) with CSN5i-3 (Figure 2C). MLN4924, a known small molecule pharmacological inhibitor of CUL-neddylation, was used as a negative control in these tests (Hakenjos et al., 2011; Soucy et al., 2009). Interestingly, fractionation of CSN5i-3-treated Col-0 extracts showed markedly reduced CSN5 levels in LMW pools, a pattern overall resembling profiles observed in *ipk1-1* or *itpk1-2* (Figure 2D). The fractionation profile of CSN1 was unaffected by CSN5i-3 (Supplementary Fig. 8C). The enrichment of supplemented His-rCSN2 from CSN5i-3-treated Col-0 extracts detected more CSN4, CSN5, and CUL1 than the control (Figure 2E). These results were also reminiscent of the *ipk1-1* and *itpk1-2* data (Figure 2A), overall suggesting that the deneddylation deficiencies were the primary cause of these retentions in the mutants. Our findings supported earlier reports that a deneddylation deficient recombinant CSN^CSN5-H138A^ binds neddylated CRLs with higher affinity *in vitro* (Enchev et al., 2012; Füzesi-Levi et al., 2020; Mosadeghi et al., 2016).

These results also prompted us to test whether the dissociation dynamics of IPK1 or ITPK1 with the CSN holo-complex are also similarly affected by the deneddylase activity. Indeed, His_6_-tagged recombinant IPK1 or ITPK1 proteins (hereafter termed as His-rIPK1 or His-rITPK1, respectively), when supplemented to CSN5i-3-treated extracts, bound more CSN4, CSN5, or CUL1 than the mock-treated samples (Figure 2F). Thus, the inhibition of CSN5 deneddylation activity also augmented CSN holo-complex associations with IPK1 and ITPK1. This indicates that the deneddylation efficiency of the holo-complex affects the dissociation rates of its mobile subunits CSN4, CSN4 and two InsP-kinases. Interestingly, CSN5i-3 treatments did not affect total InsP_6_ or InsP_7_ levels, implying that deneddylation inhibition does not interfere *per se* with the two InsP-kinase activities (Supplemental Figure 9).

### InsP_6_ potentiation on CSN deneddylase activity requires a kinase-active ITPK1

The kinase activity of IPK1 promotes its CSN association, whereas a kinetically inactive ITPK1 interacts strongly with CSN (Figures 1C, D; Supplementary Figure 2B). The known metabolic relationship between these two InsP-kinases led us to test whether their cognate catalytic products directly affect the deneddylation activity of the CSN holo-complex. In our assays, purified recombinant mammalian Ubiquitin C-terminal hydrolase-L3 (rUCHL3), a well-known deneddylase, was used as the positive control (Hemelaar et al., 2004; Wada et al., 1998). We noted that endogenous CSN holo-complex (HMW Fraction #9-13), enriched from Col-0, was enzymatically active and successfully deneddylated Nedd8-aminoluciferin (Nedd8-AML) substrate *in vitro* in a dose-dependent manner (Figure 3A). At 10 μM concentration, CSN5i-3 completely inhibited the deneddylation activity, thereby ruling out the contaminating activity of other deneddylases, such as a DENEDDYLASE 1 (DEN1) that may be present in the CSN holo-complex pool (Christmann et al., 2013; Mergner et al., 2015). Of greater purity than the enriched Col-0 CSN holo-complex, rUCHL3 was also a more potent deneddylase.

**Figure 3.**
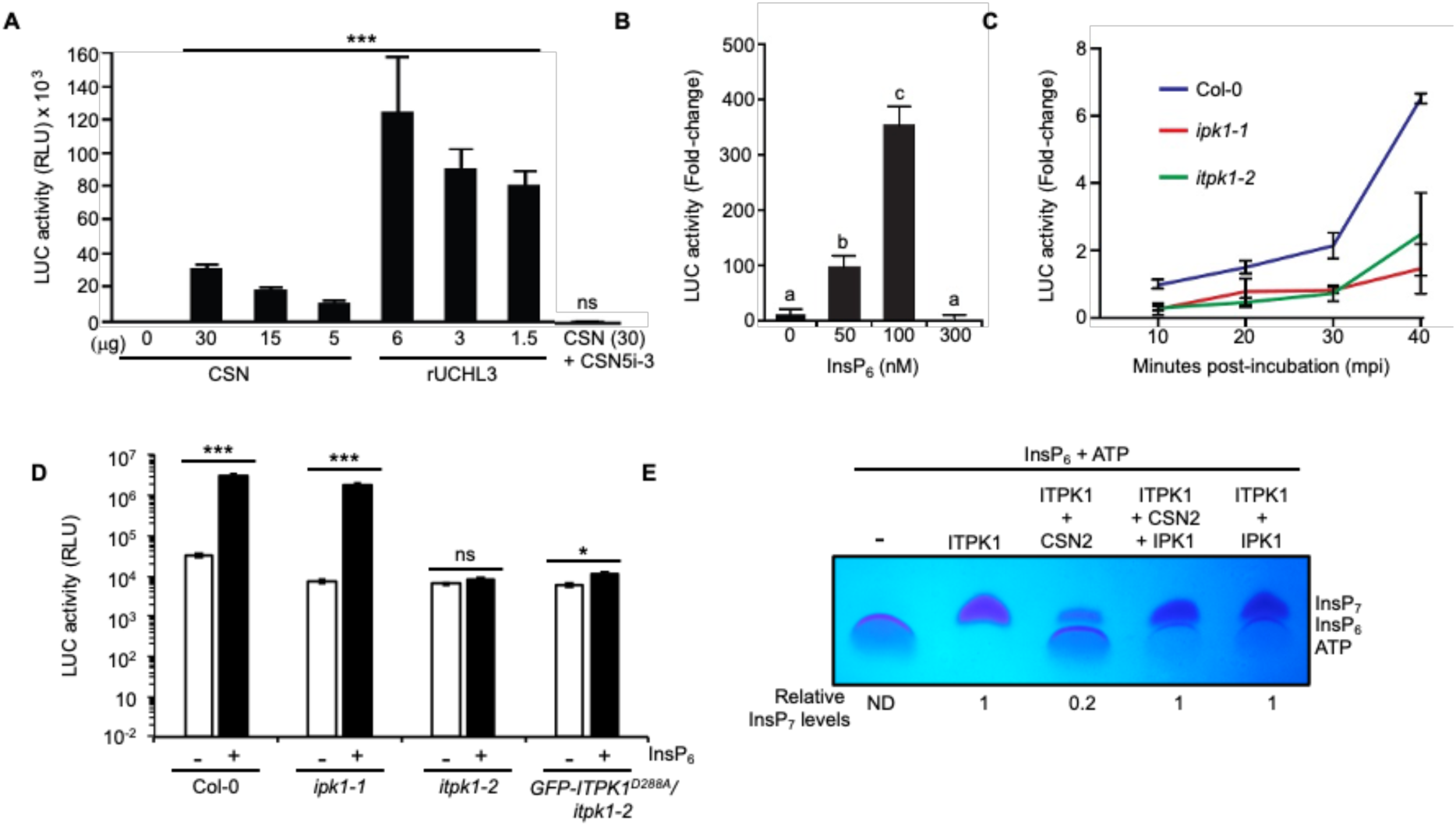
InsP_6_ potentiation on CSN deneddylase activity requires a kinase-active ITPK1. **(A)** *In vitro* deneddylation activity of isolated Col-0 CSN holo-complex on Nedd8-AML substrate. Indicated amounts (in μg) of total CSN holo-complex co-fractionated proteins, or recombinant rUCHL3 were incubated with Nedd8-AML and the Luciferase activity measured. The inhibition assay was performed with CSN5i-3 (10 μM) supplemented to the CSN holo-complex protein pool (30 μg). **(B)** Dose-dependent enhancement in deneddylation activity of isolated Col-0 CSN holo-complex in the presence of indicated amounts of InsP_6_. **(C)** Kinetics of deneddylation rates by the CSN holo-complex isolated from *ipk1-1* or *itpk1-2* plants relative to Col-0. Time points of incubation (mpi, minutes post-incubation) with the substrate are indicated. **(D)** Deneddylation activity of isolated CSN holo-complex from the indicated plants supplemented with or without InsP_6_ (50 nM). **(E)** InsP_7_ synthesis *in vitro* by ITPK1 is inhibited by CSN2 and restored by IPK1. Numerical values below the gel image represent InsP_7_ levels relative to the only ITPK1 reaction. Data shown is mean ± SD (n=3) and represented as relative luminescence units (RLU) **(A and D)**, or relative to minus InsP_6_ sample **(B)**, or as fold-change relative to Col-0 **(C)**. Statistical analysis is according to **(A and D)** Student’s *t*-test with pairwise comparison to minus CSN holo-complex protein samples, (\*\*\**P*<0.0001, ns= not significant), or **(B)** with ANOVA (Tukey, *p* < 0.05).

Because InsP_6_ is commercially available, we first tested whether this metabolite directly affects the deneddylation activity of a plant CSN holo-complex. Remarkably, InsP_6_ addition enhanced, in a dose-dependent manner, the deneddylation activity of Col-0 CSN holo-complex (Figure 3B). Compared to untreated extracts, deneddylation activity was ∼90- or ∼350-fold higher with 50 nM or 100 nM InsP_6,_ respectively. At 300 nM concentration, InsP_6_ inhibited Nedd8-AML deneddylation, likely due to its known protein-precipitation effects (Ried et al., 2021; Veiga et al., 2006). The lack of commercially available 5-InsP_7_, the product of ITPK1 kinase activity, prevented us from directly testing the effect of this metabolite in the deneddylation assays.

To reaffirm the deneddylation deficiencies in *ipk1-1* or *itpk1-*2 extracts noted earlier (Supplementary Figures 8A, B), CSN holo-complex pools were collected from these mutants, as done for Col-0. At similar total protein concentrations normalized with CSN1, *ipk1-1* or *itpk1-2* CSN holo-complex displayed significantly lower basal and reduced deneddylation kinetics on Nedd8-AML, in comparison to Col-0 (Figure 3C). These results validated our earlier findings of deneddylation deficiencies in *ipk1-1* or *itpk1-2* mutants. Most interestingly, we noted that exogenous supplementation of 50 nM InsP_6_ in the deneddylation reaction of *ipk1-1*, but not in *itpk1-2* or *GFP-ITPK1^D288A^/itpk1-2#3*, improved deneddylation rate of the enriched CSN holo-complex significantly (Figure 3D). These results, consolidated with the earlier data (Figure 3B), implied that the supplemented InsP_6_ required an enzymatically active ITPK1, and likely InsP_7_ synthesis, to stimulate deneddylation. Again, the unavailability of commercial 5-InsP_7_ prevented a direct test of this metabolite in rescuing the deneddylation defects of the InsP-kinase mutants.

Similar to the mammalian counterpart, Arabidopsis CSN2 likely binds to InsP_6_ (Lin et al., 2020). Thus, CSN2, either as a direct competitor and/or allosterically, may limit substrate availability for the enzymatic activity of ITPK1 at the CSN platform. The kinase activity-dependent interaction of IPK1 with the holo-complex (Figure 1D; Supplemental Figures 2A, 4B), may ameliorate this suppression either by allosteric interactions with CSN2 or by catalytically providing an increased local concentration of InsP_6_ substrate to ITPK1. We tested this hypothesis by evaluating the efficiency of InsP_7_ synthesis by ITPK1, detected via PAGE (Riemer et al., 2021). As anticipated, wild-type ITPK1, but not ITPK1^D288A^, was enzymatically competent to generate InsP_7_ from InsP_6_ (Supplemental Figure 10A). Remarkably, His-rCSN2, added to the reaction, suppressed InsP_7_ synthesis, whereas the InsP_6_-binding deficient CSN2 (His-rCSN2^K3A^, harboring alanine substitutions of conserved K64, K67, Q68 residues; Lin et al., 2020) did not (Supplementary Figures 10B, C). Most interestingly, His-rIPK1 added to the reaction, mitigated CSN2-mediated inhibition, increasing InsP_7_ synthesis by ITPK1 (Figure 3E). Cumulatively, these results implied that metabolic coupling between the two InsP-kinases that lead to enzymatic activation of ITPK1 may define the rate-limiting step of deneddylation activity of the holo-complex. This activity, in turn, regulates the association-dissociation dynamics of not only the InsP-kinases but also of CSN5 and CSN4 through feedback mechanisms.

### Pi-starvation alters the dynamics of CSN5 and its deneddylase functions

Known consequences in *ipk1-1* and *itpk1-2* mutants are reduced auxin-sensitivity and constitutive PSR (Kuo et al., 2014; Kuo et al., 2018; Laha et al., 2022; Riemer et al., 2021). MLN4924 is known to inhibit the degradation of CRL substrates in Arabidopsis and confer auxin-insensitivity (Hakenjos et al., 2011). To test whether the increased *PSI*-gene expression in the InsP-kinase mutants is linked with deneddylation deficiencies and enhanced CUL^Nedd8^ levels, we treated these mutants with MLN4924. Additionally, upregulated expression of several *PSI*-genes (*PHT1;3, PHT1;2, SPX1*, and *IPS1*), as well as high endogenous Pi-level in these InsP-kinase mutants, were suppressed by MLN4924 (Figure 4A; Supplementary Figure 11A). Similarly, MLN4924 also suppressed the induction of *PSI*-genes in Pi-starved Col-0, implying that changes in neddylation-related processes regulate PSR (Figure 4B). Curiously, a prominent increase in CUL1^Nedd8^ levels that was also reversed by MLN4924 was noted in the Pi-starved Col-0 extracts (Figure 4C). Similarly, the resupply of phosphates to the Pi-deprived media also restored CUL1^Nedd8^ amounts to Pi-replete levels, validating that these responses are consequences of Pi-depravation (Figure 4D). Most intriguingly, the CSN5 partitioning profile under the Pi-deplete condition was peculiarly similar to the *ipk1-1* or *itpk1-2* pattern, with higher levels in HMW than LMW fractions (Figure 4E). Resupplied phosphate restored CSN5 partitioning changes to the Pi-replete pattern. The elution profile of CSN1 was unaltered in the extracts from the Pi-starved plants (Supplementary Figure 11B).

**Figure 4.**
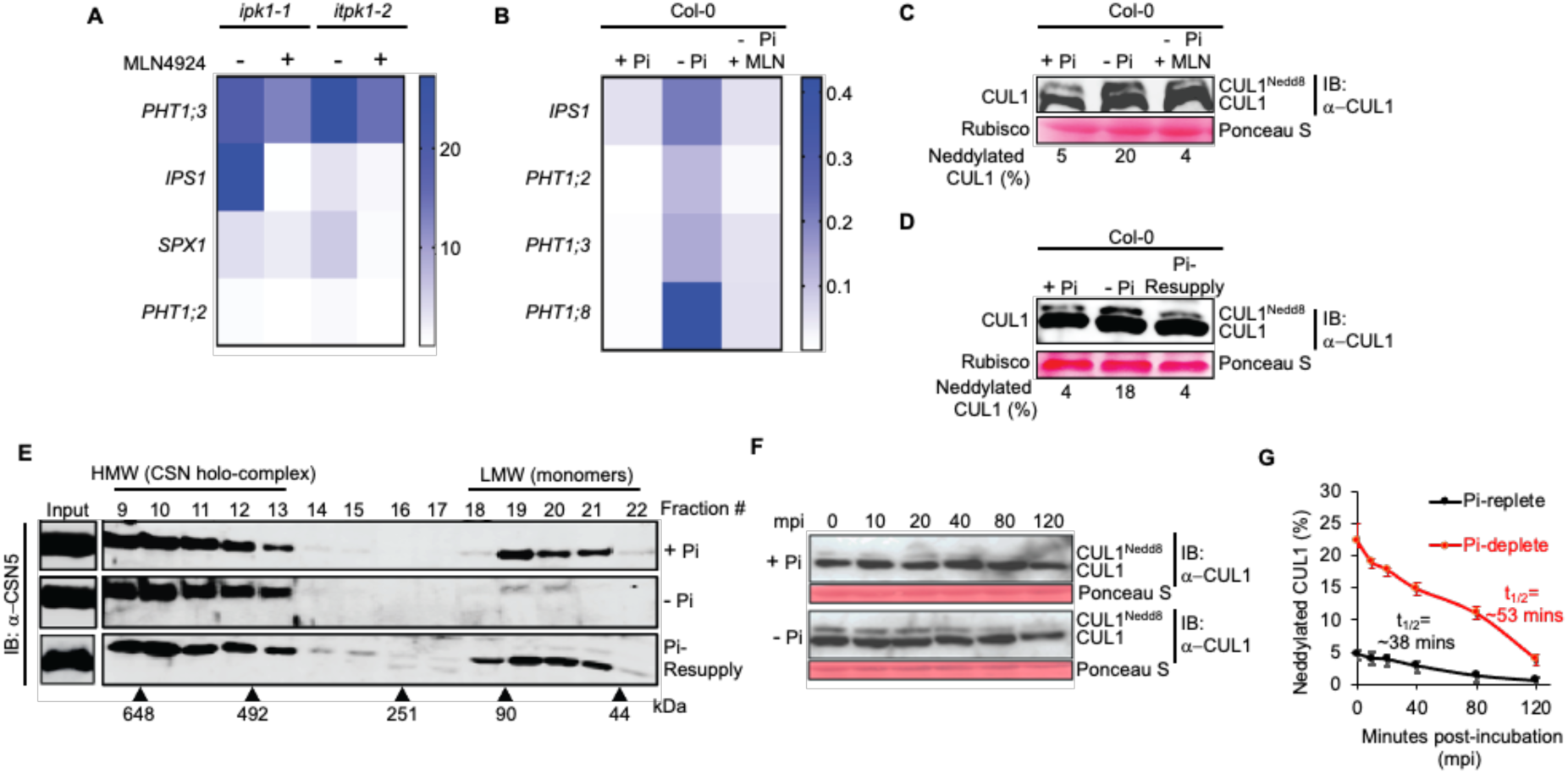
InsP-kinase and CUL^Nedd8^ activities are essential for *PSI*-gene induction during PSR. **(A and B)** *PSI*-gene expression in *ipk1-1, itpk1-2,* or in Pi-replete (+Pi) or Pi-deplete (-Pi) Col-0 in the presence or absence of MLN4924. Gene expressions are relative to *MON1* internal control. Data is mean ± SD (n=3). Heatmaps were generated using GraphPad Prism 8.4.3 software using the single gradient function. **(C and D)** Anti-CUL1 immunoblot showing relative levels of neddylated: unneddylated CUL1 (CUL1^Nedd8^: CUL1) in the indicated samples (+Pi: Pi-replete; -Pi: Pi-deplete; -Pi + MLN: Pi-deplete samples treated with MLN4924; Pi-Resupply: Pi-deplete samples replenished with Pi). **(E)** Size exclusion fractionation profile of CSN5 from extracts of Pi-replete (+Pi), Pi-deplete (-Pi), or Pi-Resupply plants. Fraction numbers, elution positions of molecular weight standards (in kDa), and expected fractions indicative of CSN holo-complex-incorporated (HMW) or free CSN5 subunit monomers (LMW) are marked. Input panels show protein amounts in the total extracts. **(F)** *In lysate* deneddylation assays at indicated time points (mpi, minutes post-incubation) on Pi-replete or Pi-deplete Col-0 extracts probed (IB) with anti-CUL1 antibodies. **(G)** Scatter plot showing the half-life of CUL1^Nedd8^ in the Pi-replete or Pi-deplete Col-0 extracts. In the above immunoblots, the relative migration positions of CUL1^Nedd8^: CUL1 are marked. Ponceau S staining denotes comparable protein loading across samples. The numerical values below images **(C and D)** represent the percentage of CUL1^Nedd8^ relative to unneddylated CUL1.

*In lysate* deneddylation assay, performed as earlier, demonstrated that CUL1^Nedd8^ levels were more stable across progressive time points of reaction in Pi-deplete samples than in Pi-replete extracts, indicative of deneddylation deficiencies upon Pi-starvation (Figure 4F, G). However, unlike the data from the InsP-kinase mutants (Supplementary Figure 8A), gradual CUL1 instability across progressive time points was not apparent in the Pi-deplete samples. Moreover, deneddylation efficiencies on Nedd8-AML substrate by the enriched CSN holo-complex were lower in the Pi-deplete than in Pi-replete samples (Supplementary Figure 11C). Thus, the enhanced CUL1^Nedd8^ levels associated with PSR are caused by dampened deneddylation activities of the endogenous CSN holo-complex and are similar to the occurrence in the two InsP-kinase mutants. Consolidated, these results implied that Pi-starvation dampens CSN deneddylase activity and the elevated CULI^Nedd8^ is essential for the transduction of PSR.

### *csn* mutants or the functional inhibition of CSN5 phenocopy *ipk1-1* or *itpk1-2* properties

Our above results suggested that CSN functions regulate PSR. To determine whether *PSI*-gene expressions are globally affected by functional deficiencies of CSN, we mined the available microarray data of differentially expressed transcripts in the various *csn* null mutants (*csn3*, *csn4* and *csn5ab*) (Dohmann et al., 2008). Strong upregulation in the expression of several *PSI*-genes that were categorized to Pi-remobilization and Pi-transport processes were evident in the transcriptomic data from the *csn* mutants (Supplemental Figure 12). Expression levels of *PHR1* or *PHL1* transcripts remain unaltered in these mutants, similar to *ipk1-1* and *itpk1-2* data (Kuo et al., 2014; Kuo et al., 2018). *IPK1* or *ITPK1* transcript levels are also unaffected in the *csn* mutants. Thus, CSN dysfunctions affect *PSI*-gene expressions.

To explore this further, we utilized the *csn5a-2* plants. Reduced cullin deneddylation in this mutant, evident by the enhanced basal CUL1^Nedd8^ levels, has been previously reported (Gusmaroli et al., 2007), and also validated here (Supplementary Figure 1A). Further, we show that CSN holo-complex isolated from *csn5a-2* plants has reduced deneddylase activity on Nedd8-AML (Supplementary Figure 13A). Similar to *ipk1-1* or *itpk1-2*, the *csn5a-2* is also auxin-insensitive (Dohmann et al., 2005; Dohmann et al., 2008; Gusmaroli et al., 2004; Laha et al., 2022). The primary root length of *csn5a-2* plants grown on Pi-replete growth media is slightly shorter than Col-0, a feature also displayed by the *ipk1-1* or *itpk1-2* plants (Supplementary Figures 13B, C) (Riemer et al., 2021). However, endogenous Pi level in this mutant is comparable to Col-0 with slight up-regulation noticed for *PHT1;8,* or *SPX1* expression under Pi-replete conditions (Supplementary Figures 13D, E). InsP_6_ and InsP_7_ levels were also comparable to Col-0, indicating that *csn5a-2* mutation does not affect global InsP levels (Supplementary Figures 9A-C). Most interestingly, when Pi-starved, the strong inhibition of primary root length observed for Col-0 was markedly lower in *csn5a-2* or *ipk1-1* plants (Supplementary Figures 13B, C) and similar to *itpk1-2* plants as reported earlier (Riemer et al., 2021). Also, similar to the InsP-kinase mutants, the *csn5a-2* plants were also completely impaired in inducing several *PSI-*genes upon Pi-starvation (Supplementary Figures 13E, F). These results indicated that optimal CSN5 functions are essential for transducing PSR.

We also examined the direct effect of CSN5i-3 on Pi-homeostasis and PSR. On Pi-replete Col-0 plants, CSN5i-3 treatment did not dramatically affect *PSI*-gene expressions, implying that inhibited CSN5 or aggravated CUL1^Nedd8^ levels alone cannot trigger PSR (Supplementary Figure 13G). Curiously, upon Pi-starvation, CSN5i-3 suppressed the induction of *PSI*-genes in Col-0 and mirrored MLN4924 effects (Figure 4B), or responses of the *csn5a-2*, *ipk1-1* or *itpk1-2* plants, under similar conditions (Supplementary Figure 13E-G). Overall, these results highlight that unregulated dampening of CSN deneddylase activity negatively impacts the induction of *PSI*-genes during PSR.

### SPX4 is a target of CRLs

Neddylation of cullins activates CRLs, thereby promoting its assembly with the E2 ubiquitin-conjugating enzyme (UBC) and initiating the ubiquitylation of substrate proteins (Cope et al., 2002; Duda et al., 2008). The rice and Arabidopsis SPX4 proteins are known negative regulators of PSR and undergo rapid turnover when the plants are Pi-deprived (Collins et al., 2024; Lv et al., 2014; Osorio et al., 2019; Ruan et al., 2019). Whether SPX4 proteins are *bona fide* targets of CRLs has not yet been directly demonstrated. Increased CUL^Nedd8^ levels prompted us to first evaluate the abundance of global ubiquitin-conjugates in the InsP-kinase mutants. Immunoblot with anti-ubiquitin antibodies detected higher than Col-0 levels of polyubiquitin conjugates in *ipk1-1* and *itpk1-2* extracts but not in their respective complemented lines (Figure 5A). These results suggested that the InsP-kinase mutants display enhanced CRLs-dependent ubiquitylation activities. Interestingly, similar enhancements in the global polyubiquitin conjugates are not apparent in the Pi-starved samples, although a modest increase is detected in the Pi-resupplied extracts (Supplementary Figure 14A). Together, these observations implied that, unlike misregulations that occur in the InsP-kinase mutants, activation of PSR is associated with regulated CRL activities and downstream ubiquitylation of substrates.

**Figure 5.**
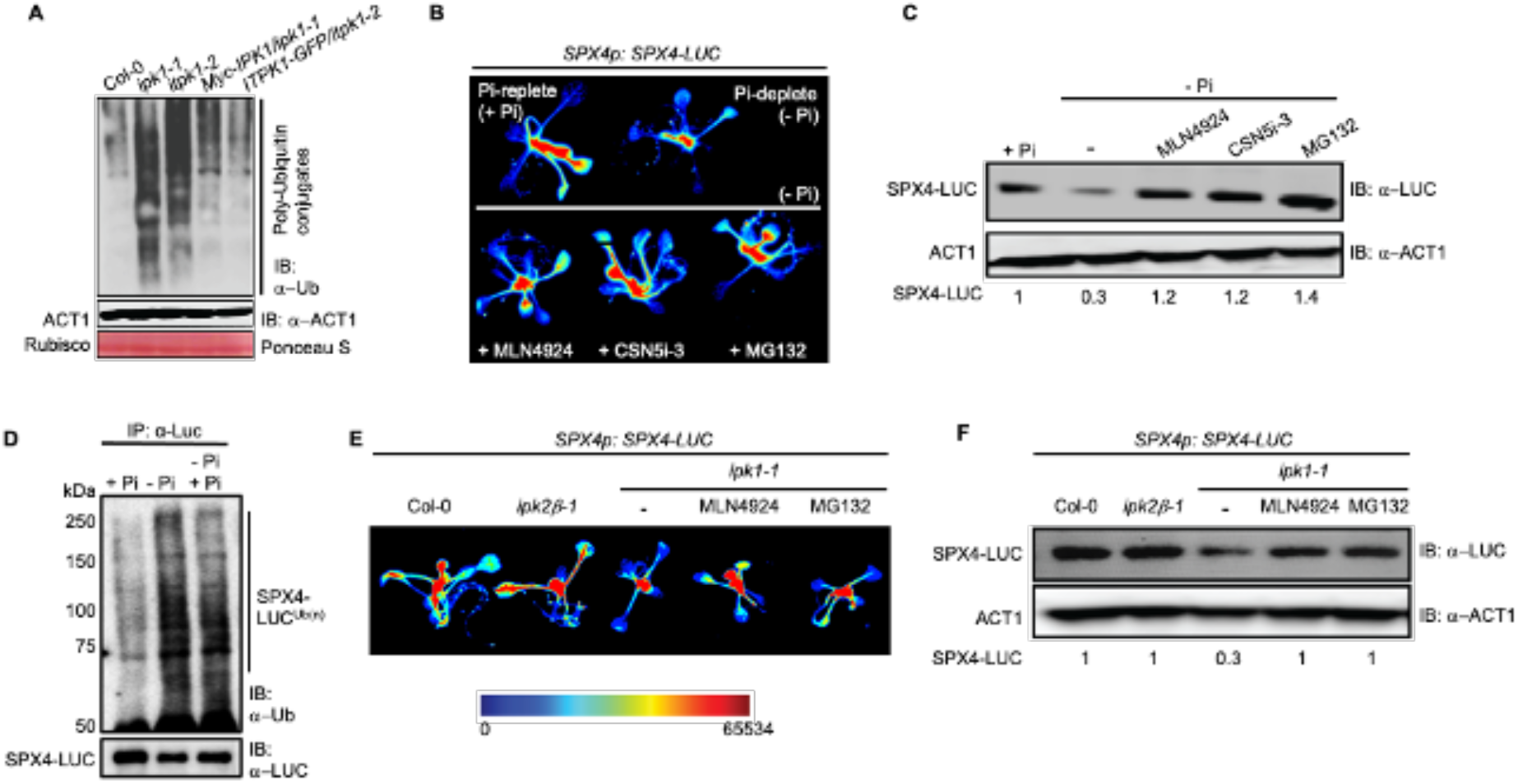
Activated CRLs target SPX4 degradation during PSR. **(A)** Total poly-ubiquitin conjugates in protein extracts of Col-0, *ipk1-1, itpk1-2,* and the respective complemented lines, detected with anti-Ub antibodies. **(B and C)** LUC luminescence and SPX4-protein levels in the *SPX4p:SPX4-LUC* transgenic plants subjected to Pi-replete (+Pi), or Pi-deplete (-Pi) conditions in the absence or presence of MLN4924, MG132, or CSN5i-3. **(D)** Polyubiquitinated SPX4-LUC in +Pi, -Pi and Pi-resupply (-Pi + Pi) detected via immuno-enrichment (IP) with anti-LUC antibodies and probed (IB) with anti-Ub antibodies. Relative levels of unmodified SPX4-LUC are shown with anti-LUC immunoblot. Migration position of molecular weight standards (in kDa) are indicated. **(E and F)** LUC luminescence and SPX4-protein levels in *SPX4p:SPX4-LUC* plants in wild-type (Col-0), *ipk2b-1*, untreated *ipk1-1*, and *ipk1-1* treated with MLN4924 or MG132 inhibitors. Anti-ACT1 immunoblot or Ponceau S staining for rubisco large subunit represents comparable protein loading across samples. Numerical values below the immunoblots **(C and F)** represent SPX4-LUC levels normalized to the corresponding ACT1 levels.

To elucidate whether the activated CRLs target SPX4 upon Pi-starvation, its levels were evaluated in transgenic plants expressing *SPX4p:SPX4-LUC* (Osorio et al., 2019) grown under Pi-replete and Pi-deplete conditions. SPX-LUC protein levels and LUC activity were lower in Pi-deplete versus Pi-replete plants, in agreement with the degradation of SPX4 under Pi-starved conditions, shown earlier (Figure 5B; Supplementary Figure 14B) (Osorio et al., 2019). Global levels of poly-ubiquitylated SPX4 enriched from Pi-deplete plants were considerably higher in Pi-replete than in Pi-deplete plants (Figure 5D). Further, Pi re-supplementation in the Pi-deplete samples caused a noticeable reduction in the levels of poly-ubiquitylated SPX4. Interestingly, treatment of MLN4924, or MG132, a 26S proteasome inhibitor, under Pi-deplete conditions resulted in increased LUC activity and SPX4-LUC protein levels, similar to the Pi-replete plants, indicating that CRL-dependent ubiquitylation indeed regulates SPX4 turnover upon Pi-starvation (Figures 5B, C; Supplementary Figure 14B) (Osorio *et al*., 2019). Most remarkably, the presence of CSN5i-3 in the Pi-deplete medium also suppressed SPX4 turnover and completely restored LUC activity or SPX4-LUC protein amounts to Pi-replete levels (Figures 5B, C; Supplementary Figure 14B). *SPX4* transcript levels were mostly unaffected by Pi-starvation in the presence of these inhibitors (Supplementary Figure 14C). Thus, as earlier, irreversible inhibition of the deneddylation function by the endogenous CSN holo-complex impairs the execution of downstream responses to Pi-starvation.

To test the fate of SPX4 in *ipk1-1*, we generated *SPX4p:SPX4-LUC/ipk1-1* plants by genetic crossing. Likewise, *SPX4p:SPX4-LUC/ipk2β-1* plants were also generated. Imaging showed that LUC activity was significantly reduced in *ipk1-1*, but not affected in *ipk2β-1* (Figure 5E). Similarly, SPX4-LUC protein levels were lower in *ipk1-1*, but unaffected in the *ipk2β-1* background, implying that IPK1, but not IPK2β, regulates the SPX4 turnover (Figure 5F). Further, levels of poly-ubiquitylated SPX4 were higher in *ipk1-1* than in the control or *ipk2β-1* plants (Supplementary Figure 14D). SPX4-LUC protein levels in the *csn5a-2* plants were comparable to the wild-type plants (Supplementary Figure 13H). Transcript levels of *SPX4* were unaffected in either of these mutant backgrounds (Supplementary Figure 14E). Remarkably, SPX4-LUC protein levels and LUC activity in the *SPX4p:SPX4-LUC/ipk1-1* plants were restored to near wild-type levels with either MLN4924 or MG132 treatments, suggesting that enhanced CUL1^Nedd8^ levels caused their heightened turnover in *ipk1-1* (Figure 5E, F). Taken together, our results revealed novel dynamics of CSN-CRL functions modulated by IPK1 and ITPK1 activities in the regulation of plant Pi-homeostasis, steady-state levels of SPX4, and its CRL-mediated turnover during adaptation to Pi-starvation.

## Discussion

### Kinase activities of IPK1 and ITPK1 regulate their self and CUL1 associations with the CSN holo-complex subunits

In this study, we demonstrated that a strategic metabolic coupling of the two Arabidopsis InsP-kinases IPK1 with ITPK1, and more specifically the enzymatic activity of the latter, modulates the deneddylation efficiency of cullins by the CSN holo-complex (Figure 6). Engagements of these two InsP-kinases with the CSN holo-complex are antagonistically affected by their kinase activities. A kinase-deficient IPK1 (IPK1^K168A^) binds weakly, whereas the kinase-deficient ITPK1 (ITPK1^D288A^) associates strongly with the CSN (Figures 1E, F; Supplementary Figures 2A, B, 4B). At the CSN platform, where the two InsP-kinase activities are functionally coupled, IPK1 tethering to the holo-complex is promoted by its catalytic activity in generating InsP_6_. The increased InsP_6_ levels at this platform relieve the enzymatic suppression of ITPK1, caused either by allosteric interaction with the CSN2 subunit and/or the competitive propensity to bind InsP_6_ (Figure 3E). Synthesis of InsP_7_ by ITPK1 (Adepoju et al., 2019; Laha et al., 2019; Riemer et al., 2021; Whitfield et al., 2020) activates the deneddylation activity of the holo-complex, leading to dissociation of this kinase, and likely IPK1, from the CSN platform (Figure 2F). These mechanisms likely define the lack of significant cellular pools of either of the two InsP-kinase enzymes in the CSN holo-complex fractions (Supplementary Figure 3A). The mammalian IP5K and IP6K1 also behave similarly in terms of their kinase-defined associations and dissociation, respectively, with the CSN holo-complex (Rao et al., 2014; Scherer et al., 2016).

**Figure 6.**
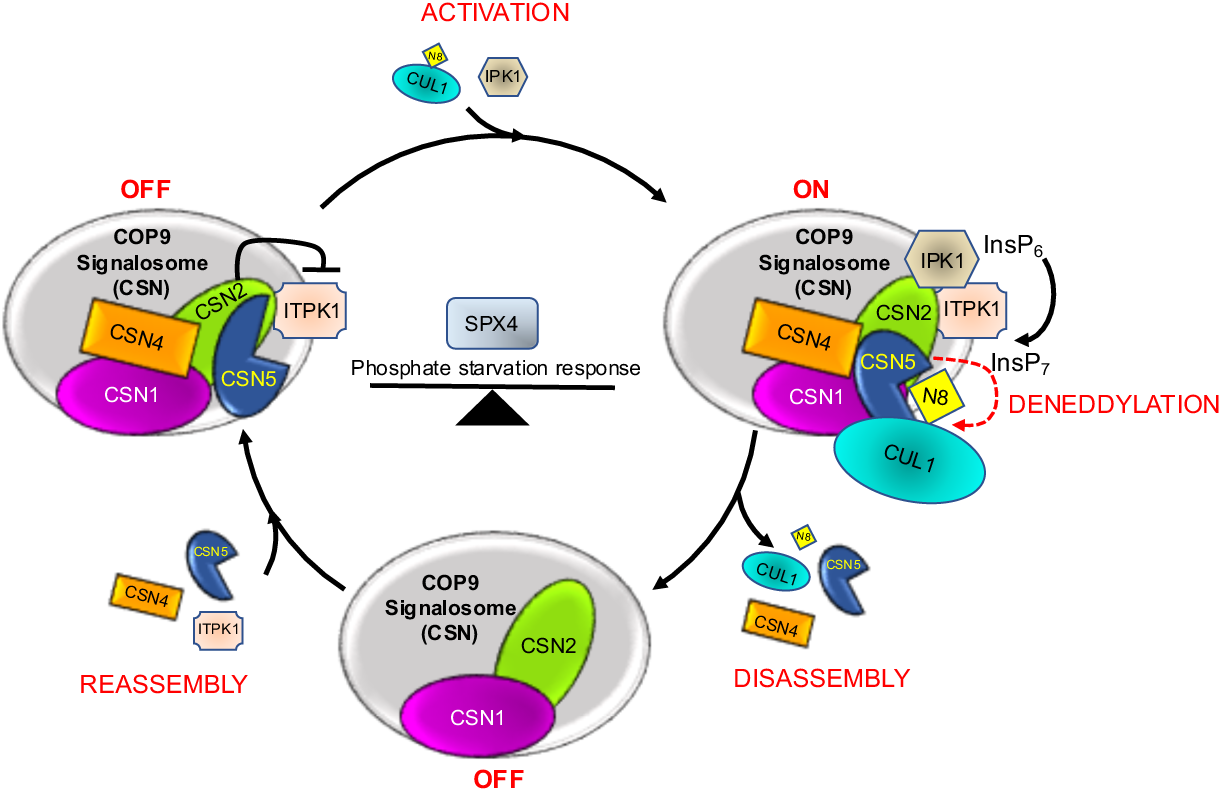
Schematic model of dynamics of COP9 signalosome (CSN) deneddylation cycles regulated by IPK1 and ITPK1. The kinase activity of ITPK1 at the CSN platform is inhibited by CSN2, either through allosteric interactions or via direct competition for InsP_6_ binding. The rate-limiting InsP_7_ synthesis affects the deneddylation rates of the holo-complex. Kinase activity-dependent interaction of IPK1 with CSN2 increases the local synthesis of InsP_6_, relieving the suppression of ITPK1 activity and enhancing subsequent InsP_7_ production. Increased concentration of this metabolite stimulates the deneddylation activity of the holo-complex on cullins (ON). The deneddylation activity leads to the dissociation of deneddylated cullins, CSN4, CSN5 and the InsP-kinases from the holo-complex, thereby causing its inactivation (OFF). The dynamics of these processes maintain the equilibrium of active CRLs that, in turn, regulate the steady-state fate and function of SPX4 in phosphate homeostasis and phosphate starvation response. N8, Nedd8; CUL1, Cullin1.

CUL1 association *per se* with the holo-complex, however, does not seem to require InsP_6_ since more CUL1 was retained with CSN in the *ipk1-1* plants (Figure 2A). This is contrastingly different from the mammalian CUL4-CSN interactions, which are augmented by the InsP_6_ metabolite (Scherer et al., 2016). It is noteworthy that null mutants of Arabidopsis IPK1 are not viable and that the *ipk1-1* used here is an incomplete loss-of-function allele retaining detectable amounts of InsP_6_, although significantly lower than the wild-type levels (Gulabani et al., 2022; Kuo et al., 2018; Laha et al., 2015; Laha et al., 2022; Land et al., 2021; Stevenson-Paulik et al., 2005). Thus, either the residual concentrations of InsP_6_ in *ipk1-1* suffice as a co-factor for the CSN-CRL association, or these interactions in plants have evolved to be independent of this metabolite. Alternatively, changes in other InsPs, such as InsP_5_ 2-OH, which is highly accumulated in *ipk1-1* plants (Gulabani et al., 2022; Stevenson-Paulik et al., 2005), may also be a contributing factor. Further studies need to clarify whether additional InsPs play a role in this process. Mammalian CUL4-CSN interactions *in vitro* are also strengthened by InsP_7_, although at a lower efficiency than InsP_6_ (Scherer et al., 2016). However, again considering that more CUL1 is also retained with CSN2 in *itpk1-2* (Figure 2A), which harbors physiological levels of InsP_6_ but lower level of 5-InsP_7_ (Laha et al., 2022; Riemer et al., 2021), a role of this metabolite in promoting this tether is again unlikely. These observations instead can be interpreted as deficiencies in CUL1 dissociation from the CSN holo-complex due to lower InsP_7_ or inactive ITPK1. The same metabolite is also low in *ipk1-1* due to the low availability of InsP_6_ substrate for ITPK1 (Kuo et al., 2018; Laha et al., 2022; Riemer et al., 2021), thus favoring this interpretation. Indeed, 5-InsP_7_ has been suggested to introduce steric clashes in the interface pocket of the CRL-CSN complex, thereby affecting the stability of this association (Lin et al., 2020). Consolidated, coordinated metabolic activities of the two InsP-kinases affect CUL1 association dynamics with the CSN.

While displaying relatedness to the mammalian IP5K and IP6K1 activities (Rao et al., 2014; Scherer et al., 2016), our results raise the question of how ITPK1, a major InsP_6_-kinase in plants, is evolutionarily endowed with a similar function in coordinating CRL and CSN activities as the mammalian IP6K1, even though these proteins do not share any sequence or structural homology. Perhaps equally surprising is that a mammalian ITPK1 homolog has been reported to engage with the CSN holo-complex (Sun et al., 2002). While it has been shown that mammalian ITPK1 also executes InsP_6_-kinase activity *in vitro* (Laha et al., 2019), it remains to be determined, whether this homolog has a physiological role *in vivo* as an IP_6_-kinase (IP6K) and/or in CRL-CSN activities.

### CSN5 deneddylase activity auto-regulates the functional equilibrium of the holo-complex

Previous studies have shown that the CSN holo-complex is structurally intact but functionally inert without CSN5 presence (Lingaraju et al., 2014). The dynamics of holo-complex-associated and free monomeric forms of CSN5 hence play a vital rate-limiting role in maintaining deneddylation homeostasis and basal levels of active CRLs (Wu et al., 2006). Our data here is the first to identify a correlation between the deneddylation activity of holo-complex with the dissociation equilibrium of the catalyzing enzyme CSN5, and of CUL1, and the two InsP-kinase, with/from the CSN platform. Deneddylation defects in the InsP-kinase mutants, or due to CSN5i-3 treatments, ameliorate their dissociation, retaining increased amounts of these proteins with the holo-complex (Figures 2A, B, D, E). An *in vitro* reconstituted holo-complex containing a deneddylation-deficient CSN5^H138A^ protein was shown to bind neddylated CRLs with higher affinity (Lingaraju et al., 2014). Our findings here reveal a key auto-regulatory mechanism that maintains an equilibrium of deneddylation activity of the plant CSN holo-complex. Holo-complex-free lower sub-complexes of CSN4/5 are known to occur in diverse organisms, including in plants, although their individual roles remain mostly unexplored (Dohmann et al., 2005; Fukumoto et al., 2005; Oron et al., 2002). In mammalian cell lines, CSN5 sub-complexes function as a cargo carrier modulating the nucleocytoplasmic export and subsequent CSN-CRL mediated turnover of the tumor suppressor p53, or the cyclin-dependent kinase inhibitor p27 in the cytoplasm (Oh et al., 2006; Tomoda et al., 2002). Whether, in plants, such CSN5 sub-complexes facilitate similar export of substrates to their subsequent fates upon a stimulus awaits further studies.

### InsP_6_ potentiation on CSN deneddylation activity requires kinase-active ITPK1

The catalytic activity-dependent association of IPK1 with the CSN subunits empirically correlates with the InsP_6_-dependent enhancement of CSN deneddylase activity, which we observed in our assays (Figure 3B). Similar potentiation by InsP_6_ is also seen for the mammalian CSN holo-complex (Lin et al., 2020; Scherer et al., 2016). Structural studies show that holo-complex integrated CSN5 interfaces with CSN4-CSN6 subunits that, in turn, inhibit its deneddylase function in the absence of a bound CRL (Enchev et al., 2012; Lingaraju et al., 2014). Upon binding a neddylated CRL, a series of conformational changes prompted by CSN4 orients the CSN5 catalytic site towards the neddylated cullins. Conserved neighboring lysines in CSN2 α-helices, proximal to the CSN5 catalytic pocket, coordinate InsP_6_ to bridge and stabilize these CRL associations likely for subsequent deneddylation of cullins (Lin et al., 2020). Arabidopsis CSN2 harbors most of these conserved lysines (Supplementary Figure 10B). Therefore, compromised InsP_6_ synthesis in *ipk1-1* plants agrees with the reduced deneddylation rates by the CSN holo-complex (Figure 3B). However, lower InsP_6_ availability would also antagonize the catalytic activity of ITPK1, thereby reducing InsP_7_ synthesis. Our observation that *itpk1-2* plants, which are deficient in InsP_7_ synthesis but have wild-type levels of InsP_6_, exhibit deneddylation defects similar to those in *ipk1-1* plants (Figure 3C) strongly suggests that InsP_7_ is the primary potentiator of the holo-complex’s deneddylation activity. Our data demonstrating that exogenously supplemented InsP_6_ recovers deneddylation defects of *ipk1-1* but not *itpk1-2* or *GFP-ITPK1^D288A^/itpk1-2* supports this interpretation (Figure 3D). The steric clashes that an InsP_7_ molecule is predicted to impose at the CRL-CSN juncture may provide the stimulus to potentiate deneddylation on the bound cullin. This, in turn, would destabilize the complex, causing the dissociation of CUL1, CSN5, and the InsP-kinases from the CSN holo-complex, reverting back to its inert state. We acknowledge that conclusive results can only be obtained from direct tests of InsP_7_ effects in potentiating the deneddylation efficiency of CSN holo-complex and in the recovery of deneddylation defects of *itpk1-2* and perhaps *ipk1-1* plants. However, the limited availability of pure InsP_7_ prevents us from performing these assays.

### Pi-starvation suppresses CSN deneddylation efficiency and targets SPX4 via CRLs

A previous study has demonstrated the proteasome-dependent turnover of SPX4 in Pi-deplete samples (Osorio et al., 2019). However, CSN-CRL activities have not been directly implicated in this turnover or in the downstream response to Pi-starvation. Here, we show that pharmacological inhibition of neddylation with MLN4924 suppresses SPX4 degradation during PSR (Figures 5B, C). The lower SPX4 stability in *ipk1-1* plants is also restored to wild-type levels with MLN4924 or MG132 treatments (Figures 5E, F). Furthermore, MLN4924 suppresses the constitutive PSR and elevated Pi-levels in the two InsP-kinase mutants (Figure 4A; and Supplementary Fig. 11A). In wild-type plants, this inhibitor also dampened the upregulation of *PSI*-gene expression under Pi-deplete conditions (Figure 4B). These data, combined with the observation of elevated CUL1^Nedd8^ levels in extracts of Pi-deficient plants, which are abolished by MLN4924 or restored to basal levels upon Pi resupply (Figure 4C, D), strongly support the idea that CRL activities play a key role in the elicitation of PSR.

Similarly, the very critical modulatory role of CSN deneddylase functions in the transduction of PSR via CRL activities is also emphasized in our data. We show that Pi starvation is associated with reduced deneddylation rates and higher holo-complex associated pools of CSN5, which overall mirror the patterns from the two InsP-kinase mutants (Figures 2B; 4E-G; Supplementary Figure 11C). Reduced cellular levels of InsP_7_, which occur under Pi-deplete conditions (Riemer et al., 2021), also strongly parallel the two InsP-kinase mutants and can be attributed to the observed consequences. Furthermore, the increased CRL activities resulting from their heightened neddylated status likely orchestrate PSR processes, such as SPX4 turnover, as shown in our data.

Most importantly, our study reveals that the physiological response to Pi-starvation involves more complex cellular adaptations, which can be only partially ascribed to functional changes of the two InsP-kinases and the deneddylation activity of CSN holo-complex. This is best evident in the observations that the two InsP-kinase mutants and the *csn5a-2* plants remain recalcitrant to primary root growth inhibition under Pi-deplete conditions (Supplementary Figures 13B, C). Additionally, these mutants and the CSN5i-3-treated plants are inefficient in inducing *PSI*-gene expressions under low Pi (Supplementary Figures 13E-G). Additionally, basal expression levels of *PSI*-genes are not altered either in *csn5a-2* or upon CSN5i-3 treatment on Col-0 (Supplementary Figures 13E, G). Further, global ubiquitination levels are not enhanced in Pi-deplete extracts, unlike those observed in the InsP-kinase mutants (Figure 5A; Supplementary Figure 14A). These results advocate that the constitutive inhibition of CSN5 or of the InsP-kinase activities is detrimental to the successful transduction of PSR. Indeed, under low Pi conditions, ITPK1 was shown to switch its kinase role to an ADP-phosphotransferase that converts InsP_7_ back to InsP_6_ (Riemer et al., 2021; Whitfield et al., 2020). Further investigations are needed to explore the detailed activities of the InsP-kinases and CSN holo-complex under stressed conditions. In conclusion, our studies here highlight the remarkable coordination of plant IPK1 and ITPK1 roles in regulating CRL-CSN functions and their further implications in responding to Pi-imbalances.

## Materials and Methods

### Plant material and growth conditions

Details of *Arabidopsis thaliana* (ecotype Col-0) T-DNA insertion lines *ipk1-1, itpk1-2, ipk2β-1*, *itpk4-1*, *csn5a-2*, *vih2-4, ITPK1-GFP/itpk1-2,* and *Myc-IPK1/ipk1-1* have been described earlier (Gulabani et al., 2022; Jia et al., 2015; Kuo et al., 2014; Kuo et al., 2018; Laha et al., 2015; Laha et al., 2022; Riemer et al., 2021). T-DNA and respective gene-specific primers used for genotyping are listed in Table S1. Seeds were stratified for 2 days at 4°C in the dark and then sterilized using a 30% bleach solution. They were germinated either directly on soil or on ½ strength Murashige and Skoog (MS) Pi-replete (625 µM KH_2_PO_4_) agar plates in growth chambers maintained at 22°C with 70% Relative humidity (RH) and 16 hrs: 8 hrs; light: dark having light intensity 100 μmol μm^-2^s^-1^. For fractionation, Pi-starvation assays, and inhibitor treatments, plants grown on media plates as above were transferred 7 days post-germination to liquid ½ MS medium in 12-well sterile culture plates and proceeded accordingly as indicated in respective experiments. For immunoprecipitation assays, either 4-week-old soil-grown plants or plants subjected to Pi-changes with or without inhibitors for the indicated time period were used.

### Generation of constructs

Arabidopsis *IPK1*, *ITPK1*, *CSN1*, *CSN2* and *CSN5* cDNAs were amplified via RT-PCR (iScript cDNA Synthesis Kit, Bio-Rad, USA) from total RNA isolated from 2-week-old Col-0 plants using RNAiso (Takara-Bio, Japan). Primers for these are listed in Supplementary Table 1. Gateway cloning was first performed into *p*DONR207 and then into *p*BA-Myc (for Myc-IPK1), *p*BA-HA (for HA-CSN2) (Bhattacharjee et al., 2011), *p*SITE-2CA (Chakrabarty et al., 2007) (for GFP-ITPK1), nVenus- or cCFP binary vectors (Bhattacharjee et al., 2011) using BP and LR Clonase, respectively (Invitrogen, USA). BiFC vectors for GUS expression have been described earlier (Bhattacharjee et al., 2011).

For generating wild-type and kinase-deficient constructs of the InsP-kinases (IPK1^K168A^ and ITPK1^D288A^), and InsP_6_ binding-deficient clone of CSN2 (CSN2^K3A^), overlapping PCR with indicated primers using the respective wild-type clones in *p*DONR207 were performed. The confirmed IPK1^K168A^ and ITPK1^D288A^ plasmids were sub-cloned into *p*MDC43 (GFP-containing binary vector (Curtis and Grossniklaus, 2003) using the LR Clonase to generate GFP-IPK1^K168A^ or GFP-ITPK1^D288A^, respectively. The above binary vector clones were electroporated into the agrobacterium strain GV3101 for further *in planta* expression assays.

For generating clones for expression and purification of recombinant proteins (His-GFP, His-rCSN2, His-rCSN2^K3A^, His-rIPK1, or His-rITPK1), the constructs were generated via Clonase LR reaction with corresponding *p*DONR207 vectors and *p*DEST17 (Thermo Fisher Scientific, USA). BL21 (DE3) cells were transformed with the confirmed vectors (or clones), induced, and respective proteins purified using Ni^2+^ affinity resin and size exclusion chromatography. Approximately, 5 μg of indicated proteins were used for enrichment assays.

### Generation of transgenic lines

For generating wild-type and kinase-deficient IPK1 or ITPK1 expressing transgenic plants, *ipk1-1* or *itpk1-2* plants, as appropriate, were floral-dipped with the respective agrobacterium strains (Clough and Bent, 1998). Multiple independent T1 transgenic plants were selected on hygromycin (20 μg/ml) containing ½ MS agar plates, propagated to homozygosity, and then used for indicated assays.

*SPX4p:SPX4-LUC/ipk1-1, SPX4p:SPX4-LUC/ipk2β-1,* or *SPX4p:SPX4-LUC/csn5a-2* plants were generated by crosses between the respective parents. F2 and F3 segregating populations were screened by genotyping PCR to identify the appropriate lines. The primers used are according to previous reports (Dohmann et al., 2005; Gulabani et al., 2022; Osorio et al., 2019) and are also listed in Supplementary Table 1.

### Yeast two-hybrid assays

The indicated constructs were first generated via Gateway cloning into destination *p*BK-RC (bait) and *p*GAD-RC (prey) plasmids (Ito et al., 2000), respectively. The investigated bait and prey combinations were then used to co-transform yeast strain MaV203 using the commercial Yeast Transformation Kit (Sigma-Aldrich, USA), and transformants were selected on SD/LT-media plates. Five independent transformants for each combination were resuspended separately in 100 µl of sterile water, adjusted to similar density, and their serial dilutions were spotted on SD/LT- and SD/HLT-containing 15 mM 3-AT (3-Amino-1,2,4-triazole) plates. The growth of the yeast cells was noted and imaged after 3-days post-incubation. Qualitative X-Gal (5-bromo-4-chloro-3-in-dolyl-β-d-galactoside) and quantitative ONPG (O-nitrophenyl β-d-galactopyranoside) cleavage assays were performed as described earlier (Mockli and Auerbach, 2004). Briefly for X-gal assays, transformants grown in liquid SD/LT-media till 0.8-1 OD_600nm_ were pelleted and permeabilized by three repeats of freeze-thaw cycles of 3 mins each with liquid N_2_ and incubation in a 37⁰C water bath. The pellet was then resuspended in 20 µl of water and added to 96 well plates containing 100 µl of 1X PBS 7.4 pH with 500 µg/ml of X-Gal (Sigma-Aldrich, USA), 0.5% Agarose (w/v), 0.05% (v/v) β-mercaptoethanol and incubated at 30⁰C. Images were acquired at 12 hours post-incubation. For quantitative ONPG assays, transformants harvested as earlier were resuspended in Breaking Buffer (100 mM Tris-HCl pH 8.0, 4% Glycerol, 1 mM β-Mercaptoethanol, 1X Protease inhibitor cocktail; G-biosciences, USA) and an equal volume of acid-washed beads (Sigma-Aldrich, USA). The suspension was lysed via vortexing (3 cycles of 2 min vortex followed by 1 min rest) and then pelleted by centrifugation at 15000 rpm for 15 mins at 4⁰C. To a 96-well plate with each well containing 140 µl Z buffer (60 mM Na_2_PO_4_, 40 mM KH_2_PO_4_, 10 mM KCl, 1 mM MgSO_4_), 20 µl of the supernatant was added. The reaction was started by adding 40 µl of ONPG (4 mg/ml stock; Sigma-Aldrich, USA) to each well, and the plate was incubated at 30⁰C. After 4 hours post-incubation, the reaction was stopped by adding 100 µl of 1 M Na_2_CO_3_ to each reaction well, and absorbance was measured at 420 nm. The total protein content in each lysate was determined by the Bradford method. Miller Unit was calculated using the standard formula (Miller Units = (Abs_420nm_/ ml of cell lysate) * (1000) / (mg of total protein/ ml of lysate) * (reaction time in minutes).

### Phosphate-starvation/re-supplementation assays

After germination of seeds as described earlier, seedlings were transferred to liquid ½ MS medium in 12-well sterile culture plates with Pi-replete conditions (625 µM KH_2_PO_4_) and placed in the growth chamber. At 4 days post-transfer, the plants were either maintained with Pi-replete or transferred to Pi-deplete (10 µM KH_2_PO_4_) liquid media and continued for another 6 days (total of 17 days post-germination) before being used for Pi-estimation, qRT-PCR, total protein extraction, fractionation, or immuno-precipitation assays. For Pi-resupply, plants in Pi-deplete medium were transferred to Pi-replete medium for 24 hrs before analysis.

### Pull down assay/Immunoprecipitation assays

For co-immunoprecipitation assays between Myc-IPK1 and GFP-ITPK1, agro-infiltrated *N. benthamiana* leaves harvested at 48 hpi were used. For pull-downs, including from Arabidopsis transgenic lines *Myc-IPK1/ipk1-1*, *ITPK1-GFP/itpk1-2*, or *35S:GFP* plants, extracts were homogenized in cold RIPA buffer [5 mM Tris-HCl pH 7.5, 150 mM NaCl, 10 mM MgCl_2_, 1 mM EDTA, 1% NP-40, 1 mM Sodium deoxycholate and 1X protease inhibitor cocktail (Sigma-Aldrich, USA)]. After removing cell debris via centrifugation, the supernatant was pre-cleared with IgG agarose beads (Sigma-Aldrich, USA) at 4°C for 1 hr with rotation. The mix was then centrifuged, IgG agarose bead-pellet washed and saved as non-specific controls. The supernatant was added to indicated antibody-conjugated beads (anti-GFP Agarose, BioBharati Life Sciences, India; anti-Myc Agarose, Sigma-Aldrich, USA). After tumbling overnight at 4°C, the mix was centrifuged, and agarose-bead pellet washed 3-times with RIPA buffer. Beads were then resuspended in 1X Laemmli loading dye and used for immunoblots.

### Protein extraction and immunoblots

Total protein extracts from plants were prepared in 6 M urea and used for western blots. For immuno-detection, anti-GFP, anti-Myc (BioBharati Life Sciences, India), anti-CSN1, anti-CSN2, anti-CSN4, anti-CSN5, anti-CUL1 antibodies (ENZO Life Sciences, USA; AS23 4927 Agrisera, Sweden), anti-ACT1 (Abiocode, USA), or anti-LUC (Santa Cruz Biotechnology, USA), as indicated were used. The blots were developed using an ECL kit (Bio-Rad, USA) and imaged in the Alpha Image Quant system (GE-Healthcare, USA).

### Bimolecular Fluorescence Complementation (BiFC) assays

Agrobacterium strains containing the indicated combination of binary vectors were co-infiltrated at equal cell densities into mature *N. benthamiana* leaves. At 48 hpi (hrs post-infiltration), tissue sections from the infiltrated patches were viewed under an SP8 confocal microscope (Leica-microsystems, Germany). Images were acquired with both bright field and YFP filters.

### Size-exclusion chromatography of plant extracts

Gel-filtration protocol was according to (Huang et al., 2013). Briefly, 14-day-old plants ground to a fine powder with liquid nitrogen were homogenized in Gel filtration buffer (50 mM Tris-HCl pH 7.5, 150 mM NaCl, 1 mM EDTA, 10% Glycerol, 10 mM MgCl_2_, 1 mM DTT, 1 mM PMSF and 1X Protease inhibitor cocktail). Homogenates were centrifuged for 10 mins at 4°C, and supernatant passed through 0.2 µM syringe filters. Total protein extracts were fractionated through Superdex 10/300 GL Column (GE-Healthcare, USA) and eluted in the fractionation buffer at 0.2 mL/min flow rate.The eluents were TCA (Trichloroacetic acid) precipitated, washed with acetone, and resuspended in 1X Laemmli loading dye and used for indicated immunoblots.

### *In lysate* deneddylation assay

*In lysate* deneddylayion assay was performed according to Lin et al., (2020). Briefly, 14-day-old seedlings were quickly homogenized in deneddylayion buffer (10 mM Tris-HCl pH7.5, 150 mM NaCl and 0.5% NP-40) in the absence of protease inhibitors. The extracts were incubated at 37°C for indicated times, the reaction stopped by adding Laemmli loading dye to 1X, and then used for immunoblots.

### *In vitro* deneddylation assays

CSN holo-complex was collected by pooling the HMW fractions (#9-12) from the indicated plant extracts via size exclusion chromatography, as earlier (the gel filtration buffer lacked EDTA and protease inhibitors to maintain CSN activity). Total protein was concentrated using Amicon ultracentrifuge filters (Merck Millipore, USA). Nedd8-AML was obtained from Novus Biologicals (USA), and stocks were made in 50 mM HEPES pH 7.5, 100 mM NaCl. For each reaction, ∼15-30 μg protein from the CSN holo-complex pools were used with 1 μM Nedd8-AML in a total reaction volume of 20 μL in Buffer A (20 mM Tris-HCl pH 7.6, 8 mM EDTA, 2 mM ATP). The reaction tube was incubated in the dark at 37^0^C for 30 mins. The entire reaction volume was then transferred to a 96-well luminometer plate, and 80 μL of Assay Buffer B (20 mM Tris-HCl pH 7.6, 8 mM MgSO_4_, and freshly added 2 mM ATP, 6 mg/ml BSA, 33 mM DTT) containing 10 ng of QuantiLum recombinant Luciferase (Promega, USA) was added. For testing effects on deneddylation efficiencies, indicated amounts of InsP_6_ (Sigma-Aldrich, USA), rUCHL3, or CSN5i-3 (10 μM) were added to the reaction prior to the incubation step. Luminescence was measured in a GloMax Luminometer (Promega, USA) and represented as RLU (Relative Luminescence Unit).

### qRT-PCR analysis

cDNAs prepared as earlier, were used for qRT-PCRs with HOT FIREPol EvaGreen qPCR Mix Plus (ROX) (Solis BioDyne, USA) and primer combinations according to the manufacturer’s instructions. Primers used are listed in Supplementary Table 1. The reaction was performed on QuantStudio 6 Flex Real-Time PCR system (Applied Biosystems, USA) with three biological and technical replicates. *MON1* (*At2g28390*) expressions were used as internal control (Gulabani et al., 2022; Kim et al., 2010). Relative expression was calculated according to the PCR efficiency^-ΔΔCt formula.

### Pi-estimation assay

Total Pi measurements are according to (Chiou et al., 2006). In brief, fresh tissue was frozen with liquid nitrogen and homogenized in extraction buffer (10 mM Tris-HCl, 1 mM EDTA, 100 mM NaCl, 1 mM PMSF). The homogenized sample was mixed in a ratio of 1:9 with 1% glacial acetic acid and incubated at 42°C for 30 mins. Then, the reaction was mixed in a ratio of 3:7 with Pi-assay solution (0.35% NH_4_MoO_4_, 0.86 N H_2_SO_4_ and 1.4% ascorbic acid) and incubated at 42°C for 30 mins. Pi content was measured at A820_nm_ using a spectrophotometer. The standard graph was made with known amounts of KH_2_PO_4_.

### Luciferase imaging and quantification

Bioluminescence assays for Luciferase expression and quantification of Luciferase activity (Luciferase assay system, Promega, USA) in the indicated plants were performed according to Osorio et al., (2019). Imaging of LUC expression was done with the Bio-Rad ChemiDoc system (Bio-Rad, USA). Serial dilution of QuantiLum Recombinant Luciferase (Promega, USA) was used to generate the standard graph and values from plant samples presented as fmol/mg total protein.

### Inhibitor treatments

Stock solutions of MLN4924 (Cayman Chemicals, USA), CSN5i-3 (Novartis Inc, Switzerland), and MG132 (Cayman Chemicals, USA) were prepared in DMSO. For treatments,10 µM final concentration of MLN4924 or CSN5i-3 was used. MG132 was used at 50 μM final concentration. For assays on Col-0, *ipk1-1*, or *itpk1-2*, inhibitor treatments were performed 24 hrs prior to analysis. For effects on PSR, inhibitors were added 24 hrs prior to harvesting the samples for analysis.

### *In vitro* ITPK1 kinase, inhibition and recovery assays

The InsP_6_ kinase assay was performed according to the previously published protocol (Riemer et al., 2021) Briefly, 5 μM of purified recombinant ITPK1 (rITPK1) or its kinase-deficient version (rITPK1^D288A^) was used in a reaction buffer containing 5 mM of MgCl_2_, 20 mM of HEPES (pH 7.5), 1 mM of DTT, 12.5 mM ATP, 1 mM of IP_6_ (Calbiochem, USA), 5 mM of phosphocreatine, and 1 unit creatine kinase. After incubation at room temperature for 6 hrs the reaction separated by 33% PAGE and visualized by staining with toluidine blue. To test CSN2 or CSN2^K3A^ (InsP_6_ binding-deficient version) effects on InsP_7_ synthesis, 2 μM of the respective purified recombinant protein (His-rCSN2 or His-rCSN2^K3A^) was added to the InsP_6_ kinase reaction, incubated as above, and analyzed accordingly. Similarly, to test IPK1 effects on CSN2-mediation inhibition on InsP_6_ kinase activity, 2 μM of recombinant IPK1 (His-rIPK1) was added to the reaction and analyzed as earlier.

### Inositol phosphate extraction and CE-ESI-MS analyses

Inositol phosphate extraction from plant tissues was performed as described previously (Pullagurla et al., 2024; Wilson et al., 2015). In short, 12 mg of TiO_2_ beads (GL Sciences Inc., USA) were weighed for each sample and prepared by washing once with ultrapure water and twice with ice-cold 1 M perchloric acid (PA). Using a pestle, the plant tissues were homogenized in ice-cold PA and incubated on ice for 10 minutes, then centrifuged at 18,000 × g for 10 minutes at 4 °C. The collected supernatants were transferred into new 1.5 mL tubes, and the centrifugation step was repeated for another 10 minutes at 20,000 × g at 4 °C. The supernatants were then mixed with the pre-washed TiO_2_ beads and incubated for 30 to 60 minutes at 4 °C on a rotor. After incubation, the TiO_2_ beads in the supernatant were pelleted by centrifugation at 6,000 × g for 1 minute at 4 °C. The pelleted TiO_2_ beads were washed twice with 500 µL of ice-cold PA (1 M). Next, to elute the inositol phosphates, the beads were resuspended in 200 μL of freshly prepared 10% ammonium hydroxide and incubated on rotation for 5 minutes at room temperature. After centrifugation, the supernatants were transferred into 1.5 mL tubes, and the elution was repeated. The eluted samples were lyophilized using a speed-vac. CE–ESI–MS analyses of the plant extracts were performed as previously detailed (Chalak et al., 2024; Qiu et al., 2023).

### Global and SPX4-LUC ubiquitination assays

For determining global polyubiquitination levels, extracts from indicated plants prepared in Gel filtration buffer in the presence of 50 μM proteasome inhibitor MG132 were incubated on ice for 2 hrs. Immunoblotting was then performed with anti-ubiquitin antibodies (P4D1; SantaCruz Biotechnology, USA). For investigating SPX4-LUC ubiquitination, total protein extracts from Pi-replete, Pi-deplete, or Pi-resupply samples were homogenized in Immunoprecipitation (IP) buffer (25 mM Tris-HCl, 7.5 pH; 150 mM NaCl; 0.5% Sodium deoxycholate; 1% Triton X-100; 0.1% NP-40; 40 μM MG-132; 20 mM Iodoacetamide; 10 mM N-ethylmaleimide; 5 μM PR-619 (Cayman Chemicals, USA); 1 mM PMSF; 2X plant protease arrest). The homogenates were centrifuged at 11000 g at 4°C for 5 mins to remove cell debris. To the supernatant, anti-luciferase antibodies conjugated to protein A^PLUS^ beads (BioBharati Life Sciences, India) were added and kept for binding at 4°C with constant rotation for 5 hrs. The samples were then centrifuged at 2000 rpm for 5 minutes at 4°C to pellet the beads. The supernatant was discarded, and the beads were washed three times with IP buffer (without inhibitors). After the third wash, the beads were resuspended in 2X Laemmli loading dye and used for immunoblots.

### Densitometric analysis

The densitometric analysis of the protein bands was performed using the ImageJ Software (NIH, USA). The values presented are normalized to the indicated controls.

### Statistical analysis

All data presented here are representative of at least three biological and technical replicates for each. Statistical analysis details are mentioned in the respective figure legends.

## Funding

This work was supported by financial support from Regional Centre for Biotechnology (RCB) core funds and Grants (No. BT/PR45561/AGIII/103/103/2023 and EMR/2016/001899) to SB and SB, Young Investigator award and DBT-RA fellowship (DBT-RA/2021/January/N/218) to YW, and SERB N-PDF award (No. PDF/2023/001938) to BCS. DL acknowledges the Indian Institute of Science for start-up funds and Infosys Young Investigator award (IISc). G.S. acknowledges funding from the Deutsche Forschungsgemeinschaft under Germany’s Excellence Strategy – EXC 2070 – 390732324 (PhenoRob) and grant SCHA 1274/5-1.

## Author Contributions

SB, YW, MN and SB conceived and designed the research. SB generated all transgenic lines used here. SB and YW generated all constructs and performed most experiments unless otherwise indicated. IB performed the Y2H assays. MN performed all PAGE assays. SB, YW, and MN performed SPX4-LUC experiments. YW, MN, and SD purified recombinant proteins used here. BCS contributed to several size exclusion assays reported here. SB, YW, and MK performed qRT-PCR assays on InsP-mutants upon Pi-changes. InsP extraction and measurements were done by NJP, DL and HJ. SB, DL and GS analyzed the data. SB wrote the manuscript with inputs from all authors.

## Acknowledgements

We thank Dr. Ricarda Jost, La Trobe Institute for Agriculture and Food, Australia, for *SPX4p:SPX4-LUC* seeds, Dr. Tushar Maiti, RCB, for rUCHL3, Dr. Eva Altmann, Novartis Institutes for BioMedical Research, Switzerland for sharing CSN5i-3, Swaroop Peddiraju and Dr. Divya Chandran, RCB for help with transcriptomic data analysis of *csn* mutants, Abhisha Roy for specific SPX4-LUC assays, and Dr. Nripendra Singh and his team at Advanced Technology Platform Centre (ATPC), Faridabad, India for constant support with sample fractionations, and Marília Kamleitner, Department of Plant Nutrition, Institute of Crop Science and Resource Conservation, University of Bonn, for critically reading this manuscript. The authors declare no conflict of interest in this study.

---

**Supplementary Figure 1.**
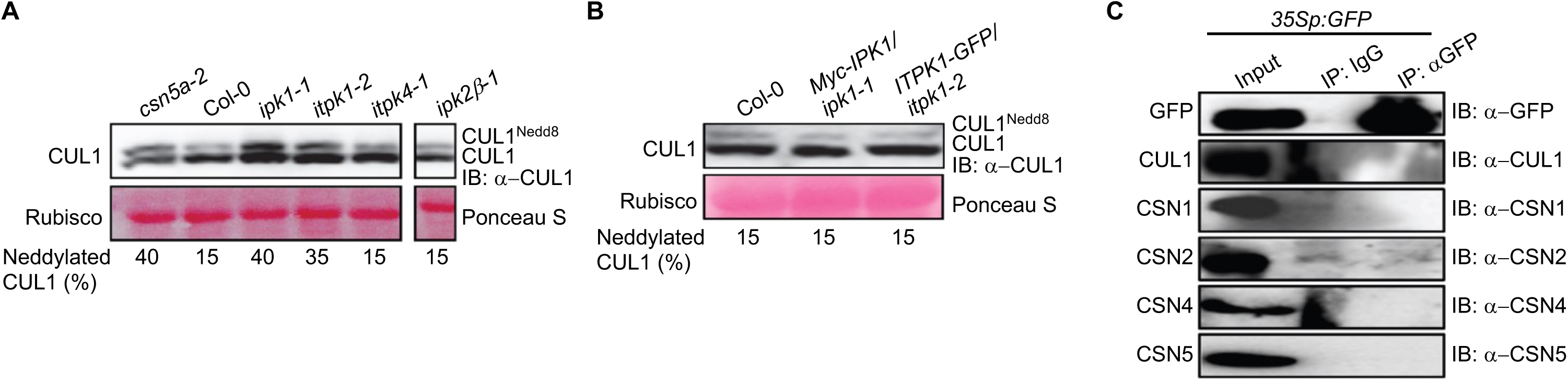
Increased CUL1^Nedd8^: CUL1 ratios are specific to *ipk1-1* or *itpk1-2* but not other InsP-kinase mutants. (A and B) Anti-CUL1 immunoblot showing relative levels of neddylated: unneddylated CUL1 (CUL1^Nedd8^: CUL1) in the indicated plant lines. Comparable protein loading between samples is shown by Ponceau S staining for rubisco large subunit. The numerical values below each lane represents the percentage of CUL1^Nedd8^ relative to unneddylated CUL1 for the corresponding sample. (C) Anti-GFP or anti-IgG-enrichments (IP) from control *35Sp:GFP*-expressing transgenic plants probed (IB) with the indicated antibodies. Relative protein amounts in the input extracts are also shown.

**Supplementary Figure 2.**
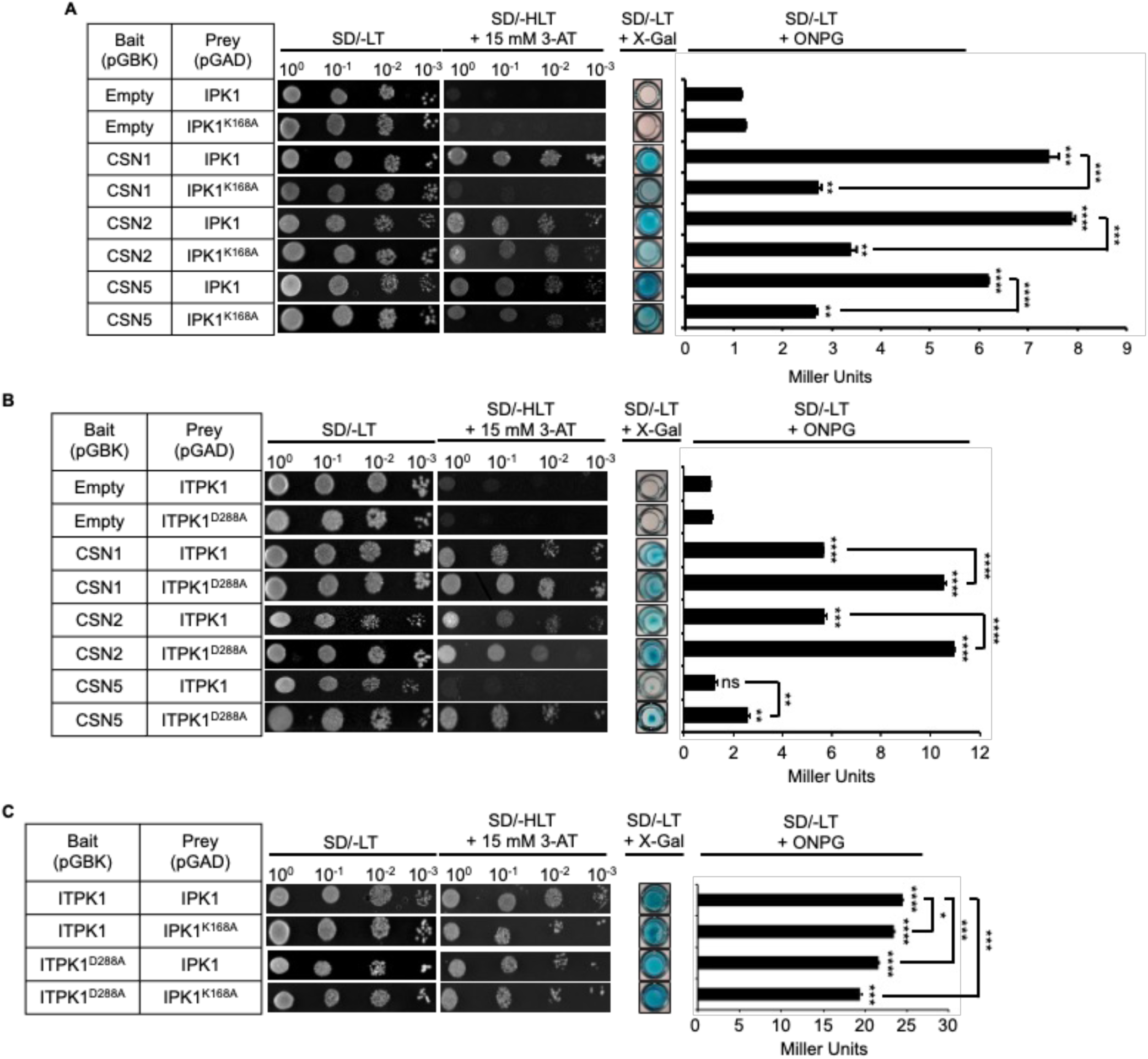
Kinase domains of IPK1 and ITPK1 affect their interaction with CSN subunits, but not with each other. **(A)** Yeast two-hybrid assays showing stronger interaction of kinase-proficient (IPK1) than kinase-deficient IPK1 (IPK1^K168A^) with CSN1, CSN2, and CSN5 subunits. **(B)** Yeast two-hybrid assays showing stronger interaction of kinase-deficient (ITPK1^D288A^) than kinase-proficient ITPK1 (ITPK1) with CSN1, CSN2, and CSN5 subunits. **(C)** Yeast two-hybrid assays showing mutual interaction of kinase-deficient and kinase-proficient IPK1 and ITPK1 (ITPK1) with each other. Yeast transformants containing the indicated bait and prey combinations were plated with the indicated serial dilution on SD/-LT or SD/-HLT + 15 mM 3-AT plates, and the growth of cells was recorded after 3 days. Qualitative X-Gal assays were performed with permeabilized yeast transformants, and the images were acquired 12 hours post-incubation. Quantitative ONPG assays were performed with lysates from the indicated transformants, and color development was measured using a spectrophotometer (420 nm). For each assay , n=3-5. Statistical analysis is according to Student’s *t*-test : firstly, between indicated combinations of the tested InsP-kinase (wild-type or kinase-dead mutant) prey and empty or the indicated CSN subunit bait; secondly, between the indicated combinations of the wild-type InsP-kinase or its kinase-dead prey version with a tested CSN subunit bait; thirdly, between the indicated combinations of wild-type IPK1 and ITPK1 or their indicated kinase dead version (\**P*<0.05, \*\**P*<0.01, \*\*\**P*<0.001, \*\*\*\**P*<0.0001, ns=not significant).

**Supplementary Figure 3.**
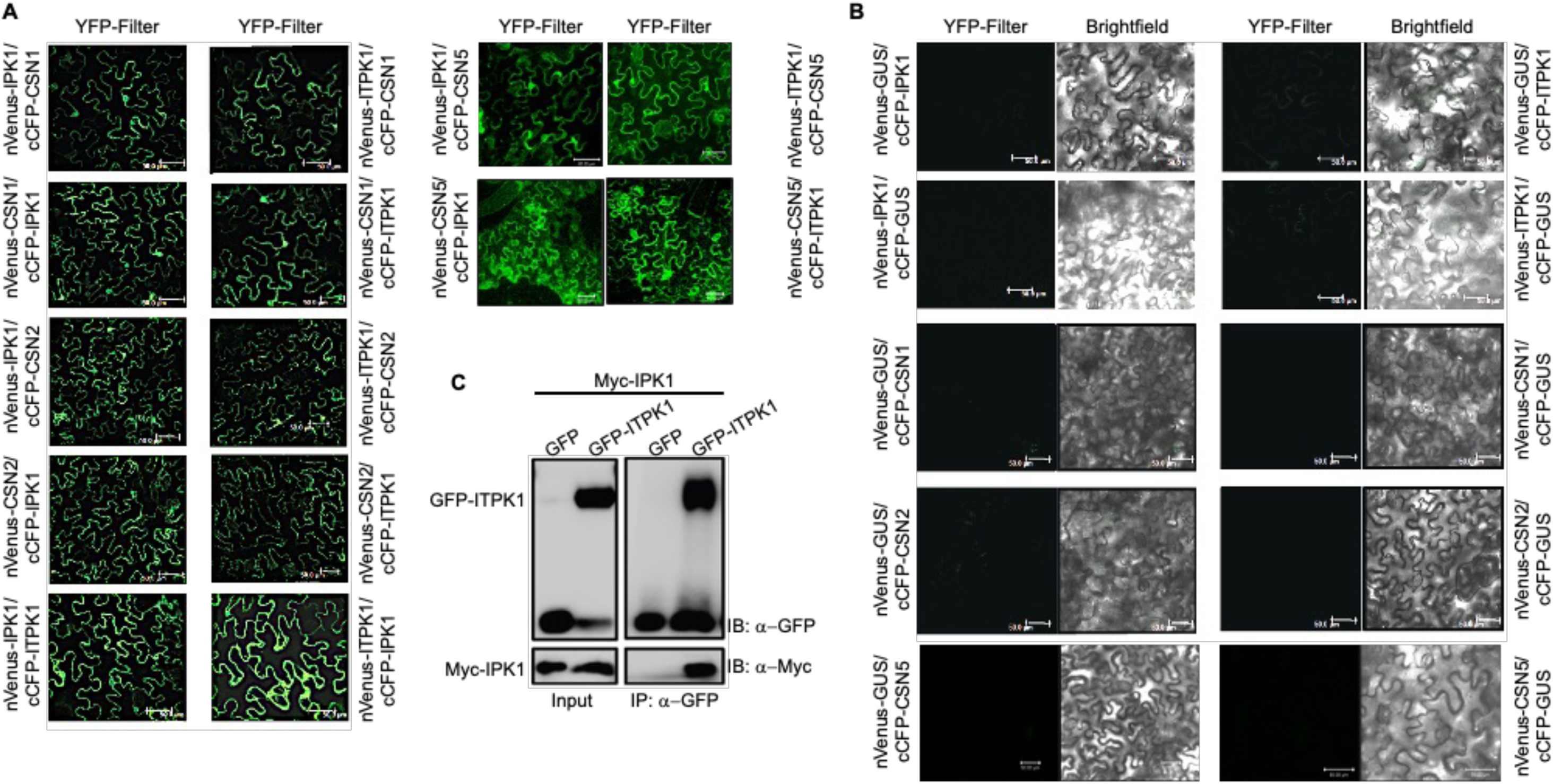
*In vivo* interactions between IPK1, ITPK1 and CSN subunits. **(A)** Confocal images of BiFC assays showing interactions between IPK1 and ITPK1 and with CSN1, CSN2, and CSN5 subunits. **(B)** Lack of interactions between the negative control GUS with IPK1, ITPK1, CSN1, CSN2 or CSN5. Images were acquired at 48 hpi (hours post-infiltration) with YFP and brightfield filters. Figures are representative of multiple images obtained for each BiFC combination. Scale bar = 50 μM. **(C)** Co-immunoprecipitation of transiently expressed GFP-ITPK1 with Myc-IPK1. Anti-GFP pulldown (IP) from GFP alone or GFP-ITPK1 co-expressed with Myc-IPK1 were probed (IB) with anti-GFP or anti-Myc antibodies. Protein amounts in the input samples are indicated.

**Supplementary Figure 4.**
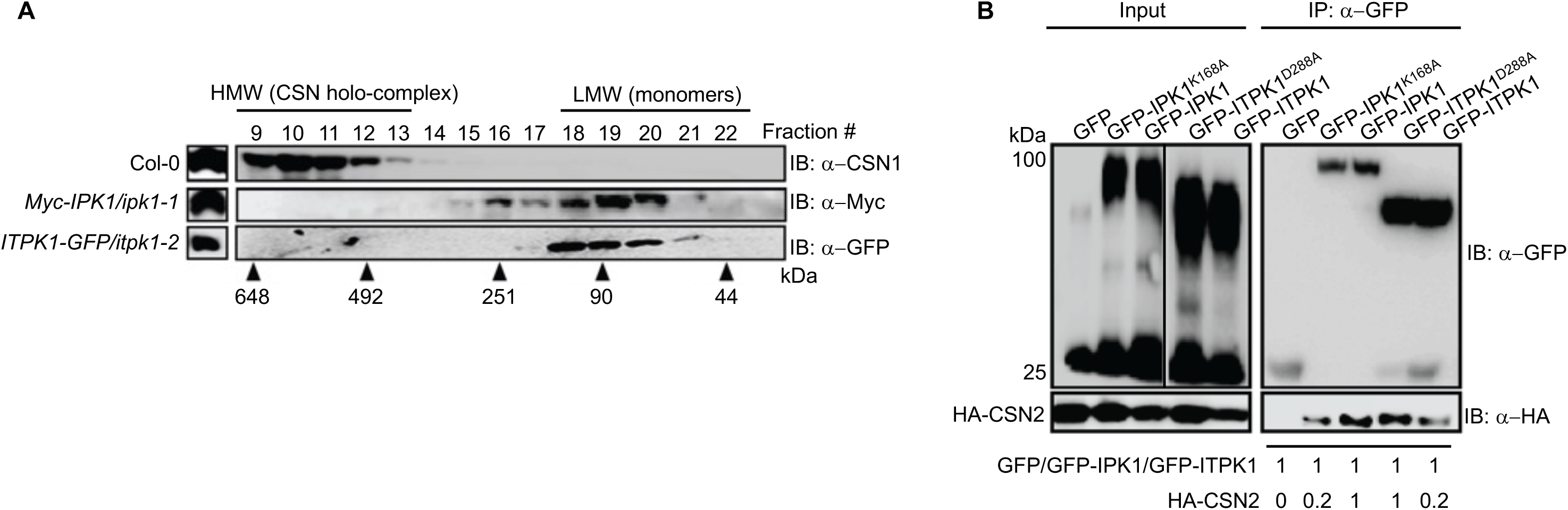
CSN2 associations of IPK1 and ITPK1 are conversely affected by their catalytic activities. **(A)** Gel-filtration fractionation pattern of Myc-IPK1 or ITPK1-GFP probed (IB) with the indicated antibodies. Fraction numbers and elution positions of molecular weight standards (in kDa) are indicated. CSN1 partitioning profile was used as a reference for the elution pattern of the CSN holo-complex. **(B)** Immuno-enrichment of GFP-tagged wild-type or kinase-dead IPK1^K168A^ or ITPK1^D288A^ co-expressed with HA-CSN2 in *N. benthamiana* leaves. Pull-down (IP) was performed with anti-GFP antibodies and the bound proteins were probed (IB) with anti-GFP or anti-HA antibodies. The migration position of molecular weight standards (in kDa) and the total protein amounts in the extracts (input) are shown. The numerical values below the IP image represents the relative CSN2 amounts normalized to the corresponding enriched GFP-tagged InsP-kinase.

**Supplementary Figure 5.**
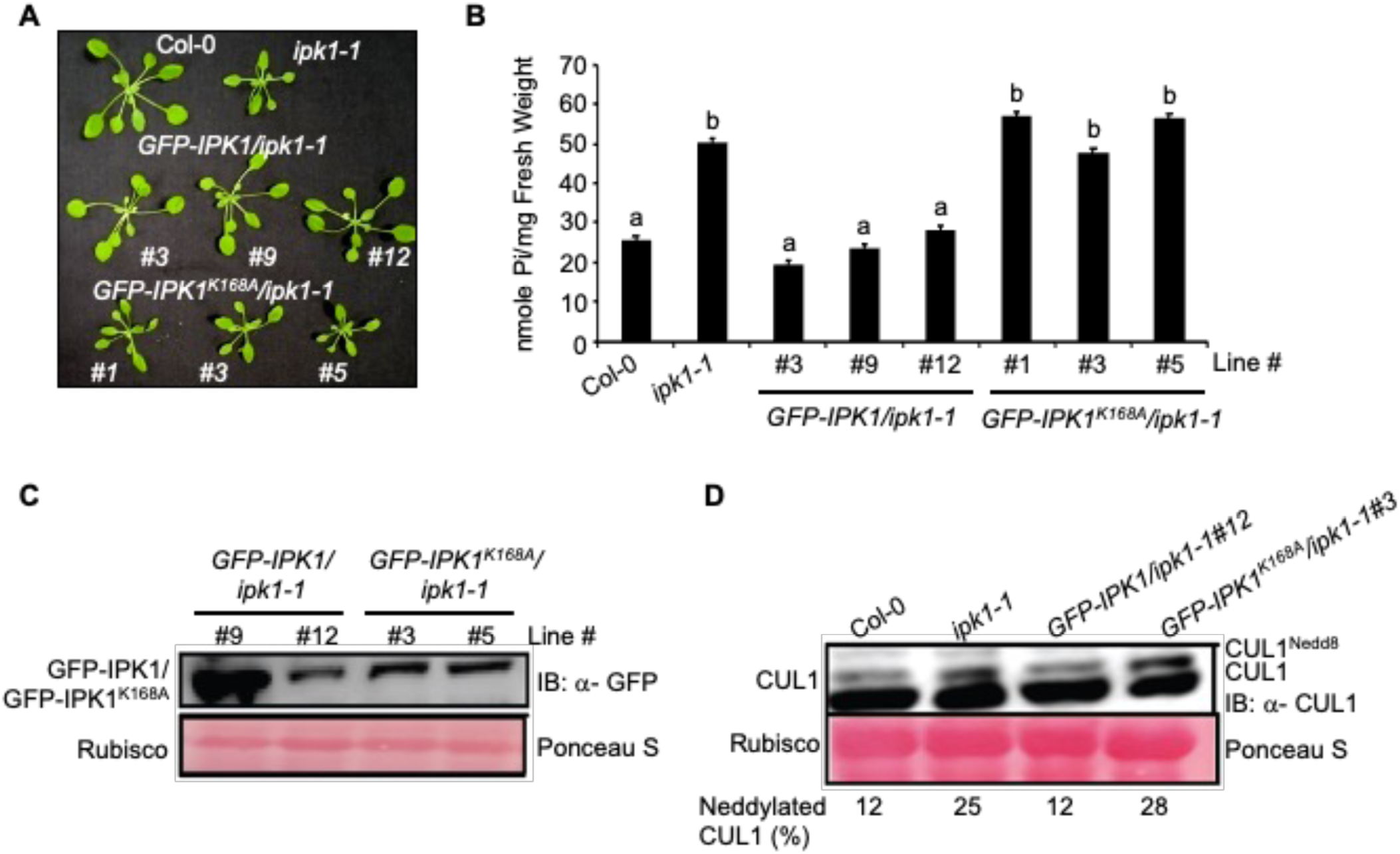
Kinase-proficient, but not kinase-deficient IPK1, restore *ipk1-1* defects. **(A)** Growth phenotypes of independent transgenic lines of *ipk1-1* plants expressing *GFP-IPK1* or *GFP-IPK1^K168A^*. **(B)** Endogenous Pi levels in the indicated genotypes. The data shown is mean ± SD (n=6-7). Different alphabets indicate significant differences (P< 0.05, ANOVA, Tukey) between samples. **(C)** Expression levels of GFP-IPK1 or GFP-IPK1^K168A^ in the indicated transgenic lines probed with anti-GFP antibodies. **(D)** Relative CUL1^Nedd8^:CUL1 levels in the indicted plant lines probed (IB) with anti-CUL1 antibodies. Relative migration positions of CUL1^Nedd8^ and CUL1 are marked. The numerical values below the image represent the percentage of CUL1^Nedd8^ relative to unneddylated CUL1 for the corresponding sample.

**Supplementary Figure 6.**
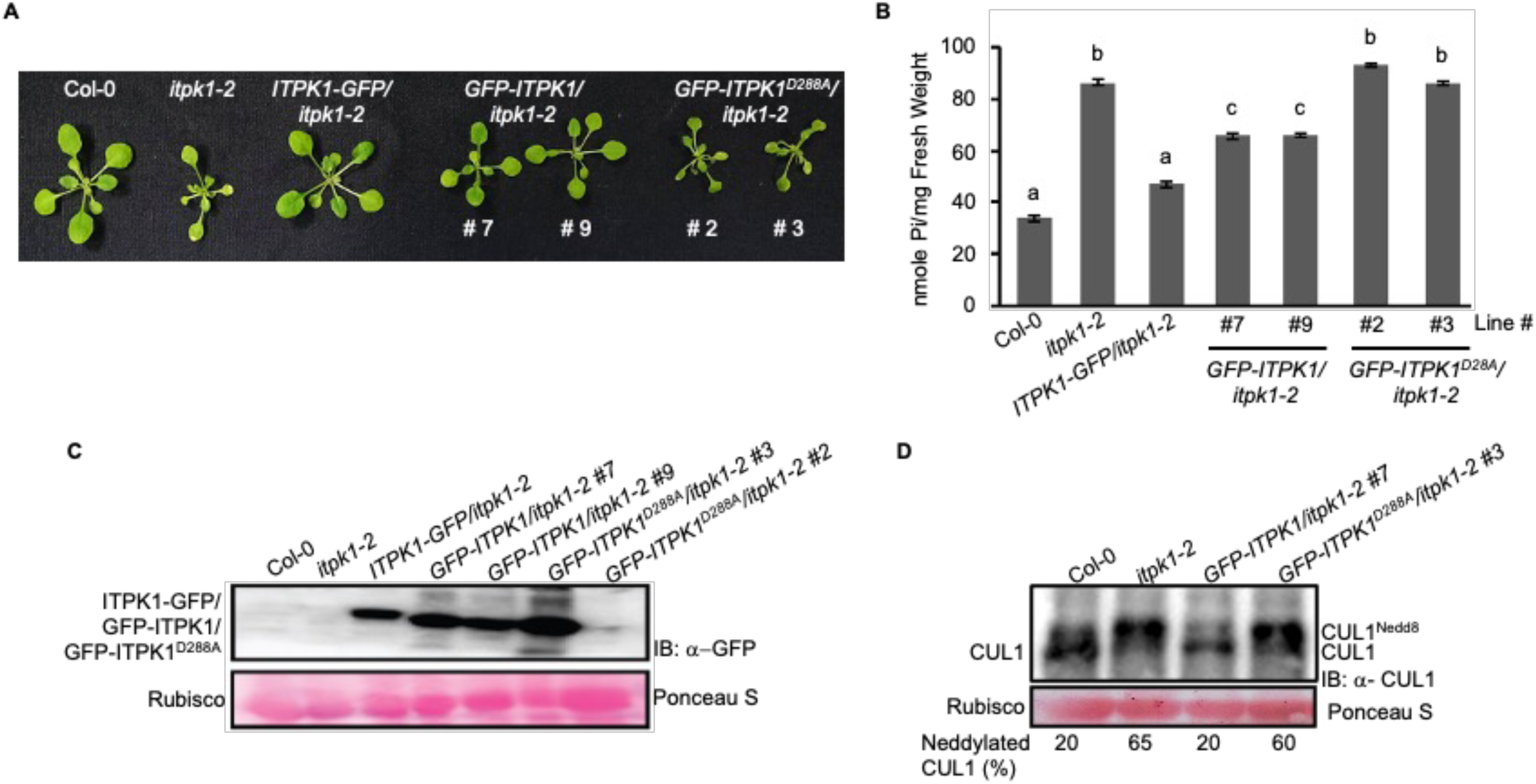
Kinase-proficient, but not kinase-deficient ITPK1, restore *itpk1-2* defects. **(A)** Growth phenotypes of independent transgenic lines of *itpk1-2* plants expressing *GFP-ITPK1* or *GFP-ITPK1^D288A^*. **(B)** Endogenous Pi levels in the indicated genotypes. The data shown is mean ± SD (n=6-7). Different alphabets indicate significant differences (P< 0.05, ANOVA, Tukey) between samples. **(C)** Expression levels of GFP-ITPK1 or GFP-ITPK1^D288A^ in the indicated transgenic lines probed with anti-GFP antibodies. **(D)** Relative CUL1^Nedd8^:CUL1 levels in the indicted plant lines probed (IB) with anti-CUL1 antibodies. The migration positions of CUL1^Nedd8^ and CUL1 are marked. The numerical values below the image represent the percentage of CUL1^Nedd8^ relative to unneddylated CUL1 for the corresponding sample.

**Supplementary Figure 7.**
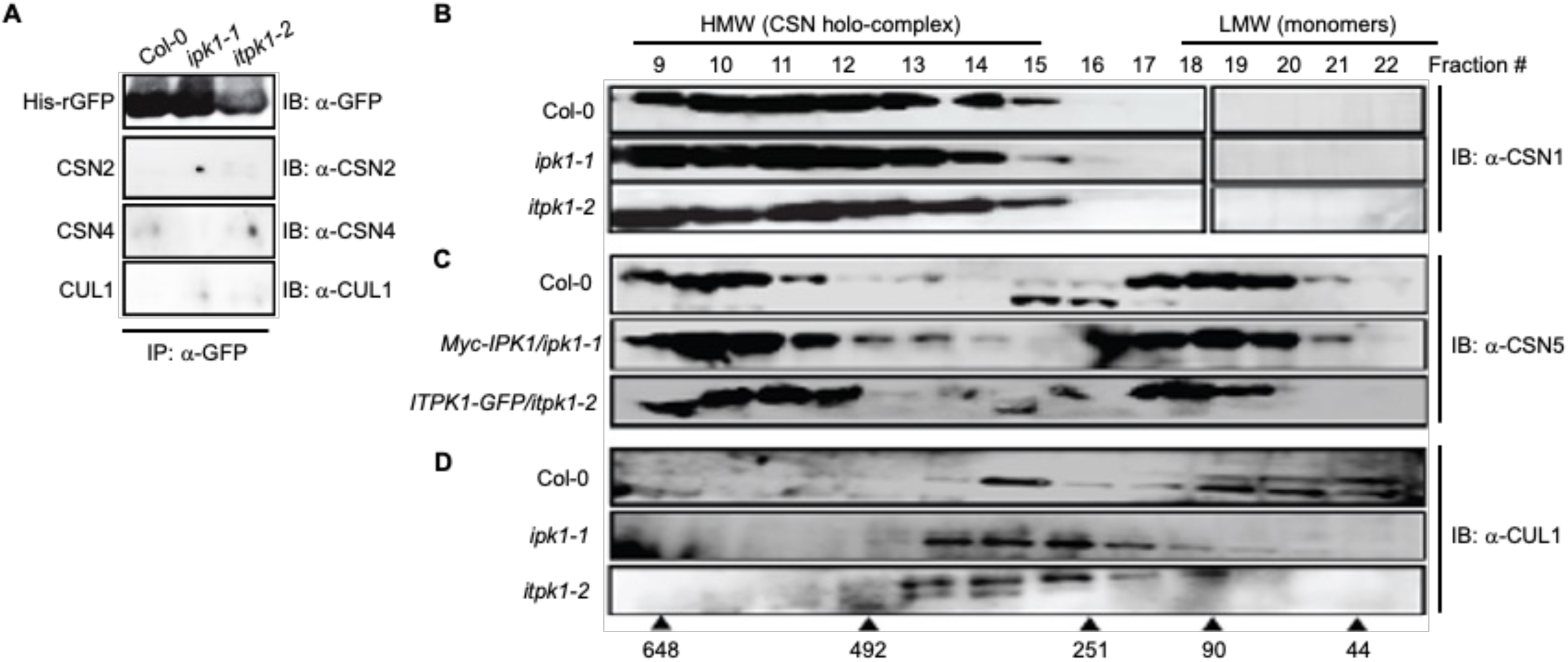
Fractionation profiles of CSN subunits and CUL1 in the indicated plants. **(A)** Control His-rGFP enrichments (IP) from supplemented extracts of indicated plants probed (IB) with the various antibodies. **(B)** Size exclusion partitioning profile of CSN1 from Col-0, *ipk1-1*, and *itpk1-2* extracts. The blots were probed (IB) with anti-CSN1 antibodies. **(C)** Size exclusion partitioning profile of CSN5 Col-0, and the complemented lines of *ipk1-1* and *itpk1-2* (*Myc-IPK1/ipk1-1* and *ITPK1-GFP/itpk1-2*, respectively) extraxts. The blots were probed (IB) with anti-CSN5 antibodies. **(D)** Size exclusion partitioning profile of CUL1 from Col-0, *ipk1-1*, and *itpk1-2* extracts. The blots were probed (IB) with anti-CUL1 antibodies. The migration positions of molecular weight standards (in kDa) are shown.

**Supplementary Figure 8.**
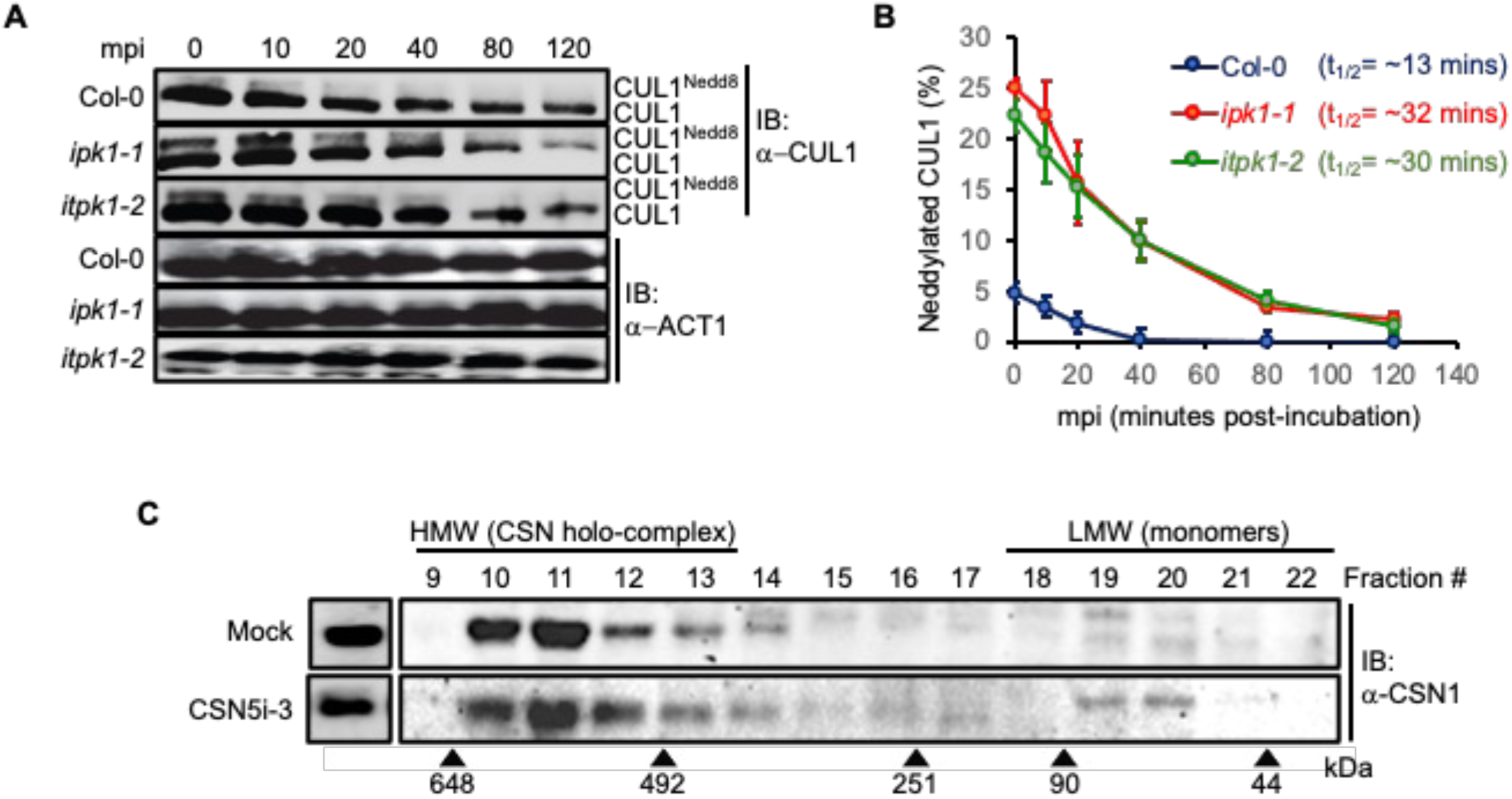
CUL1 Deneddylation rates are reduced in the InsP-kinase mutants. **(A)** *In lysate* deneddylation assay in Col-0, *ipk1-1*, and *itpk1-2* plants. Total extracts were incubated for indicated time points (mpi, minutes post-incubation) and probed (IB) with anti-CUL1 or anti-ACT1 antibodies. **(B)** Scatter plot showing half-life CUL1^Nedd8^ in the indicated plant extracts. **(C)** Size exclusion partitioning profile of CSN1 from control (mock) and CSN5i-3 treated Col-0 samples at 24 hours post-treatment (hpt). The migration positions of molecular weight standards (in kDa) are shown.

**Supplementary Figure 9.**
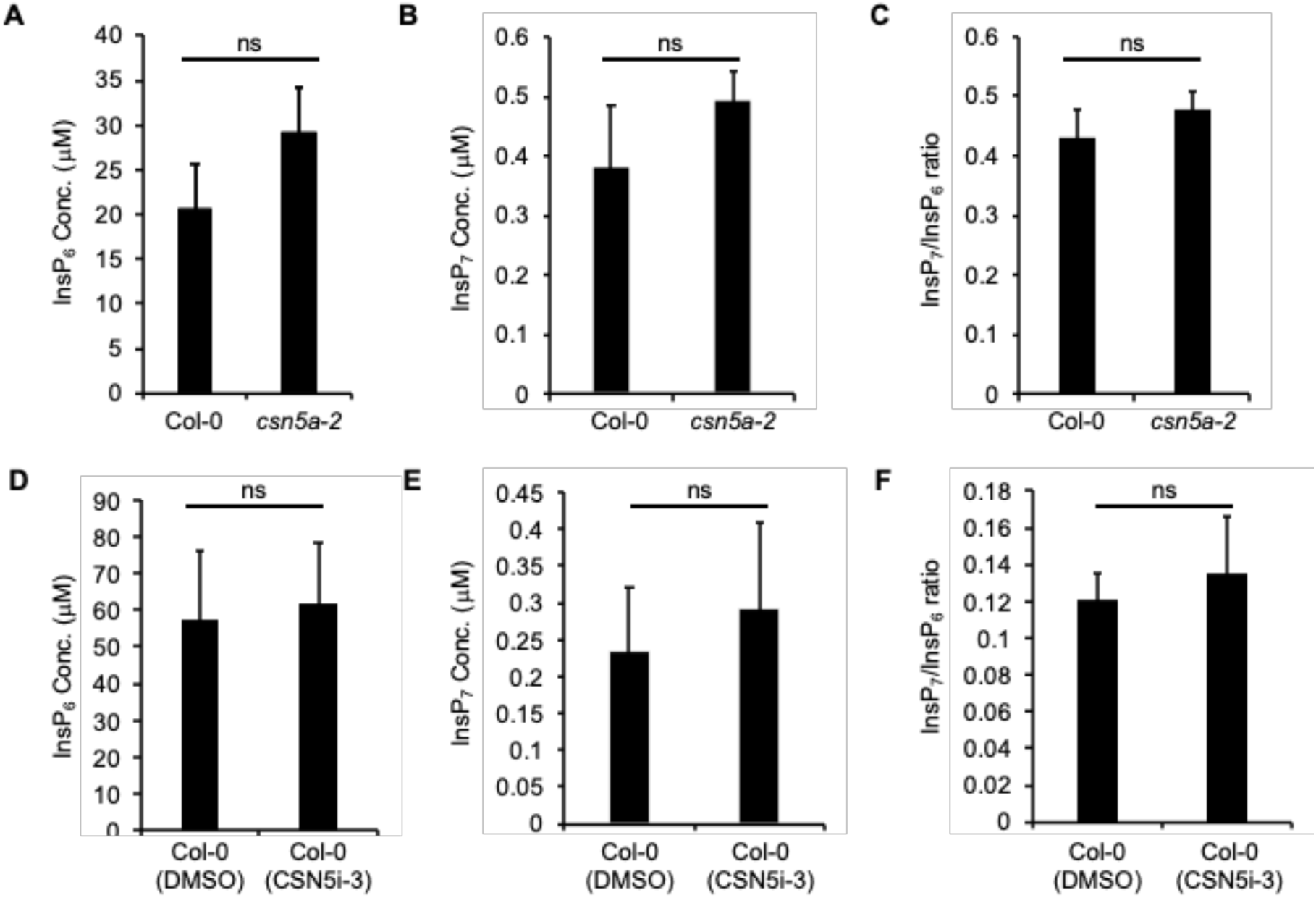
InsP_6/7_ levels are unaffected in *csn5a-2* plants or upon CSN5i-3 treatment on Col-0. **(A to C)** CE-ESI-MS analysis of InsP_6_, InsP_7_ levels and InsP_6_/InsP_7_ ratio in Col-0 and *csn5a-2* plants. **(D to F)** CE-ESI-MS analysis of InsP_6_, InsP_7_ levels and InsP_6_/InsP_7_ ratio in CSN5i-3-treated and untreated (DMSO) Col-0. Data shown here is means ± SE (n = 3). Statistical analysis is according to Student’s *t*-test (ns= non-significant).

**Supplementary Figure 10.**
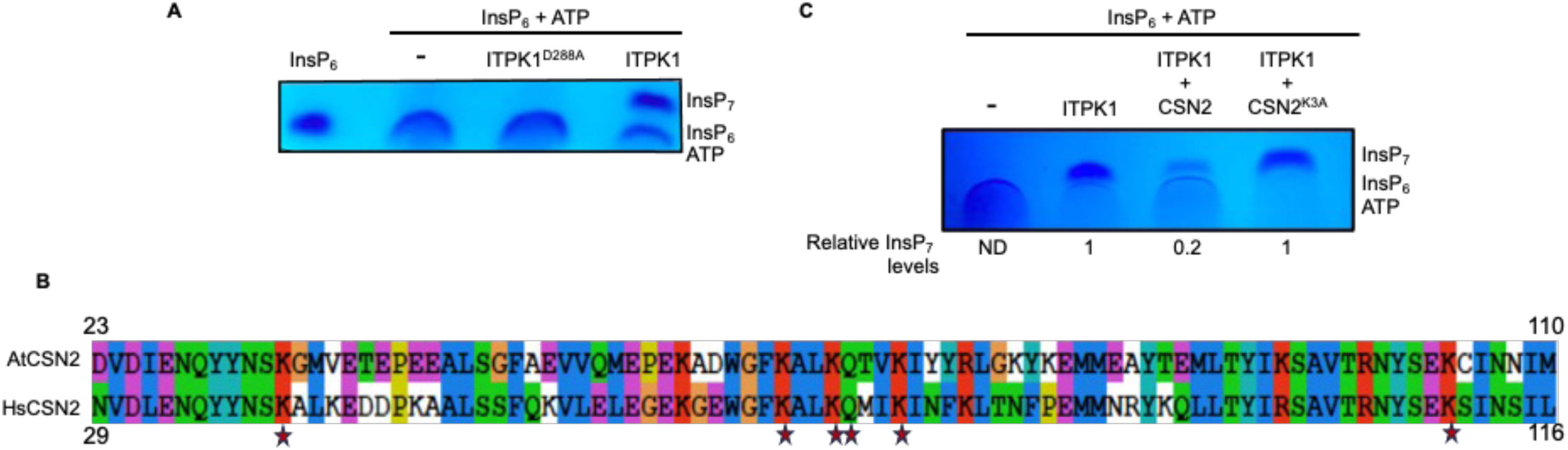
Competitive inhibition on ITPK1 kinase activity by CSN2, but not by CSN2^K3A^. **(A)** PAGE analysis of *in vitro* ITPK1 kinase reaction. **(B)** Alignment of AtCSN2 and HsCSN2 sequences. The InsP_6_-binding residues in HsCSN2 and conserved in AtCSN2 are marked with a star sign. **(C)** PAGE analysis of *in vitro* ITPK1 kinase reaction in the presence of wild-type and CSN2^K3A^ mutant. The numerical values below the gel image represent InsP_7_ levels relative to the ITPK1 alone reaction. The relative migration positions of InsP_7_, InsP_6_, and ATP are shown.

**Supplementary Figure 11.**
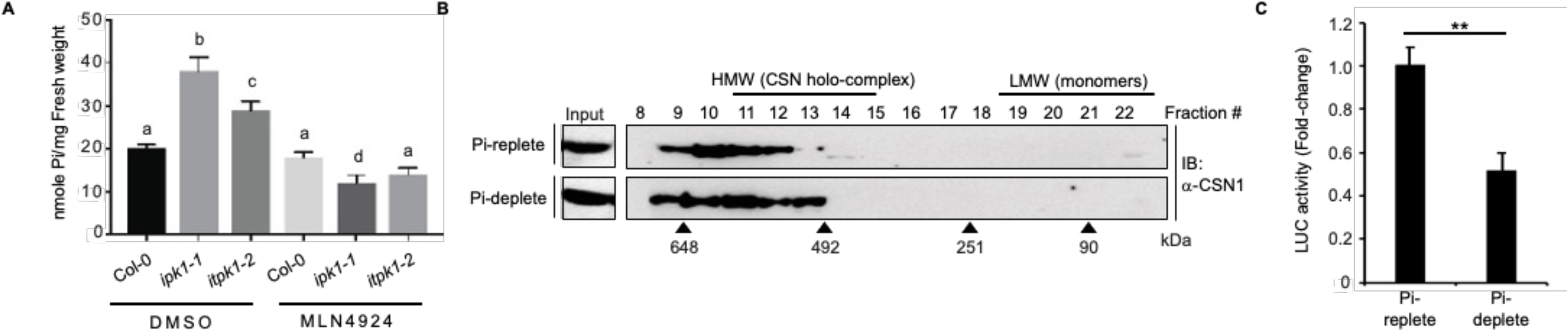
MLN4924 suppresses high endogenous Pi accumulation in the InsP kinase mutants. **(A)** Endogenous Pi levels in mock-(DMSO) or MLN4924-treated Col-0, *ipk1-1*, or *itpk1-2* plants. The data shown is mean ± SD (n=6-7). Different alphabets indicate significant differences (P< 0.05, ANOVA, Tukey) between samples. **(B)** CSN1 fractionation profile in Pi-replete or Pi-deplete Col-0 extracts detected via immunoblot (IB) with anti-CSN1 antibodies. Fraction numbers and elution positions of molecular weight standards (in kDa) are marked. CSN1 levels in the input extracts are shown. **(C)** Deneddylation activity of isolated CSN holo-complex from the indicated plants tested on Nedd8-AML substrate. LUC activity is presented as fold-change relative to Pi-replete data. Statistical analysis is according to Student’s *t*-test (**P<0.01).

**Supplementary Figure 12.**
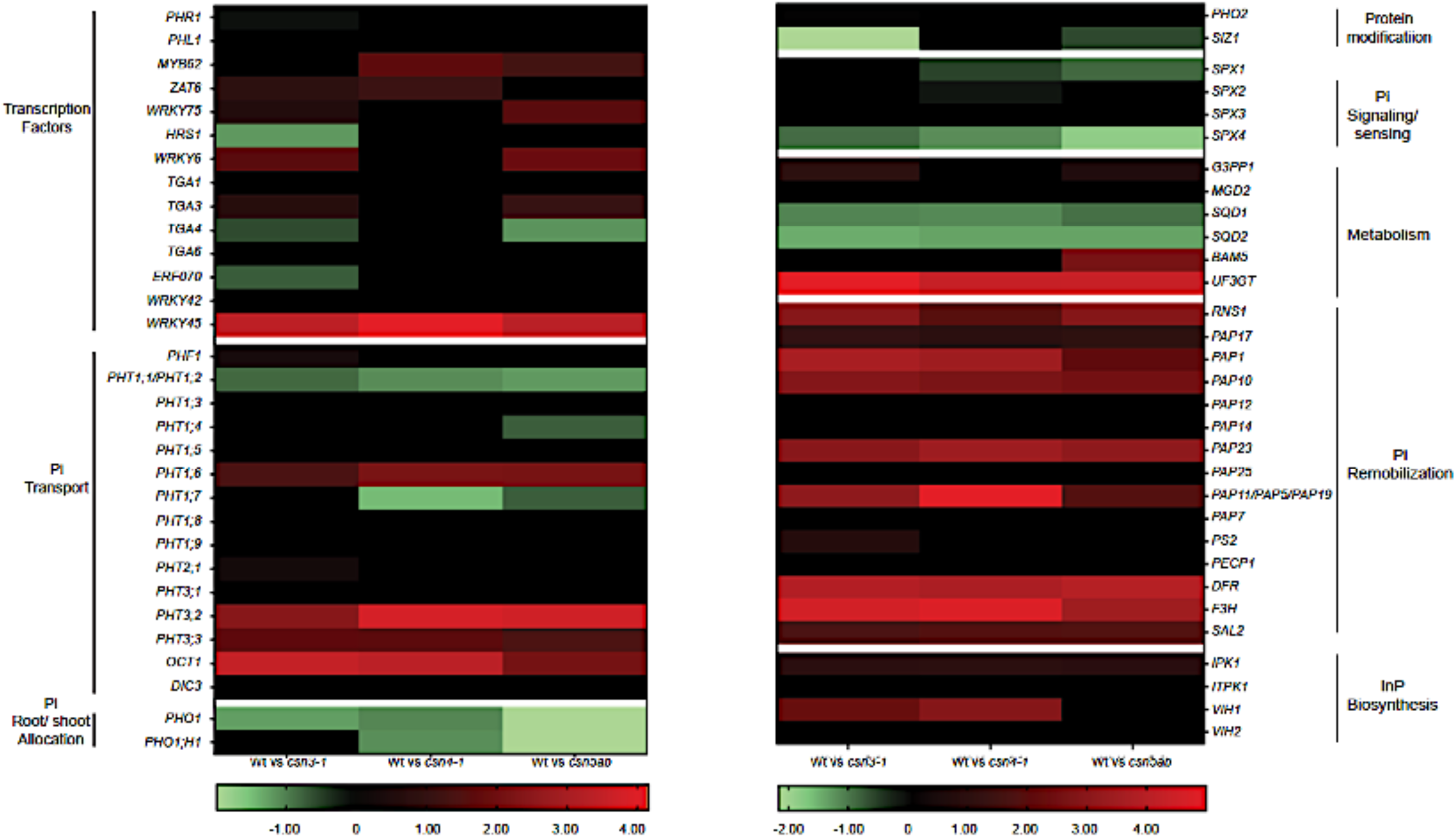
Differential expression of *PSI*-genes in *csn* mutants relative to wild-type plants (Wt). The data was curated from Dohmann *et al* (2008). The heatmap is generated using Graphpad Prism 8.4.3 using double gradient Heatmap function. The expression change is shown as log_2_FC (Fold-Change) values of different genes with Green for downregulated genes, Red for upregulated genes and black for either genes which show no change or the genes with expression values which are not significant (Adjusted *P* > 0.10). Functional categorizations are according to Kuo *et al* (2014).

**Supplementary Figure 13.**
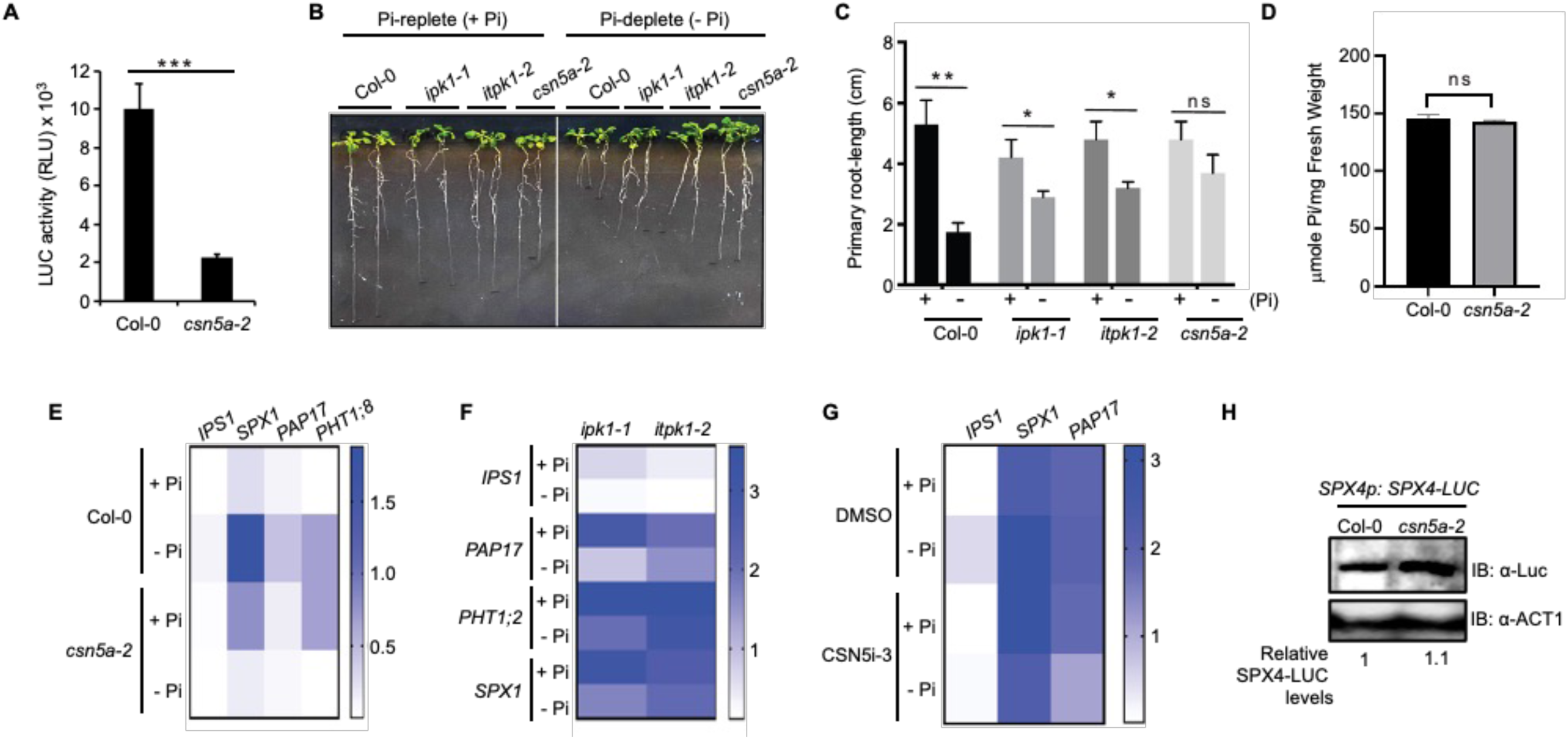
Functional inefficiency of CSN5 or InsP-kinases impair PSR. **(A)** Deneddylation activity of isolated CSN holo-complex from Col-0 versus *csn5a-2* plants tested on the Nedd8-AML substrate. **(B and C)** Primary root-growth inhibition under Pi-deplete (-Pi) versus Pi-replete (+Pi) conditions in Col-0, *csn5a-2*, i*pk1-1*, or *itpk1-2* plants. Data represents the mean (n=∼20) ± SD. **(C)** Endogenous Pi levels in Col-0 and *csn5a-2* plants. The data shown is mean ± SD (n=6-7). **(E to G)** Heatmaps of *PSI*-gene expressions in Col-0 versus *csn5a-2* **(E)**, *itpk1-1* and *itpk1-2* **(F)**, or mock (DMSO)- or CSN5i-3-treated Col-0 **(G)** under Pi-replete (+Pi) or Pi-deplete (-Pi) conditions (n=3). Gene expression were normalized to endogenous *MON1* transcripts. Heatmaps were generated using Graphpad Prism 8.4.3 software using the single gradient function. Data is indicative of mean values (± SD). Statistical analysis is by pairwise comparison with Student’s *t*-test (*P<0.05, **P< 0.01, ns= not significant). **(H)** SPX4-LUC levels in Col-0 versus *csn5a-2* plants. Total extracts were immunoblotted (IB) with anti-LUC antibodies. Anti-ACT1 immunoblot shows loading controls. The numbers below the image indicate relative SPX4-LUC amounts normalized to ACT1 levels.

**Supplemental Figure 14.**
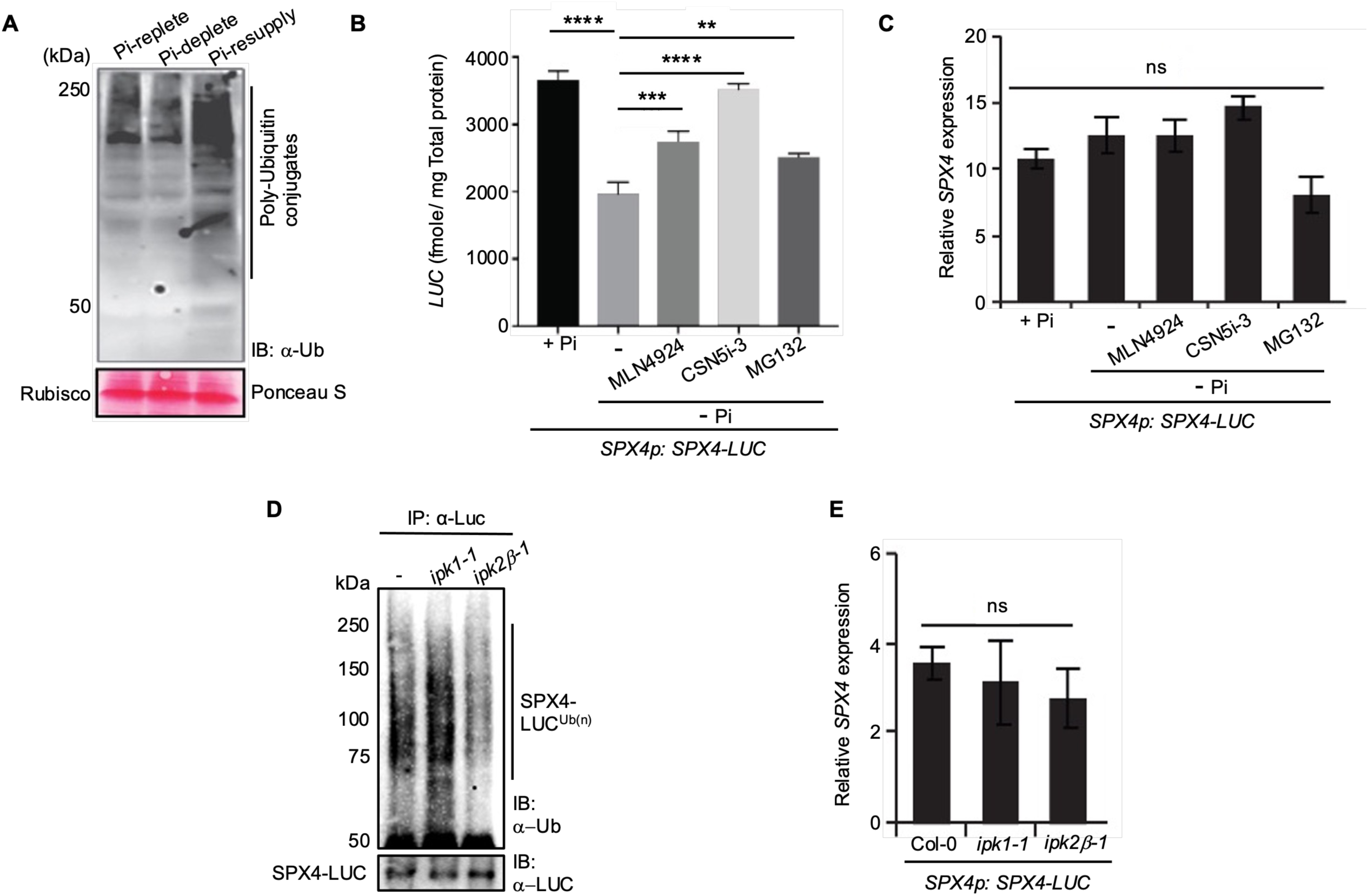
Pi-starvation causes CSN-mediated and CRL-dependent degradation of SPX4. **(A)** Global poly-ubiquitin conjugates in total extracts from Pi-replete, Pi-deplete, or Pi-deplete + resupply (Pi-resupply) Col-0 plants probed (IB) with anti-ubiquitin (Ub) antibodies. Approximate migration positions of poly-ubiquitin conjugated proteins are marked. Relative migration position of molecular weight standards (in kDa) are shown. Comparable total protein loading between the samples is denoted by Ponceau S staining. MLN4924 or CSN5i-3 reduce SPX4 degradation upon PSR. Luciferase (LUC) activity is indicative of SPX4-LUC levels under Pi-replete (+ Pi) and Pi-deplete (-Pi) conditions without or with indicated inhibitors. **(C-E)** Relative *SPX4* transcript levels in the indicated treatments **(C)**, or in the mentioned genotypes **(E)**. **(D)** Polyubiquitinated SPX4-LUC in control (-), *ipk1-1,* and *ipk2β-1* plants detected via immuno-enrichment (IP) with anti-LUC antibodies and probed (IB) with anti-Ub antibodies. Relative levels of unmodified SPX4-LUC are shown with anti-LUC immunoblot. Migration position of molecular weight standards (in kDa) are indicated. Gene expressions were normalized to endogenous *MON1* transcripts. Data represents mean ± SE (n=3). Statistical analysis is by pairwise comparison to + Pi **(B and C)**, or to Col-0 **(D)** with Student’s *t*-test (\*\**P*<0.01; \*\*\**P*< 0.001, \*\*\*\**P*<0.0001, ns= not significant).

## Notes

### Competing Interest Statement

The authors have declared no competing interest.

### Summary of Updates

New results and analysis has been added.

