## Supplemental Figures for "*Arabidopsis* inositol polyphosphate kinase activities regulate COP9 deneddylation functions in phosphate-homeostasis"

### Supplemental information

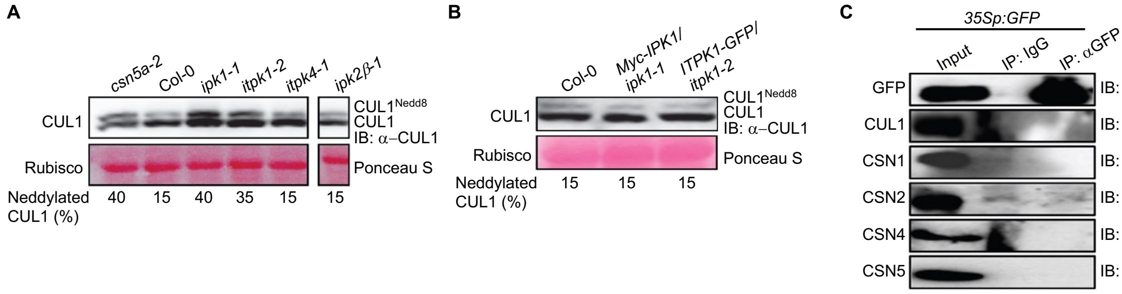


**Supplementary Figure 1. Increased CUL1$^{Nedd8}$: CUL1 ratios are specific to *ipk1-1* or *itpk1-2* but not other InsP-kinase mutants.**

**(A and B)** Anti-CUL1 immunoblot showing relative levels of neddylated: unneddylated CUL1 (CUL1$^{Nedd8}$: CUL1) in the indicated plant lines. Comparable protein loading between samples is shown by Ponceau S staining for rubisco large subunit. The numerical values below each lane represents the percentage of CUL1$^{Nedd8}$ relative to unneddylated CUL1 for the corresponding sample.

**(C)** Anti-GFP or anti-IgG-enrichments (IP) from control *35Sp:GFP*-expressing transgenic plants probed (IB) with the indicated antibodies. Relative protein amounts in the input extracts are also shown.

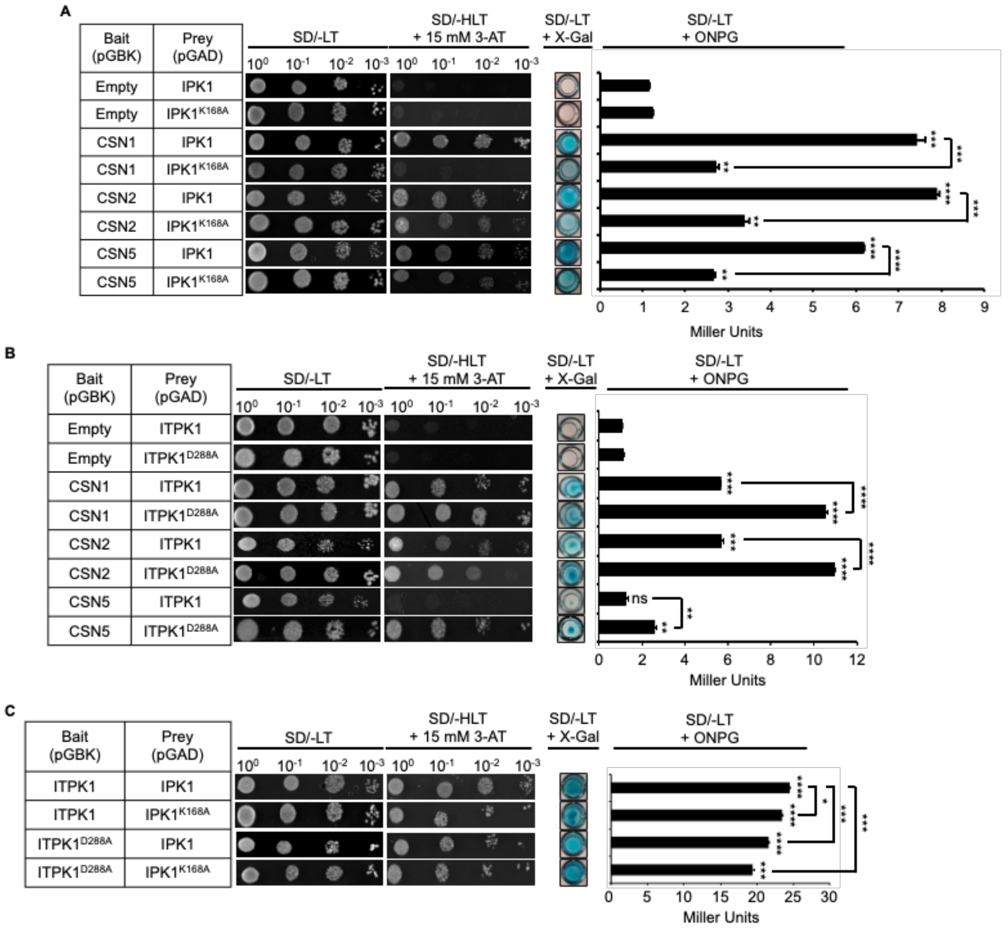


**Supplementary Figure 2. Kinase domains of IPK1 and ITPK1 affect their interaction with CSN subunits, but not with each other.**

**(A)** Yeast two-hybrid assays showing stronger interaction of kinase-proficient (IPK1) than kinase-deficient IPK1 ($IPK1^{K168A}$) with CSN1, CSN2, and CSN5 subunits.
**(B)** Yeast two-hybrid assays showing stronger interaction of kinase-deficient ($ITPK1^{D288A}$) than kinase-proficient ITPK1 (ITPK1) with CSN1, CSN2, and CSN5 subunits.
**(C)** Yeast two-hybrid assays showing mutual interaction of kinase-deficient and kinase-proficient IPK1 and ITPK1 (ITPK1) with each other.
Yeast transformants containing the indicated bait and prey combinations were plated with the indicated serial dilution on SD/-LT or SD/-HLT + 15 mM 3-AT plates, and the growth of cells was recorded after 3 days. Qualitative X-Gal assays were performed with permeabilized yeast transformants, and the images were acquired 12 hours post-incubation. Quantitative ONPG assays were performed with lysates from the indicated transformants, and color development was measured using a spectrophotometer (420 nm). For each assay, n=3-5.
Statistical analysis is according to Student's *t*-test: firstly, between indicated combinations of the tested InsP-kinase (wild-type or kinase-dead mutant) prey and empty or the indicated CSN subunit bait; secondly, between the indicated combinations of the wild-type InsP-kinase or its kinase-dead prey version with a tested CSN subunit bait; thirdly, between the indicated combinations of wild-type IPK1 and ITPK1 or their indicated kinase-dead version (*P<0.05, ***P*<0.01, ****P*<0.001, *****P*<0.0001, ns=not significant).

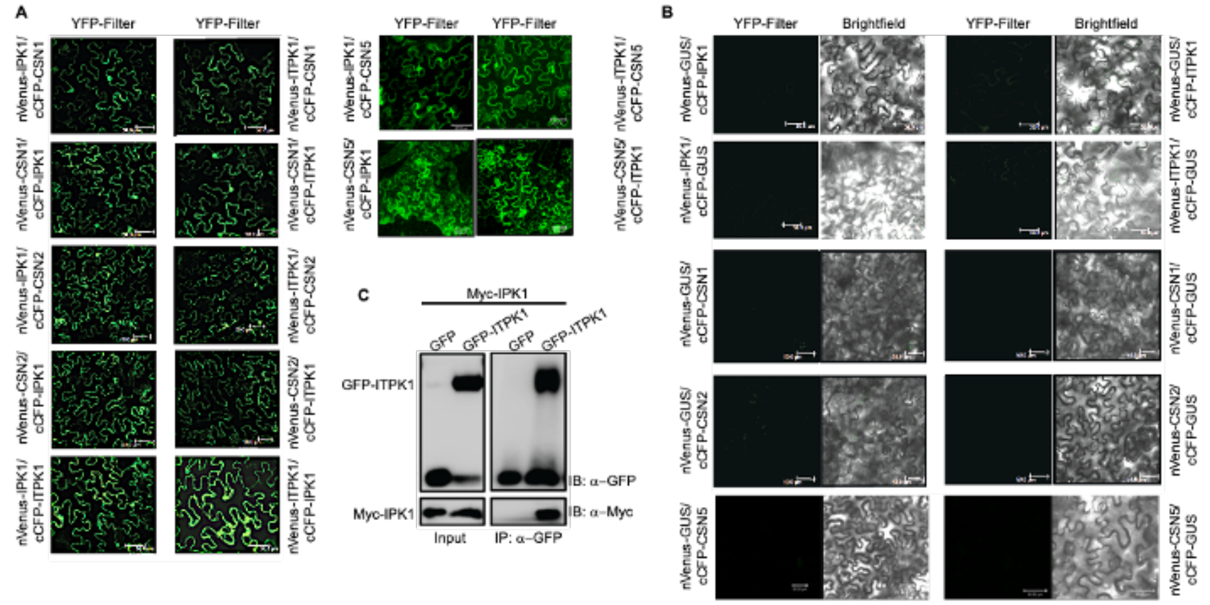


**Supplementary Figure 3.** ***In vivo*** **interactions between IPK1, ITPK1 and CSN subunits.**

**(A)** Confocal images of BiFC assays showing interactions between IPK1 and ITPK1 and with CSN1, CSN2, and CSN5 subunits.

**(B)** Lack of interactions between the negative control GUS with IPK1, ITPK1, CSN1, CSN2 or CSN5.

Images were acquired at 48 hpi (hours post-infiltration) with YFP and brightfield filters. Figures are representative of multiple images obtained for each BiFC combination. Scale bar = 50 μM.

**(C)** Co-immunoprecipitation of transiently expressed GFP-ITPK1 with Myc-IPK1. Anti-GFP pulldown (IP) from GFP alone or GFP-ITPK1 co-expressed with Myc-IPK1 were probed (IB) with anti-GFP or anti-Myc antibodies. Protein amounts in the input samples are indicated.

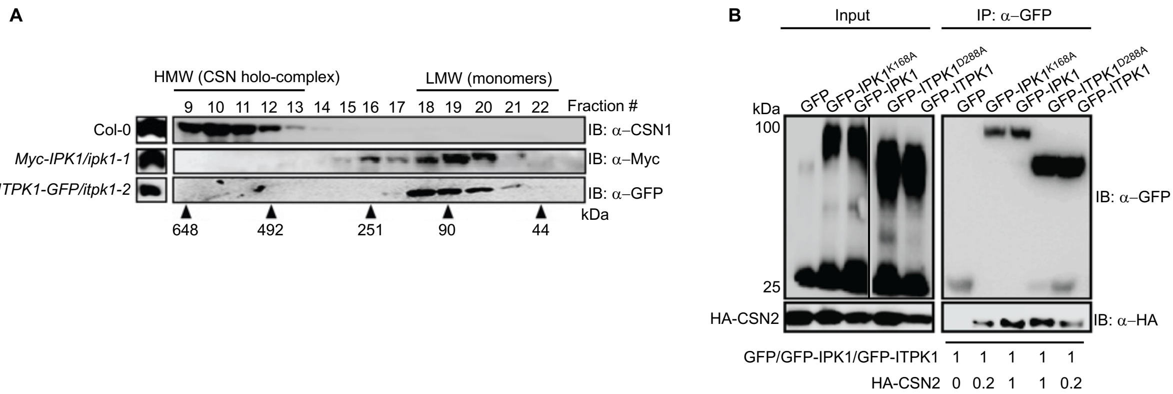


**Supplementary Figure 4. CSN2 associations of IPK1 and ITPK1 are conversely affected by their catalytic activities.**

**(A)** Gel-filtration fractionation pattern of Myc-IPK1 or ITPK1-GFP probed (IB) with the indicated antibodies. Fraction numbers and elution positions of molecular weight standards (in kDa) are indicated. CSN1 partitioning profile was used as a reference for the elution pattern of the CSN holo-complex.

**(B)** Immuno-enrichment of GFP-tagged wild-type or kinase-dead IPK1$^{K168A}$ or ITPK1$^{D288A}$ co-expressed with HA-CSN2 in *N. benthamiana* leaves. Pull-down (IP) was performed with anti-GFP antibodies and the bound proteins were probed (IB) with anti-GFP or anti-HA antibodies. The migration position of molecular weight standards (in kDa) and the total protein amounts in the extracts (input) are shown. The numerical values below the IP image represents the relative CSN2 amounts normalized to the corresponding enriched GFP-tagged InsP-kinase.

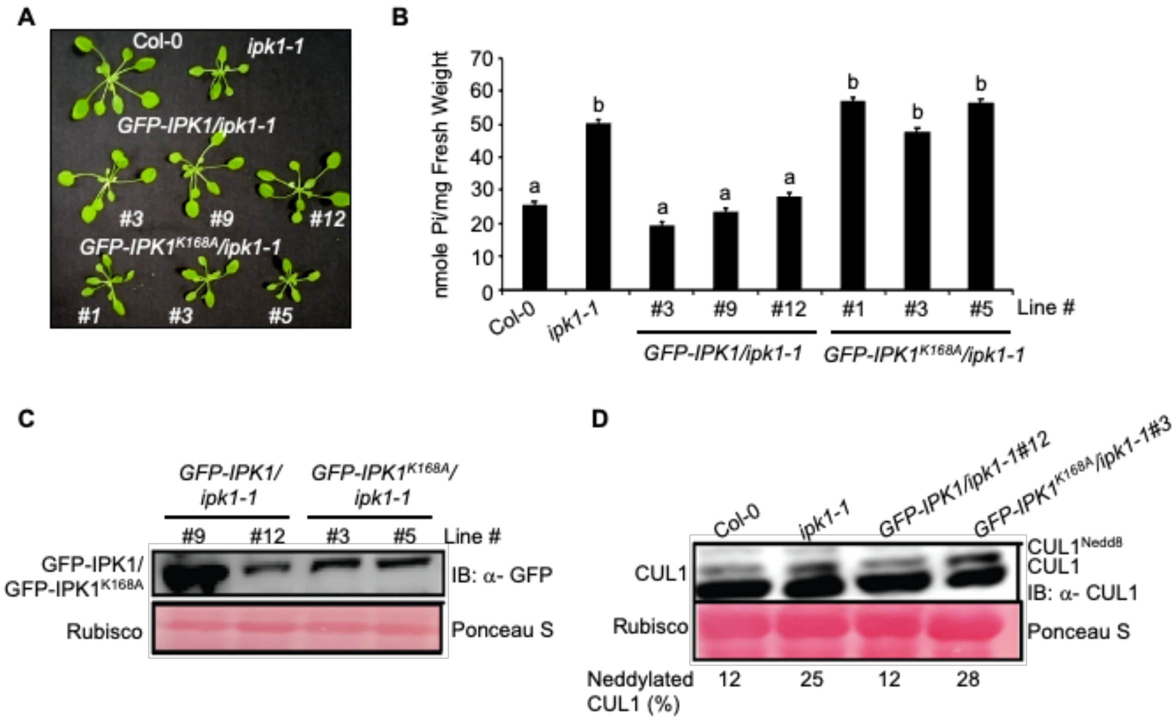


**Supplementary Figure 5. Kinase-proficient, but not kinase-deficient IPK1, restore *ipk1-1* defects.**

**(A)** Growth phenotypes of independent transgenic lines of *ipk1-1* plants expressing *GFP-IPK1* or *GFP-IPK1*$^{K168A}$.

**(B)** Endogenous Pi levels in the indicated genotypes. The data shown is mean ± SD (n=6-7). Different alphabets indicate significant differences ($P< 0.05$, ANOVA, Tukey) between samples.

**(C)** Expression levels of GFP-IPK1 or GFP-IPK1$^{K168A}$ in the indicated transgenic lines probed with anti-GFP antibodies.

**(D)** Relative CUL1$^{Nedd8}$:CUL1 levels in the indicted plant lines probed (IB) with anti-CUL1 antibodies. Relative migration positions of CUL1$^{Nedd8}$ and CUL1 are marked. The numerical values below the image represent the percentage of CUL1$^{Nedd8}$ relative to unneddylated CUL1 for the corresponding sample.

Ponceau S staining shows loading controls.

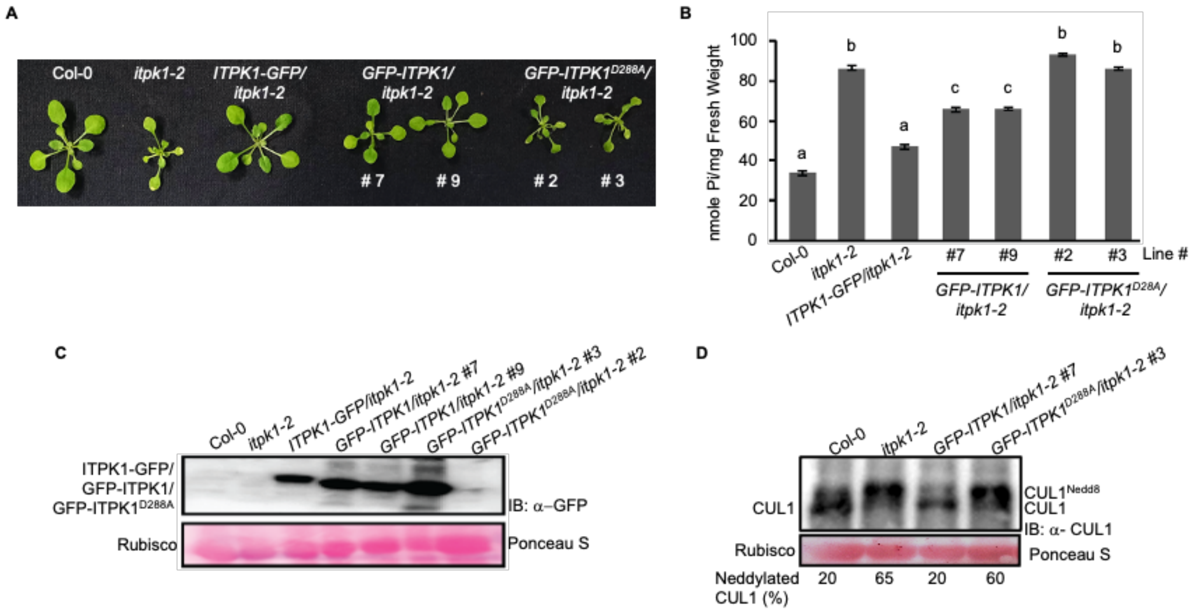


**Supplementary Figure 6. Kinase-proficient, but not kinase-deficient ITPK1, restore *itpk1-2* defects.**

**(A)** Growth phenotypes of independent transgenic lines of *itpk1-2* plants expressing *GFP-ITPK1* or *GFP-ITPK1$^{D288A}$*.

**(B)** Endogenous Pi levels in the indicated genotypes. The data shown is mean ± SD (n=6-7). Different alphabets indicate significant differences ($P < 0.05$, ANOVA, Tukey) between samples.

**(C)** Expression levels of GFP-ITPK1 or GFP-ITPK1$^{D288A}$ in the indicated transgenic lines probed with anti-GFP antibodies.

**(D)** Relative CUL1$^{Nedd8}$:CUL1 levels in the indicted plant lines probed (IB) with anti-CUL1 antibodies. The migration positions of CUL1$^{Nedd8}$ and CUL1 are marked. The numerical values below the image represent the percentage of CUL1$^{Nedd8}$ relative to unneddylated CUL1 for the corresponding sample.

Ponceau S staining shows loading controls.

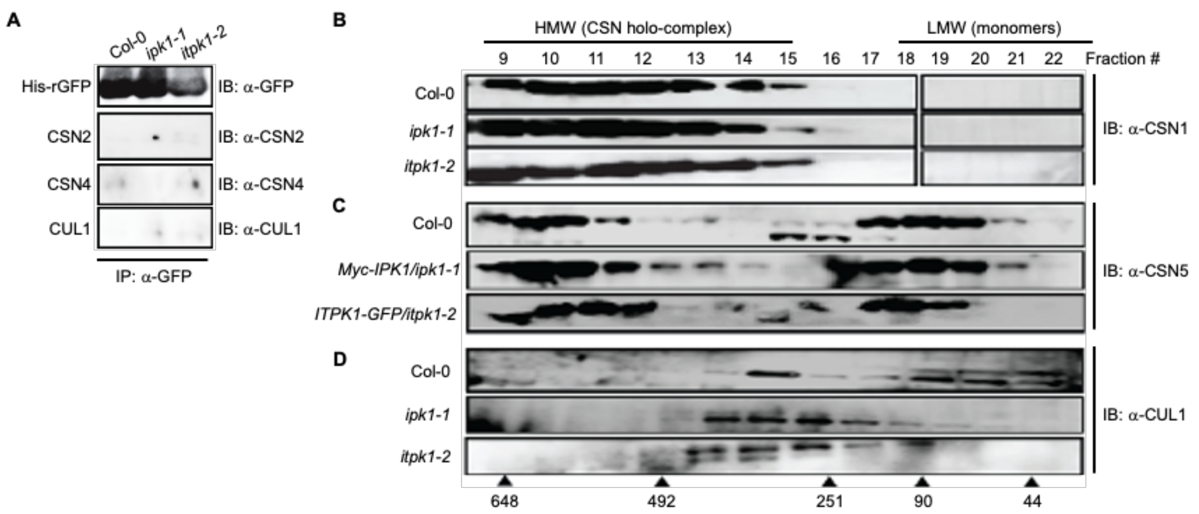


**Supplementary Figure 7. Fractionation profiles of CSN subunits and CUL1 in the indicated plants.**

**(A)** Control His-rGFP enrichments (IP) from supplemented extracts of indicated plants probed (IB) with the various antibodies.

**(B)** Size exclusion partitioning profile of CSN1 from Col-0, *ipk1-1*, and *itpk1-2* extracts. The blots were probed (IB) with anti-CSN1 antibodies.

**(C)** Size exclusion partitioning profile of CSN5 Col-0, and the complemented lines of *ipk1-1* and *itpk1-2* (*Myc-IPK1/ipk1-1* and *ITPK1-GFP/itpk1-2*, respectively) extraxts. The blots were probed (IB) with anti-CSN5 antibodies.

**(D)** Size exclusion partitioning profile of CUL1 from Col-0, *ipk1-1*, and *itpk1-2* extracts. The blots were probed (IB) with anti-CUL1 antibodies.

The migration positions of molecular weight standards (in kDa) are shown.

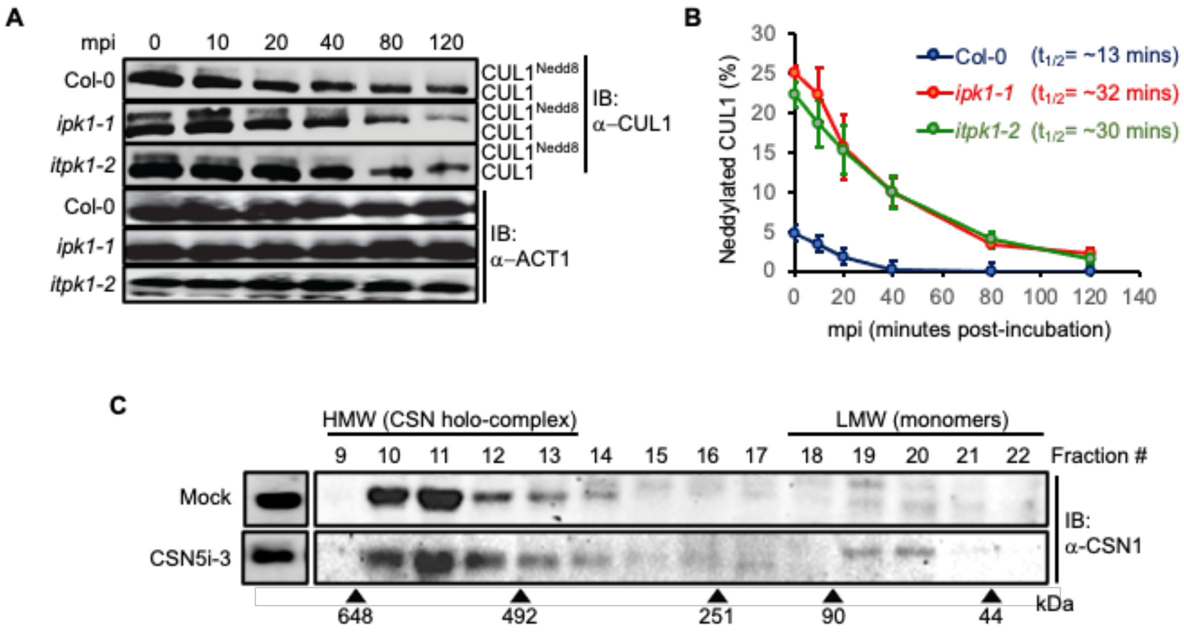


**Supplementary Figure 8. CUL1 Deneddylation rates are reduced in the InsP-kinase mutants.**

**(A)** *In lysate* deneddylation assay in Col-0, *ipk1-1*, and *itpk1-2* plants. Total extracts were incubated for indicated time points (mpi, minutes post-incubation) and probed (IB) with anti-CUL1 or anti-ACT1 antibodies.

**(B)** Scatter plot showing half-life $CUL1^{Nedd8}$ in the indicated plant extracts.

**(C)** Size exclusion partitioning profile of CSN1 from control (mock) and CSN5i-3 treated Col-0 samples at 24 hours post-treatment (hpt). The migration positions of molecular weight standards (in kDa) are shown.

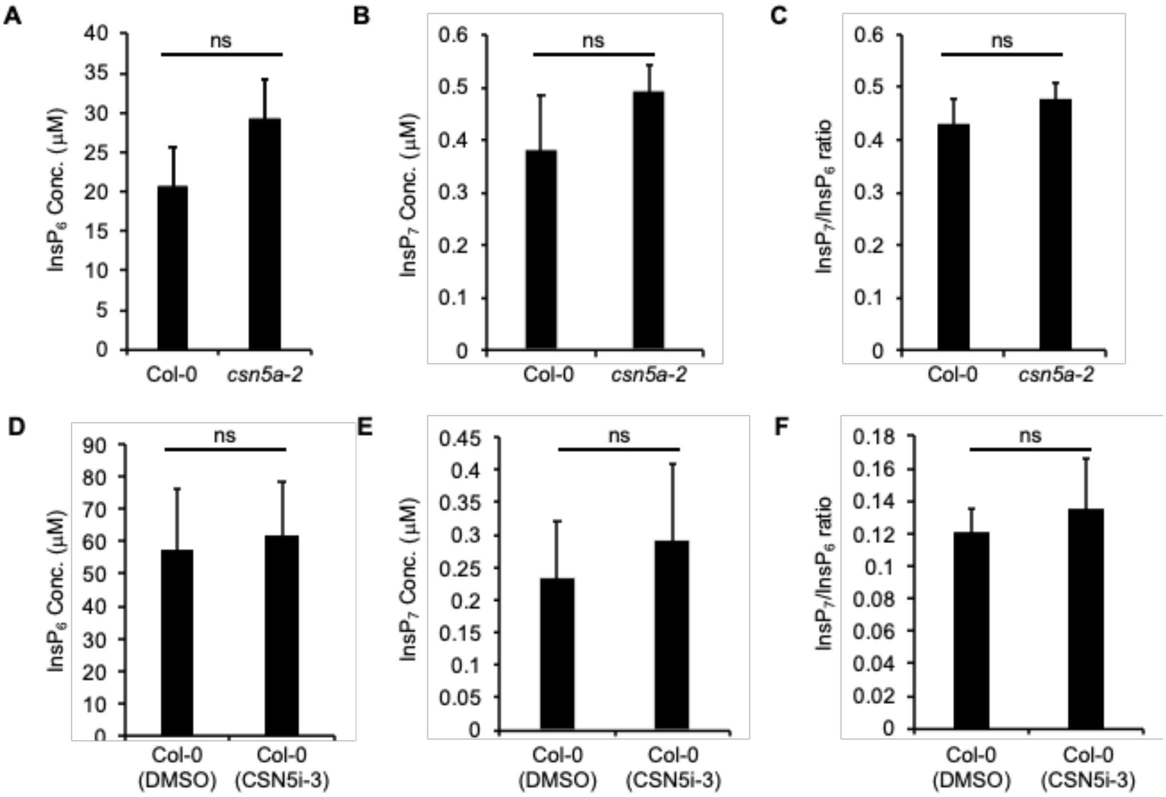


**Supplementary Figure 9. $InsP_{6/7}$ levels are unaffected in *csn5a-2* plants or upon CSN5i-3 treatment on Col-0.**

**(A to C)** CE-ESI-MS analysis of $InsP_6$, $InsP_7$ levels and $InsP_6/InsP_7$ ratio in Col-0 and *csn5a-2* plants.

**(D to F)** CE-ESI-MS analysis of $InsP_6$, $InsP_7$ levels and $InsP_6/InsP_7$ ratio in CSN5i-3-treated and untreated (DMSO) Col-0.

Data shown here is means ± SE (n = 3). Statistical analysis is according to Student's *t*-test (ns= non-significant).

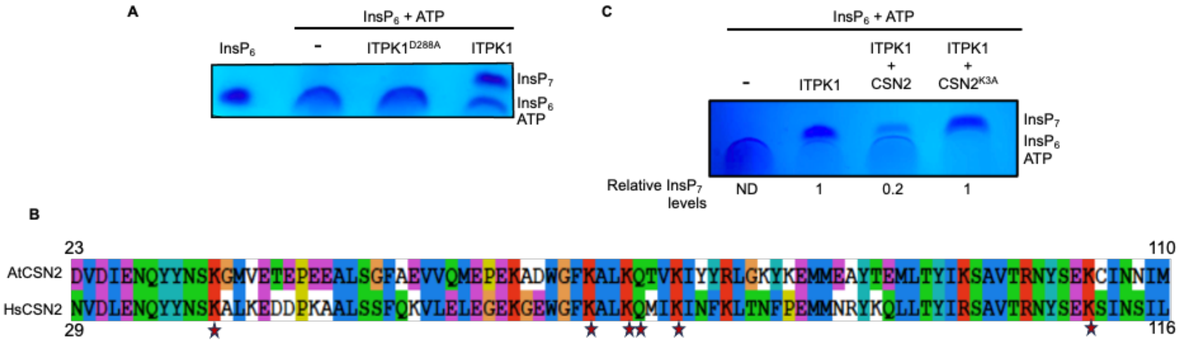


**Supplementary Figure 10. Competitive inhibition on ITPK1 kinase activity by CSN2, but not by CSN2$^{K3A}$.**

**(A)** PAGE analysis of *in vitro* ITPK1 kinase reaction.

**(B)** Alignment of AtCSN2 and HsCSN2 partial sequences. The $InsP_6$-binding residues in HsCSN2 and conserved in AtCSN2 are marked with a star sign.

**(C)** PAGE analysis of *in vitro* ITPK1 kinase reaction in the presence of wild-type and CSN2$^{K3A}$ mutant. The numerical values below the gel image represent $InsP_7$ levels relative to the ITPK1 alone reaction.

The relative migration positions of $InsP_7$, $InsP_6$, and ATP in PAGE are shown.

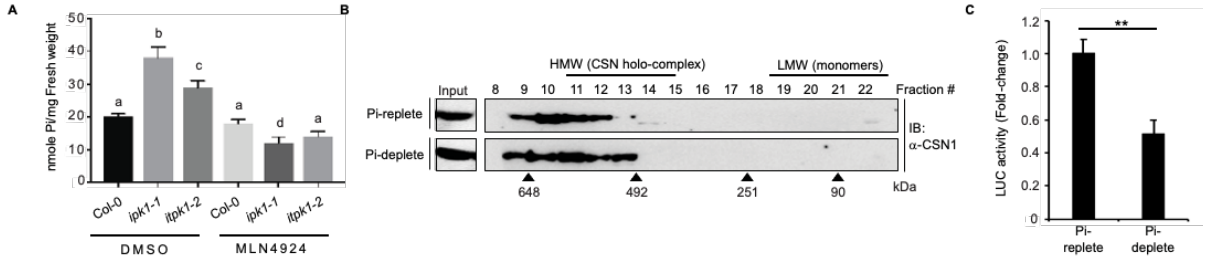


**Supplementary Figure 11. MLN4924 suppresses high endogenous Pi accumulation in the InsP kinase mutants.**

**(A)** Endogenous Pi levels in mock-(DMSO) or MLN4924-treated Col-0, *ipk1-1*, or *itpk1-2* plants. The data shown is mean ± SD (n=6-7). Different alphabets indicate significant differences (P< 0.05, ANOVA, Tukey) between samples.

**(B)** CSN1 fractionation profile in Pi-replete or Pi-deplete Col-0 extracts detected via immunoblot (IB) with anti-CSN1 antibodies. Fraction numbers and elution positions of molecular weight standards (in kDa) are marked. CSN1 levels in the input extracts are shown.

**(C)** Deneddylation activity of isolated CSN holo-complex from the indicated plants tested on Nedd8-AML substrate. LUC activity is presented as fold-change relative to Pi-replete data. Statistical analysis is according to Student's *t*-test (**P<0.01).

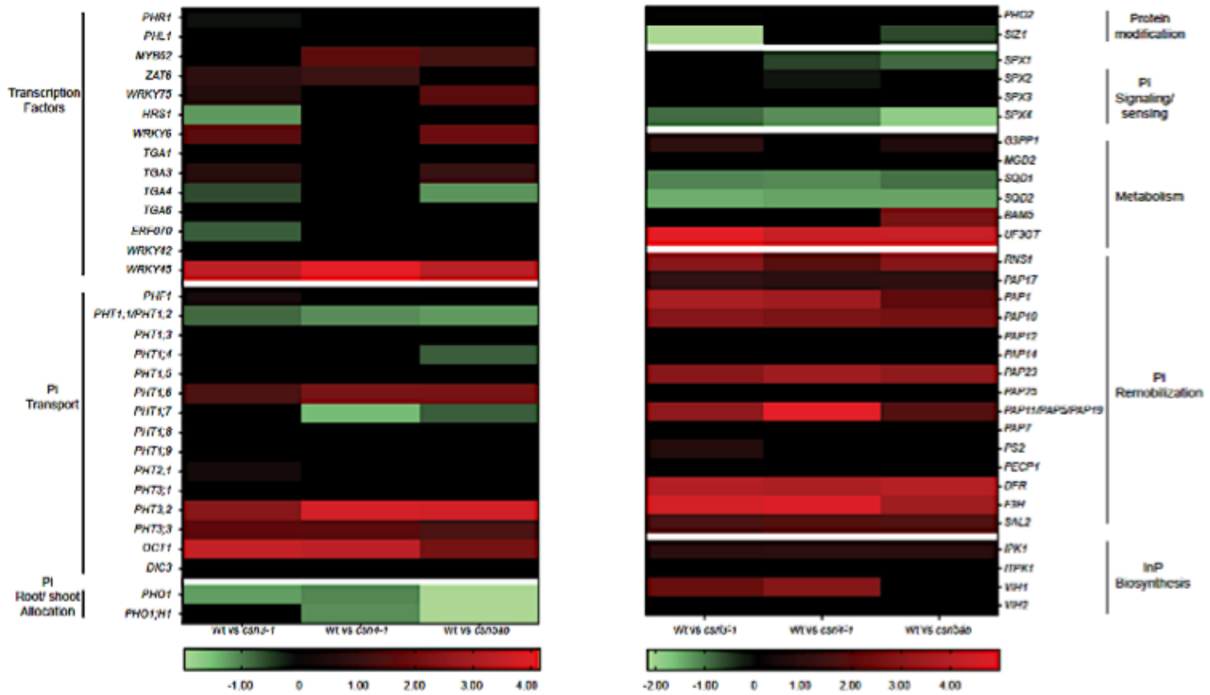


**Supplementary Figure 12. Differential expression of *PSI*-genes in *csn* mutants relative to wild-type plants (Wt).**

The data was curated from Dohmann *et al* (2008). The heatmap is generated using Graphpad Prism 8.4.3 using the double gradient Heatmap function. The expression change is shown as $log_2FC$ (Fold-Change) values of different genes, with Green for downregulated genes, Red for upregulated genes and black for either genes that show no change or the genes with expression values that are not significant (Adjusted $P > 0.10$). Functional categorizations are according to Kuo *et al* (2014).

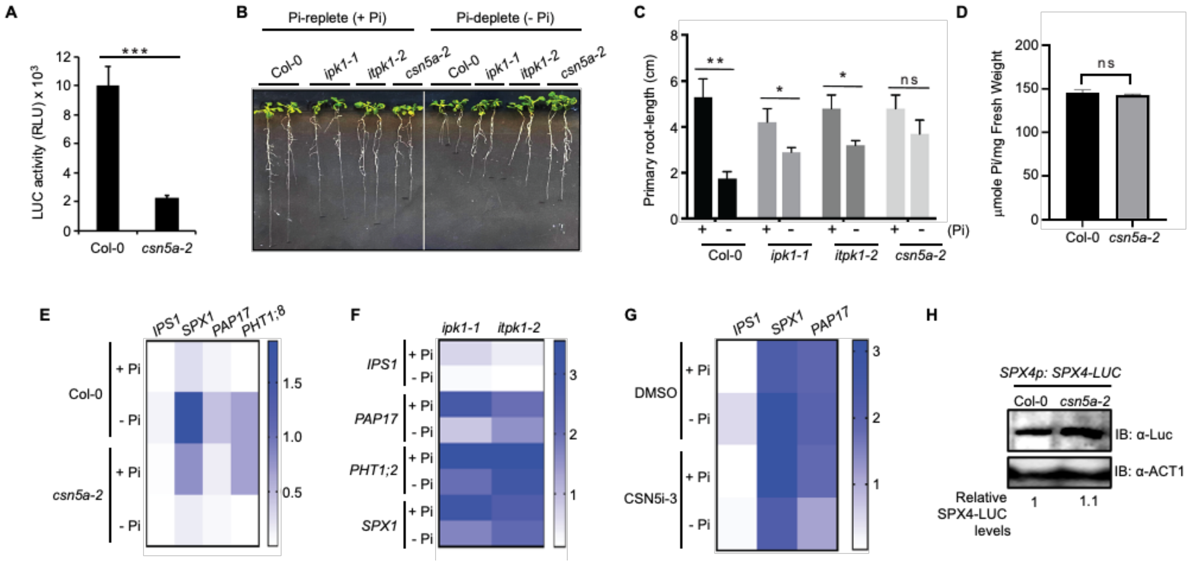


**Supplementary Figure 13. Functional inefficiency of CSN5 or InsP-kinases impair PSR.**

**(A)** Deneddylation activity of isolated CSN holo-complex from Col-0 versus *csn5a-2* plants tested on the Nedd8-AML substrate.

**(B and C)** Primary root-growth inhibition under Pi-deplete (-Pi) versus Pi-replete (+Pi) conditions in Col-0, *csn5a-2*, i*pk1-1*, or *itpk1-2* plants. Data represents the mean (n=~20) ± SD.

**(D)** Endogenous Pi levels in Col-0 and *csn5a-2* plants. The data shown is mean ± SD (n=6-7).

**(E to G)** Heatmaps of *PSI*-gene expressions in Col-0 versus *csn5a-2* **(E)**, *itpk1-1* and *itpk1-2* **(F)**, or mock (DMSO)- or CSN5i-3-treated Col-0 **(G)** under Pi-replete (+Pi) or Pi-deplete (-Pi) conditions (n=3). Gene expression were normalized to endogenous *MON1* transcripts. Heatmaps were generated using GraphPad Prism 8.4.3 software using the single gradient function.

Data is indicative of mean values (± SD). Statistical analysis is by pairwise comparison with Student's *t*-test (*P<0.05, **P< 0.01, ns= not significant).

**(H)** SPX4-LUC levels in Col-0 versus *csn5a-2* plants. Total extracts were immunoblotted (IB) with anti-LUC antibodies. Anti-ACT1 immunoblot shows loading controls. The numbers below the image indicate relative SPX4-LUC amounts normalized to ACT1 levels.

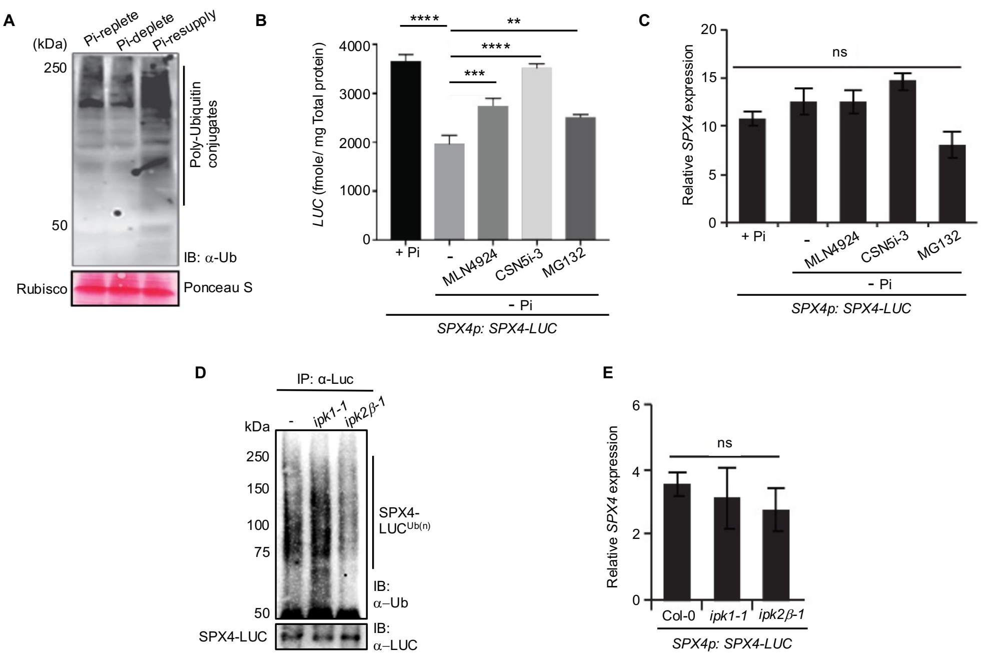


**Supplemental Figure 14. Pi-starvation causes CSN-mediated and CRL-dependent degradation of SPX4.**

**(A)** Global poly-ubiquitin conjugates in total extracts from Pi-replete, Pi-deplete, or Pi-deplete + resupply (Pi-resupply) Col-0 plants probed (IB) with anti-ubiquitin (Ub) antibodies. Approximate migration positions of poly-ubiquitin conjugated proteins are marked. Relative migration position of molecular weight standards (in kDa) are shown. Comparable total protein loading between the samples is denoted by Ponceau S staining.

**(B)** MLN4924 or CSN5i-3 reduce SPX4 degradation upon PSR. Luciferase (LUC) activity is indicative of SPX4-LUC levels under Pi-replete (+ Pi) and Pi-deplete (-Pi) conditions without or with indicated inhibitors.

**(C-E)** Relative SPX4 transcript levels in the indicated treatments (C), or in the mentioned genotypes **(E)**.

**(D)** Polyubiquitinated SPX4-LUC in control (-), *ipk1-1*, and *ipk2β-1* plants detected via immuno-enrichment (IP) with anti-LUC antibodies and probed (IB) with anti-Ub antibodies. Relative levels of unmodified SPX4-LUC are shown with anti-LUC immunoblot. Migration position of molecular weight standards (in kDa) are indicated.

Gene expressions were normalized to endogenous *MON1* transcripts. Data represents mean ± SE (n=3). Statistical analysis is by pairwise comparison to + Pi (B and C), or to Col-0 (D) with Student's *t*-test (***P*<0.01; ****P*< 0.001, *****P*<0.0001, ns= not significant).

**Table S1.** List of oligonucleotide primers used in this study.

| S. No | Primer Name | Primer Sequence (5' to 3') | Purpose |
|---|---|---|---|
| 1. | attB1 *IPK1* For | GGGGACAAGTTTGTACAAAAAAGCAG GCTCAATGGAGATGATTTTGGAGG | Gateway cloning of IPK1 cDNA in *p*DONR207 |
| 2. | attB2 *IPK1* Rev | GGGGACCACTTTGTACAAGAAAGCTG GGTATTAGCTGTGGGAAGGTTTTGAG | Gateway cloning of IPK1 cDNA in *p*DONR207 |
| 3. | attB1 *ITPK1* For | GGGGACAAGTTTGTACAAAAAAGCAG GCTCAATGTCAGATTCAATCCAG | Gateway cloning of ITPK1 cDNA in *p*DONR207 |
| 4. | attB1 *ITPK1* Rev | GGGGACCACTTTGTACAAGAAAGCTG GGTATCAGACATGATTCTTCTTAGTG | Gateway cloning of ITPK1 cDNA in *p*DONR207 |
| 5. | attB1 *CSN1* For | GGGGACAAGTTTGTACAAAAAAGCAG GCTCAATGGAGCGAGACGAAGAAGCG AGT | Gateway cloning of CSN1 cDNA in *p*DONR207 |
| 6. | attB2 *CSN1* Rev | GGGGACCACTTTGTACAAGAAAGCTG GGTATCAAAGTTTCCTTGCCGATCTGG CATG | Gateway cloning of CSN1 cDNA in *p*DONR207 |
| 7. | attB1 *CSN2* For | GGGGACAAGTTTGTACAAAAAAGCAG GCTCAATGGCTTCAGATGCTGACATG GAGGAC | Gateway cloning of CSN2 cDNA in *p*DONR207 |
| 8. | attB2 *CSN2* Rev | GGGGACCACTTTGTACAAGAAAGCTG GGTATTAACACAGTCGGCTGGTGATA TTTGA | Gateway cloning of CSN2 cDNA in *p*DONR207 |
| 9. | qRT-PCR *IPS1* For | GCGTTTTAAGATATGGAGCAATG | For qRT-PCRs |
| 10. | attb1 CSN5 For | AAAAAGCAGGCTCAATGGAAGGTTCC TCCTCAGCC | Gateway cloning of CSN5 cDNA in *p*DONR207 |
| 11. | attb2 CSN5 Rev | AGAAAGCTGGGTACGATGTAATCATG GGCTCTGGATCTGATG | Gateway cloning of CSN5 cDNA in *p*DONR207 |
| 12. | *IPK1* K168A For | GGTGGTGATTGCATTAGTGTTGAAATA GCGCCCAAATGCGGATT | For generating $\mathrm{IPK1}^{K168A}$ clones |
| 13. | *IPK1* K168A Rev | ATCCGCATTTGGGCGCTATTTCAACAC TAATGCAATCACCACC | For generating $\mathrm{IPK1}^{K168A}$ clones |
| 14. | *ITPK1* D288A For | ATGCTAATAGGTACCTTATAATTGCTA TTAACTACTTTCCTGGATATGC | For generating $\mathrm{ITPK1}^{D288A}$ clones |
| 15. | *ITPK1* D288A Rev | GCATATCCAGGAAAGTAGTTAATAGC AATTATAAGGTACCTATTAGCAT | For generating $\mathrm{ITPK1}^{D288A}$ clones |

| | | | |
|---|---|---|---|
| 16. | CSN2 K3A For | AACCTGAGAAAGCTGACTGGGGTTTC GCAGCTCTTGCGGCGACTGTGAAGAT CTATTATCGTCTAG | For generating CSN2$^{K3A}$ clones |
| 17. | CSN2 K3A Rev | CTAGACGATAATAGATCTTCACAGTC GCCGCAAGAGCTGCGAAACCCCAGTC AGCTTTCTCAGGTT | For generating CSN2$^{K3A}$ clones |
| 18. | *p*DONR 207 For | CGCGTTAACGCTAGCATGGAT | For cloning into *p*DONR 207 |
| 19. | *p*DONR 207 Rev | CCATACAAGCGATAGATTGTCGCAC | For cloning into *p*DONR 207 |
| 20. | qRT-PCR *IPS1* Rev | CGAAGCTTGCCAAAGGATAG | For qRT-PCRs |
| 21. | qRT-PCR *SPX1* For | GATTCCATTGTTGGAGCAAGA | For qRT-PCRs |
| 22. | qRT-PCR *SPX1* Rev | AATCTTGTTAGCTTCTTCTATTGTA | For qRT-PCRs |
| 23. | qRT-PCR *PHT1;3* For | AAACATTGTGAACCATGCTAACCCTT | For qRT-PCRs |
| 24. | qRT-PCR *PHT1;3* Rev | GATAAACAAAATTGATTGGAATGACC ACTA | For qRT-PCRs |
| 25. | qRT-PCR *PHT1;8* For | GTCTGAAGATGAGCCACAGATT | For qRT-PCRs |
| 26. | qRT-PCR *PHT1;8* Rev | TCGTGTCTGAAGCAAAGTGCT | For qRT-PCRs |
| 27. | qRT-PCR *PAP17* For | AGAGCCGGCTAAAAGCGAC | For qRT-PCRs |
| 28. | qRT-PCR *PAP17* Rev | ATCTCCCGTAGACACCACGA | For qRT-PCRs |
| 29. | qRT-PCR *MON1* For | AACTCTATGCAGCATTTGATCCACT | For qRT-PCRs |
| 30. | qRT-PCR *MON1* Rev | TGATTGCATATCTTTATCGCCATC | For qRT-PCRs |
| 31. | *ipk1-1* RP | TTTCAATCAAGGAATGCTTGG | Genotyping *ipk1-1* |
| 32. | *ipk1-1* LP | ATGCTACGGGAAACCATTTTC | Genotyping *ipk1-1* |
| 33. | *itpk1-2* RP | CCATGTCCCAGAAGAACTCAG | Genotyping *itpk1-2* |
| 34. | *itpk1-2* LP | ACCAATATTCGATTCGATTCCACACG | Genotyping *itpk1-2* |
| 35. | LB1.3 | ATTTTGCCGATTTCGGAAC | T-DNA Left-border primer (for *ipk1-1*) |

| | | | |
|---|---|---|---|
| 36. | LB3 | TAGCATCTGAATTTCATAACCAATCTC GATACAC | T-DNA Left-border primer (for *itpk1-2*) |
| 37. | AJHIFW2 | GTTTTGGATTAGCATTAGTCCCCAAAT C | Genotyping *csn5a-2* plants (Dohmann *et al*., 2005) |
| 38. | AJH1RV2 | TTCAAACATAAATGTGAAAAACAACA T | Genotyping *csn5a-2* plants (Dohmann *et al*., 2005) |
| 39. | LBb1 | CAGCGTGGACCGCTTGCTGCAACTCTC TCA | T-DNA Left-border primer (for *csn5a-2*)<br>(Dohmann *et al*., 2005) |
